# A conserved ubiquitin signalling module controls membrane raft integrity at the neuronal synapse

**DOI:** 10.64898/2026.11.23.768487

**Authors:** T.M. Carusone, V. Kedare, W. Marcantonio, S. Gharbi, N. Aranda, M. Rettel, M. Pelosse, A. de Kerchove d’Exaerde, S Bhogaraju

## Abstract

Activity-dependent ubiquitination of synaptic proteins is one of the key contributors of synaptic plasticity, along with localized protein synthesis and ubiquitin-dependent protein turnover. Substantial changes in the protein composition occur in the postsynaptic density upon chronic activation or inhibition of primary neurons. Precisely how a large-scale regulation of ensembles of synaptic proteins occurs through the ubiquitin proteasome system remains unknown. MAGE-D1 is a highly conserved protein enriched in brain and its deletion in mice leads to a range of behavioral phenotypes including defects in learning, memory and a complete resistance to cocaine addiction. Here, we found that MAGE-D1 acts as a scaffold bridging the ubiquitin ligase Praja-1 and the deubiquitinase Usp7. This MAGE-D1 signalling module directly interacts with and regulates the levels of Brain acid-soluble protein-1 (BASP1) which is a key component and stabilizer of neuronal membrane rafts. Disruption of MAGE-D1 complex function or chemical inhibition of Usp7 in mice results in resistance to cocaine-induced locomotor sensitization accompanied by specific degradation of oligomeric BASP1 and other key synaptic proteins in the postsynaptic density of mice thalamus. As a result, loss of MAGE-D1 in differentiated neurons leads to a dramatic disorganization of neuronal membrane rafts and anomalies in the synaptic membrane protein landscape. Finally, we mapped the structural architecture of the human MAGE-D1/Praja-1/Usp7 complex revealing the basis for the scaffolding nature of MAGE-D1. In summary, we discovered a conserved synaptic ubiquitin signalling network mediated by a protein complex that localizes to the synapse and regulates structures of membrane rafts and actin cytoskeleton.

## Introduction

The covalent attachment of ubiquitin (Ub) to target proteins is mediated through a three-step enzymatic cascade involving Ub-activating (E1), Ub-conjugating (E2) and Ub-ligating (E3) enzymes. While the majority of ubiquitinated proteins are degraded by the Proteasome, non-degradative ubiquitin signatures on a protein substrate are also prevalent and function in diverse processes, including signal transduction, membrane trafficking and DNA repair. Compared to its role in other areas of biology, the role of ubiquitin in neuronal synaptic signalling remains relatively under studied. However, seminal studies over the past several decades provided growing evidence linking ubiquitin proteasome system (UPS) to synapse maintenance, organization, and dysregulation during neurological diseases (*1*). Synaptic plasticity is at the cellular basis of learning and memory and is fine-tuned by a dynamic equilibrium between the synthesis and removal of synaptic proteins which is mediated through the UPS (*2*). Foundational studies linked the UPS to synaptic plasticity across species. In Aplysia, long-term facilitation of gill withdrawal reflux was associated with ubiquitin-dependent proteolysis of a regulatory subunit of cAMP-dependent protein kinase (PKA) (*3*, *4*). In humans, mutations in the E3 ligase UBE3A causes Angelman syndrome, an autism spectrum disorder associated with severe cognitive impairment (*5*). In mice, loss of function of Ube3a leads to decreased dendritic branching and spine density accompanied with defects in long-term potentiation and context-dependent learning (*6*, *7*). Strikingly, stimulating the synaptic activity of primary rat cortical neuronal culture results in selective ubiquitination and UPS-mediated degradation of several post-synaptic density (PSD) scaffold proteins including Shank, GKAP and AKAP79/150 (*8*). Ubiquitination of Shank and GKAP has also been shown to occur *in vivo* in the mouse hippocampus upon retrieval of conditioned fear memory (*9*). In alignment with the activity-dependent changes in the protein composition, upon synaptic activation, proteasome complexes are sequestered from the dendritic shafts into the spines to perform localized degradation (*10*). Isolation of ubiquitinated proteins from rat cortical neurons and subsequent proteomics analysis revealed that a large number of synaptic proteins are ubiquitinated (*9*). The E3 ligases involved in each of these ubiquitination events and the contribution of these towards the plasticity of synapse remains an active area of investigation.

Human MAGE-D1 is a brain-enriched protein that shares ∼90% sequence identity with the mouse counterpart indicating that it performs a conserved neuronal function (*12*, *13*). MAGE-D1 belongs to the Melanoma associated antigens (MAGEs) family, which scaffold E3 ubiquitin ligases to mediate substrate-specific ubiquitination (*12*, *14*). Phenotypes arising from the loss of function of MAGE-D1 are widely studied using KO mice (*15–18*). MAGE-D1 KO mice do not exhibit any gross physical abnormalities and are born normally as per Mendelian inheritance but they show a range of behavioural phenotypes (*18*). MAGE-D1 KO mice exhibit social reclusiveness, late onset of obesity, impaired sexual behaviour described to be reminiscent of autism spectrum disorder and Prader-Willi syndrome (*17*). Hippocampal neurons from the MAGE-D1 KO mice exhibit reduced dendritic spine density associated with defects in learning and memory in the KO animals (*15*). In our recent study, we demonstrated that conditional KO of MAGE-D1 in the paraventricular thalamus (PVT) region of the mice brain is necessary and sufficient to abolish cocaine addictive behaviours (*19*). Finally, loss of function or de novo variants of MAGE-D1 has been linked with intellectual disability and failure to thrive in humans (*20–22*). These observations establish a role for MAGE-D1 in synaptic activity associated especially with learning, memory and addiction.

Yeast-two hybrid studies showed that MAGE-D1 interacts with the E3 ubiquitin ligases Praja-1 and Praja-2 through its MAGE-homology domain (MHD) (*23*). Using endogenous immunoprecipitation (IP) of MAGE-D1 from neuroblastoma cell line SH-SY5Y and mouse thalamus, we further demonstrated that MAGE-D1 interacts with the deubiquitinase Usp7 as well as with the E3 ligases Praja-1/Praja-2 (*19*). In the mouse PVT, we also found that MAGE-D1 regulates cocaine-induced transcriptional responses through modulation of histone H2A ubiquitination (*19*). However, this ubiquitination event and the potential downstream transcriptional regulation do not explain the acute abrogation of cocaine-induced locomotor sensitization observed in MAGE-D1 KO mice (*16*). Given the synaptic modulation-linked behavioural anomalies showcased by the MAGE-D1 KO mice and its direct interactions with the components of the ubiquitin system, we hypothesized that MAGE-D1 complex regulates synaptic components through ubiquitin-mediated processes. So far, no synaptic protein has been shown to directly associate or get ubiquitinated via MAGE-D1 complexes.

Here, using differentiated human SH-SY5Y neuroblastoma cells, we identified brain acid–soluble protein 1 (BASP1) as a neuronal substrate of the MAGE-D1 complex. We show that BASP1 levels are also regulated through MAGE-D1 *in vivo* in the mice thalamus. BASP1 is ubiquitinated and degraded by the E3 ligase Praja-1 while it is stabilized through the deubiquitinase Usp7, two opposing actions under the control of MAGE-D1. MAGE-D1 is required for the proper localization of BASP1 to membrane rafts in both human neuroblastoma cells and in mouse thalamus. In differentiated SH-SY5Y cells, loss of MAGE-D1 disrupts membrane raft organization, alters membrane protein composition and a network of actin-binding proteins essential for synaptic structural integrity. We also dissected the molecular underpinnings of MAGE-D1/Praja/Usp7 interactions through a crystal structure and AlphaFold modelling. Together, these findings define MAGE-D1 as a signalling scaffold protein regulating synaptic protein organization and indicate that ubiquitin-dependent control of synaptic substrates underlies the behavioural phenotypes associated with MAGE-D1 deficiency.

## Results

### MAGE-D1/Usp7 complex stabilizes BASP1 at the neuronal synapse

We hypothesized that the proteins regulated through ubiquitination or deubiquitination by the MAGE-D1/Praja1/Usp7 complex might be depleted or enriched respectively in a MAGE-D1 dependent manner. To identify such candidate proteins, we used quantitative proteomics to compare the total proteomes of the differentiated wildtype (WT) and MAGE-D1 KO SH-SY5Y cells (Figure 1A). SH-SY5Y cells were differentiated for seven days with retinoic acid (RA) into a dopaminergic-like phenotype (*24–26*), characterized by increased expression of neuronal markers and synaptic proteins (*26*, *27*) but lack functional synapses. Several proteins are found to be differentially regulated between WT and MAGE-D1 KO SH-SY5Y cells (Figure 1A and Supplementary Table 1). Neuronal cell adhesion molecule (NRCAM), the neuronal actin binding protein (Neurabin-1), the postsynaptic density protein (PSD-93) and the neuronal membrane raft protein Brain acid-soluble protein 1 (BASP1) are among the significantly upregulated proteins in WT cells. Interestingly, a high-affinity cocaine-binding site was recently found to be associated with BASP1 in mice (*28*). Given that MAGE-D1 KO mice exhibit resistance to cocaine-addiction (*19*), we probed if there is any direct association between MAGE-D1 and BASP1. In our previous IP-MS experiments performed in SH-SY5Y cells (*19*), BASP1 was not found as an interactor of MAGE-D1. We reasoned that this is because BASP1 is known to be enriched in the detergent-resistant membrane (DRM) fraction (*29*), which is Triton X-100 insoluble and is sensitive to cell scraping. We modified the cell lysis procedure (see methods) and performed IP of endogenous MAGE-D1 from SH-SY5Y cells followed by the analysis of the interactome using quantitative proteomics (Figure 1B, Supplementary Table 2). We identified BASP1 as the most significant interactor of MAGE-D1 in these conditions. In addition, we also found cell adhesion molecule ALCAM and Dopamine beta-hydroxylase (DBH), which catalyzes the conversion of dopamine (DA) to norepinephrine (NE). BASP1 was also identified as a significant interactor of endogenous MAGE-D1 in HEK cells (Supplementary Figure 1A, Supplementary Table 3). Given the direct interaction between MAGE-D1 and the deubiquitinase Usp7 (*19*), we next examined whether Usp7 contributes to the stability of BASP1. BASP1 exhibits unusually slow electrophoretic mobility in SDS-PAGE owing to its myristoylation-dependent oligomerization that is resistant to denaturing sodium dodecyl sulphate (SDS) (*30–33*). It is possible to distinguish a monomeric species migrating at approximately 55 kDa and a higher-order oligomeric assembly migrating at approximately 200-275 kDa corresponding to the functional membrane-associated form of BASP1 (*32*, *33*). Treatment of differentiated SH-SY5Y cells with the Usp7 selective inhibitor HBX 41108 led to a dramatic decrease in SDS-resistant oligomeric BASP1 levels and to a partial reduction of MAGE-D1 in WT cells, as analyzed by Western Blot (WB) (Figure 1C). Importantly, as expected from Figure 1A, MAGE-D1-KO cells exhibited reduced BASP1 levels compared with WT. Treatment of MAGE-D1-KO cells with Usp7 inhibitor did not change BASP1 levels, indicating that Usp7 stabilizes BASP1 in a MAGE-D1 dependent manner (Figure 1C). We also observed that BASP1 in MAGE-D1 KO cells has slightly higher electrophoretic mobility in comparison to WT cells suggesting an alteration of its oligomeric assembly (*33*). Intriguingly, the levels of monomeric BASP1 remain unaffected by the absence of MAGE-D1 or the presence of Usp7 inhibitor indicating that MAGE-D1 could be selectively regulating the oligomeric form of BASP1 at least in the conditions tested. We also found treating differentiated SH-SY5Y cells with the proteasome inhibitor MG132 in addition to the Usp7 inhibitor treatment partially rescued BASP1 levels supporting a model in which Usp7 counteracts proteasome-dependent degradation of BASP1 (Supplementary Figure 1B). To assess whether MAGE-D1/Usp7 regulates BASP1 *in vivo*, we inhibited Usp7 activity in mice with P5091, a selective Usp7 inhibitor validated in our previous study for *in vivo* administration (*19*). These mice were tested for the expected Usp7-inhibitor induced reduction in cocaine-induced locomotor sensitization before the sacrifice and the thalamus isolation (Supplementary Figure 1C-D, Supplementary Table 4) (*19*). Usp7 inhibition reduced the oligomeric form of BASP1 also in the mice thalamus, consistent with our observations in human neuroblastoma cells (Figure 1D). We also observed that, in mouse brain, BASP1 is detected predominantly in its oligomeric form. We next isolated synaptosomes (Syn.) and post-synaptic density (PSD) fractions from the mice thalamus and probed whether MAGE-D1/Usp7 mediated regulation of BASP1 occurs at the synapse (Figure 1E). Firstly, we found that BASP1 and MAGE-D1 are enriched in the PSD fractions and secondly, this data revealed that the levels of BASP1 are reduced in response to the Usp7 inhibition directly in the PSD. Given that the oligomeric form of BASP1 corresponds to its membrane-associated state, we next probed the effect of MAGE-D1 on the membrane localization of BASP1 in differentiated SH-SY5Y cells. For this, we performed Stochastic Optical Reconstruction Microscopy (STORM) which enabled spatial resolution of BASP1 localization at the nanoscale. In the WT cells, we observed a specific BASP1 membrane localization enriched at the cell-cell junctions, while in the MAGE-D1 KO cells BASP1 membrane localization is significantly reduced and it mostly appears dispersed in the cytoplasm (Figure 1F-G, Supplementary Figure 1E). It is also worth noting that MAGE-D1 KO cells displayed a distinct morphology, appearing flatter and larger than WT cells, likely as a result of cytoskeletal defects (Supplementary Figure 1E, see below). These results indicate that MAGE-D1-dependent stabilization of the oligomeric BASP1 is required for its proper membrane localization. Taken together these findings suggest that MAGE-D1 together with Usp7 regulates the abundance of membrane-associated BASP1 by preventing its degradative ubiquitination at the postsynaptic density.

**Figure 1.**
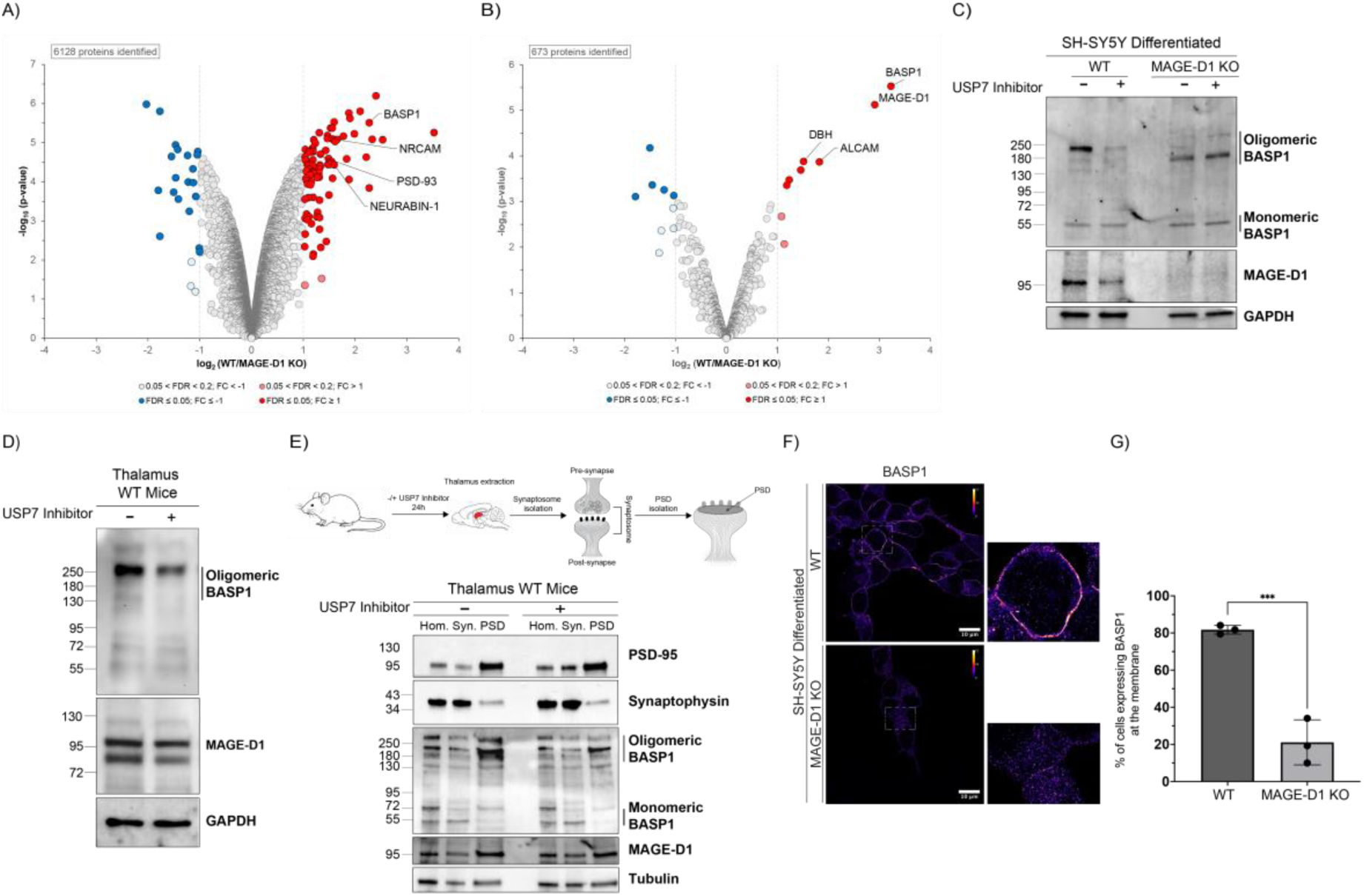
MAGE-D1 and Usp7 are required for BASP1 stability and membrane localization. A) Volcano plot of total protein levels isolated from WT and MAGE-D1 KO differentiated SH-SY5Y cells. Results are representative of three independent biological replicates. Significantly altered proteins are shown in dark red or blue (FDR ≤ 0.05, log2FC ≥I1I) and light red or blue (0.2 <FDR <0.05, log2FC > I1I) (moderated t-test). B) Volcano plot of enriched proteins isolated after endogenous immunoprecipitation of MAGE-D1 from WT and MAGE-D1 KO undifferentiated SH-SY5Y cells. Results are representative of three independent biological replicates. Significantly altered proteins are shown in dark red or blue (FDR ≤ 0.05, log2FC ≥I1I) and light red or blue (0.2 <FDR <0.05, log2FC > I1I) (moderated t-test). C) Immunoblotting of BASP1, MAGE-D1 and GAPDH (loading control) from total lysate of differentiated SH-SY5Y WT and MAGE-D1 KO cells under vehicle control condition (DMSO) or Usp7 Inhibition (HBX 41108, 10uM) for 6 hours. Oligomeric and monomeric BASP1 forms are indicated. Results are representative of five independent biological replicates. D) Immunoblotting of BASP1, MAGE-D1 and GAPDH (loading control) from total lysate of mice thalamus under vehicle control condition (5% DMSO + 20% SBE-β-CD in saline) or Usp7 Inhibition (P5091, 10 mg/kg; intraperitoneal injection) for 7 days. Oligomeric form of BASP1 is indicated. E) Schematic representation of the protocol used for PSD isolation from the thalamus of mice brain (top panel). Immunoblotting of PSD-95, Synaptophysin, BASP1, MAGE-D1 and Tubulin (loading control) from total homogenate (HOM.), synaptosome (SYN.) and post-synaptic density (PSD) fractions isolated from thalamus of mice under vehicle control condition (5% DMSO + 20% SBE-β-CD in saline) or Usp7 Inhibition (P5091, 10 mg/kg; intraperitoneal injection) for 7 days. Oligomeric and monomeric BASP1 forms are indicated (bottom panel). Results are representative of three independent biological replicates. F) STORM images of differentiated SH-SY5Y WT and MAGE-D1 KO cells immunostained with anti-BASP1 antibody. Images are rendered using the Fire LUT; the accompanying color scale represents relative localization density, with brighter colors indicating regions containing a greater number of detected BASP1 localizations. Scale bar 10 µm. G) Percentage of cells displaying plasma membrane localization of BASP1 in WT and MAGE-D1 KO differentiated SH-SY5Y cells calculated from confocal microscopy images (Supplementary Figure 2A). Data are shown as mean ± SD from three independent biological replicates (total cells analyzed: WT, n=311; MAGE-D1 KO, n=228). Statistical significance was determined using an unpaired t-test (*p* =0.0001, ***).

### MAGE-D1 complex is essential for BASP1 membrane localization and neuronal membrane raft organization

Since the emergence of the “lipid rafts” concept (*34*), their presence and form has been brought into question (*35*, *36*). But it is generally accepted that membrane micro domains of 10-200 nm form into so called “membrane rafts” that act in part to cluster signalling proteins (*37*, *38*). These membrane rafts are typically enriched in cholesterol and sphingolipids and are thought to be stabilized by membrane-interacting oligomerizing proteins (*39*). A string of previous studies which involved cellular and biophysical characterization of BASP1 indicate that it is a membrane-raft stabilizing protein. Notably, NAP-22, a homologue of BASP1 in rats is one of the most abundant protein in the rat brain accounting for 0.4-0.8% of the total brain protein (*40*). NAP-22 is also one of the most-abundant proteins in the so called “detergent-resistant membrane” fraction or membrane rafts of the rat brain (*29*, *40*). Through non-specific ionic interactions mediated through its positively charged residues, BASP1 co-clusters with negatively charged lipids such as phosphatidylinositol (4,5) bisphosphate (PIP2) both *in vitro* and in cells (*41*, *42*). Myristoylation of BASP1 is also known to trigger oligomerization and membrane binding of the protein *in vitro* (*32*, *33*). BASP1 also interacts with Calmodulin (CaM) in a Ca^2+^-dependent manner and binding to CaM disrupts oligomer formation and membrane interaction (*32*, *40*). Taken together, these observations strongly suggest that BASP1 is an important architect of membrane organization in neuronal cells. Since we observed a reduction of BASP1 membrane localization in MAGE-D1 KO cells (Figure 1F-G), we next examined whether MAGE-D1 similarly regulates BASP1 membrane association *in vivo*. Due to the limited amount of thalamic tissue available, membrane rafts were isolated from total brains of WT and MAGE-D1 KO mice (Supplementary Figure 2A). As proof of principle for the isolation of membrane rafts from crude lysates, we detected the oligomeric form of BASP1 co-fractionating with flotillin-1 (FLOT-1), a planar membrane raft marker (Supplementary Figure 2A). Consistent with our findings in differentiated neuroblastoma cells, loss of MAGE-D1 also dramatically reduced the total BASP1 levels found in the membrane raft fractions *in vivo* (Figure 2A). To test if loss of function of BASP1 triggered through the depletion of MAGE-D1 affects the membrane-raft integrity, we used two independent membrane raft markers: FLOT-1 a scaffolding protein that acts as a planar membrane raft marker and Cholera toxin B (CTB) which binds to the external ganglioside GM1. In MAGE-D1 KO cells, both markers displayed significantly reduced membrane localization, indicating compromised membrane raft organization (Figure 2B-C). Confocal microscopy of differentiated SH-SY5Y cells showed that the endogenous MAGE-D1 colocalized with the membrane raft marker cholera toxin B (CTB) (Figure 2D), supporting the presence of MAGE-D1 in these specialized membrane compartments. Membrane rafts are thought to be important for the clustering of membrane proteins mediating various signalling processes (*43*). In neurons, several AMPA receptors and NMDA receptors have been shown to be present in the detergent resistant membranes which are thought to represent membrane rafts (*44–46*). We reasoned that disorganization of membrane raft triggered through displacement of BASP1 in MAGE-D1 KO cells might lead to defects in the proper localization of membrane-raft localized proteins. To test this, differentiated SH-SY5Y cells were treated with a membrane-impermeable, amine-reactive, sulfo-NHS-SS-biotin to label cell surface proteins. Cells were then lysed, and biotinylated proteins were captured using NeutrAvidin agarose and analyzed by proteomics (Figure 2E, Supplementary Table 5). WB analysis indicated that while the membrane protein Transferrin receptor-1 got pulled down with NeutrAvidin agarose, the absence of GAPDH indicates efficient isolation of the membrane protein fraction (Supplementary Figure 2B). Comparative proteomics of membrane protein fraction of WT and the MAGE-D1 KO cells revealed that there are dozens of membrane proteins differentially expressed in a MAGE-D1 dependent manner (Figure 2E). Interestingly, BASP1 is depleted from the membrane fraction of MAGE-D1 KO cells as expected. Even though BASP1 is only known to be a membrane-anchored protein, it might have strong associations with transmembrane proteins that were labeled using sulfo-NHS-SS-biotin. Other notable synaptic membrane associated proteins depleted from the MAGE-D1 KO cells include the metabotropic Glutamate receptors mGluR7/mGluR8, adhesion molecules ELFN1/LRFN2/NRCAM/NRXN1 and DBH. Furthermore, Gene Ontology (GO) analysis of proteins enriched in WT cells showed a significant overrepresentation of synaptic membrane and postsynaptic density components, further supporting their depletion from the membrane fraction of MAGE-D1 KO cells (Figure 2F). Previous immunohistochemical localization showed that BASP1 homologue in rat brains is localized to both post and pre-synaptic membranes and also to synaptic vesicles (*47*). Our biochemical fractionation (Figure 1E) of the mice thalamus also showed Usp7-dependent regulation of BASP1 in the PSD of the mice thalamus. Taken together, it is likely that MAGE-D1-mediated regulation of BASP1 plays a critical role in the proper membrane-raft organization at the neuronal synapse. Based on our observations in the differentiated SH-SY5Y cells, MAGE-D1/BASP1 axis is also likely to play a key role in the homeostatic localization of membrane proteins at the neuronal synapse.

**Figure 2.**
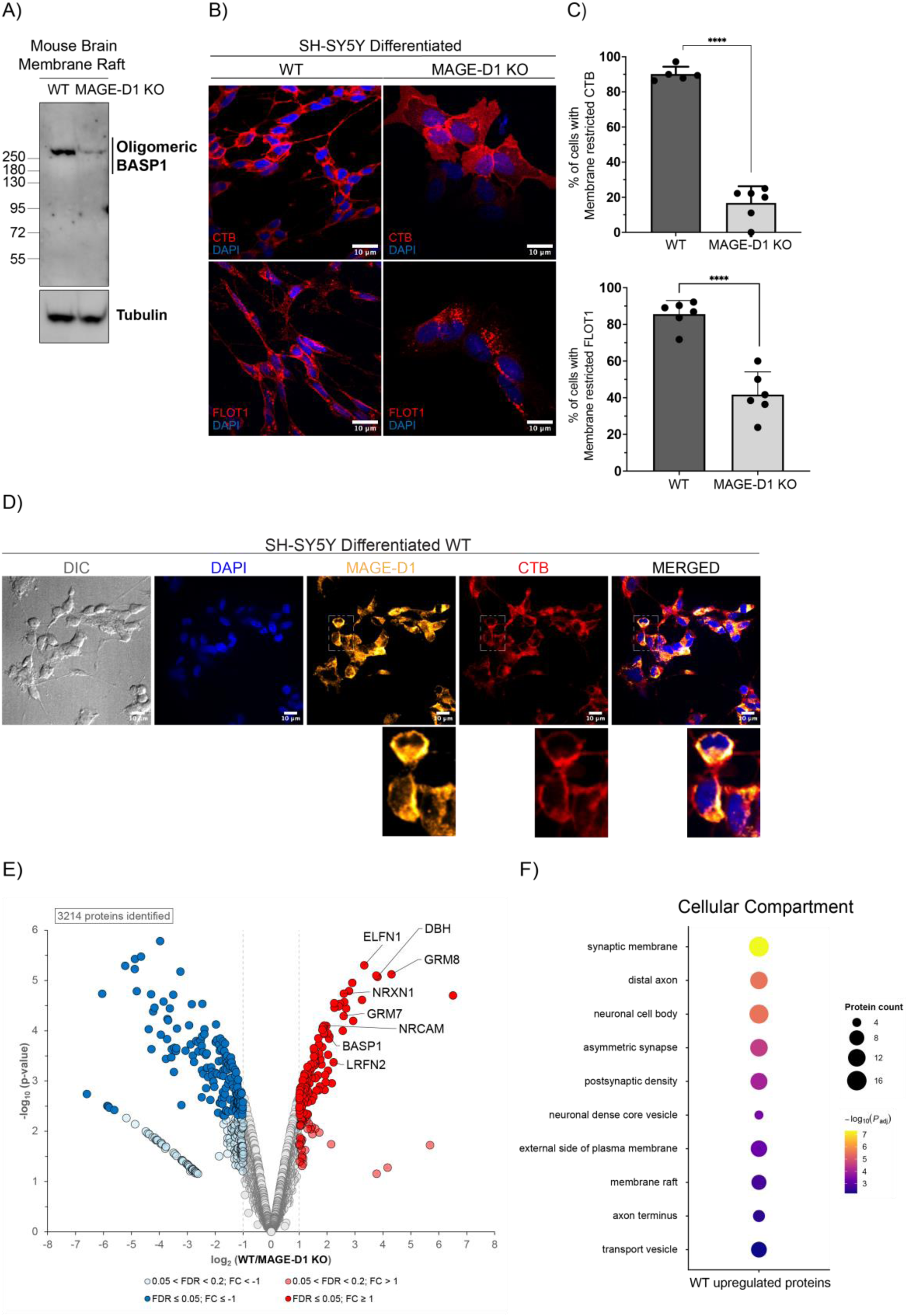
MAGE-D1 KO displaces BASP1 from membrane rafts and alters neuronal membrane raft organization. A) Immunoblotting of BASP1 and Tubulin (loading control) from membrane rafts isolated from thalamus of WT and MAGE-D1 KO mice. Oligomeric BASP1 form is indicated. B) Confocal microscopy images of differentiated SH-SY5Y WT and MAGE-D1 KO cells. Top panel: staining with CTB (membrane raft marker, red) and DAPI (nuclei, blue). Bottom panel: immunostaining with anti-FLOT1 antibody (membrane raft marker, red) and DAPI (nuclei, blue). Scale bar 10 µm. C) Percentage of cells displaying membrane restricted CTB localization (top panel) and FLOT1 (bottom panel) in WT and MAGE-D1 KO differentiated SH-SY5Y cells. For CTB, data are shown as mean ± SD from five fields of view for WT and seven fields of view for MAGE-D1 KO cells (total cells analyzed: WT, n=207; MAGE-D1 KO, n=43). Statistical significance was determined using an unpaired t-test (*p*=0.0001, ****). For FLOT1 data are shown as mean ± SD from three fields of view obtained from two biological replicates for condition (total cells analyzed: WT, n=317; MAGE-D1 KO, n=107). Statistical significance was determined using an unpaired t-test (*p*=0.0001, ****). D) Confocal microscopy images of differentiated SH-SY5Y WT and MAGE-D1 KO cells. Differential interference contrast (DIC) images are shown alongside fluorescence images of cells stained with DAPI (nuclei, blue), immunostained with anti-MAGE-D1 antibody (orange), and stained with CTB (red). Scale bar 10 µm. Colocalization between CTB and MAGE-D1 was assessed using Pearson’s correlation coefficient (PCC). The mean PCC was 0.52 ± 0.21 (mean ± SD, n=74 cells; 95% CI, 0.48–0.57). Additionally, 67% of analyzed cells exhibited PCC values > 0.5. E) Volcano plot of cell surface proteins isolated from WT and MAGE-D1 KO differentiated SH-SY5Y cells. Results are representative of three independent biological replicates. Significantly altered proteins are shown in dark red or blue (FDR ≤ 0.05, log2FC ≥I1I) and light red or blue (0.2 <FDR <0.05, log2FC > I1I) (moderated t-test). F) Gene Ontology (GO) cellular component enrichment analysis of WT-upregulated proteins. Bubble plot showing significantly enriched Gene Ontology (GO) Cellular Compartment (CC) terms among proteins upregulated in the WT group. The y-axis lists the top enriched cellular compartments. Bubble size represents the number of proteins associated with each GO term, while bubble color indicates the enrichment significance expressed as −log10(adjusted P value), with warmer colors representing higher statistical significance.

### MAGE-D1/Usp7 complex co-regulates several synaptic cytoskeletal proteins

A previous study has found that PSD scaffold proteins Shank, GKAP and AKAP79/150 are ubiquitinated in response to neuronal activity in the mouse cortical neurons while other synaptic proteins such as NR2B, SAP102, PP1 and PKA-RIIβ were also co-regulated by the UPS despite no detectable ubiquitination of these proteins (*8*). It was suggested that UPS regulates PSD in units of “ensembles” of closely interacting proteins in response to the chronic activation or inhibition of synapse (*48*). Based on this, we hypothesized that MAGE-D1 complexes might be regulating more synaptic proteins in addition to BASP1. To test this, we searched through the total proteome and interactome of MAGE-D1 from differentiated SH-SY5Y (Figure 1A) and from the mice thalamus that we previously published (Supplementary Figure 3, Supplementary Table 6) (*19*). We consistently found that several neuronal cytoskeletal proteins including Neurabin-1, Neurabin-2, Drebrin-1 and the post-synaptic scaffold protein PSD-95 were among the notable interactors of MAGE-D1. Interestingly, similar to BASP1, Neurabin-1 and PSD-93, which is a member of PSD-95 family essential for PSD structure, were also found to be differentially regulated at the whole proteome level in a MAGE-D1 dependent manner (Figure 1A). Based on these observations, we probed for the levels of Neurabin-1, Neurabin-2 and DBN-1 in the differentiated SH-SY5Y WT and MAGE-D1 KO cells. We observed that all these neuronal cytoskeletal regulating proteins were down regulated in MAGE-D1 KO cells (Figure 3A). To test if this regulation is Usp7 dependent, we inhibited Usp7 in both differentiated SH-SY5Y and in the mice thalamus. Both in our cellular model and in the mouse thalamus, immunoblotting revealed that Usp7 inhibition leads to the down regulation of Neurabins (Figure 3B-C). In addition to the cytoskeletal proteins, we were also intrigued by the reported interaction between BASP1 and the planar membrane-raft markers Flotillin-1/Flotillin-2 that is consistently detected in several different large-scale interactome studies (*49*, *50*). Similar to the cytoskeletal proteins that we probed, levels of FLOT-1 were also regulated in a Usp7 activity-dependent manner both in the differentiated SH-SY5Y cells and in the mice thalamus (Figure 3D-E). Neurabin-1 and 2 are actin-binding proteins that also contain a PDZ domain interacting with the plasma membrane (*51*, *52*). Neurabins have been shown to stabilize cortical actin/membrane interactions and play an important role in maturation of neuronal filopodia into dendritic spines (*52*). DBN-1 is another postsynaptic protein that possesses multiple actin-binding domains and is responsible for the stability of actin filaments (*53*). Importantly, BASP1 deleted cultures of mouse dorsal root ganglion (DRG) cells exhibited thin and highly twisted neurite outgrowth which is remarkably similar to what WT DRG neurons exhibited when treated with cytochalasin D, an inhibitor of actin polymerization (*54*). Cortical actin network and membrane rafts are known to be intricately linked together and this interaction plays a critical role for the clustering of raft localized membrane proteins (*55–57*).

**Figure 3.**
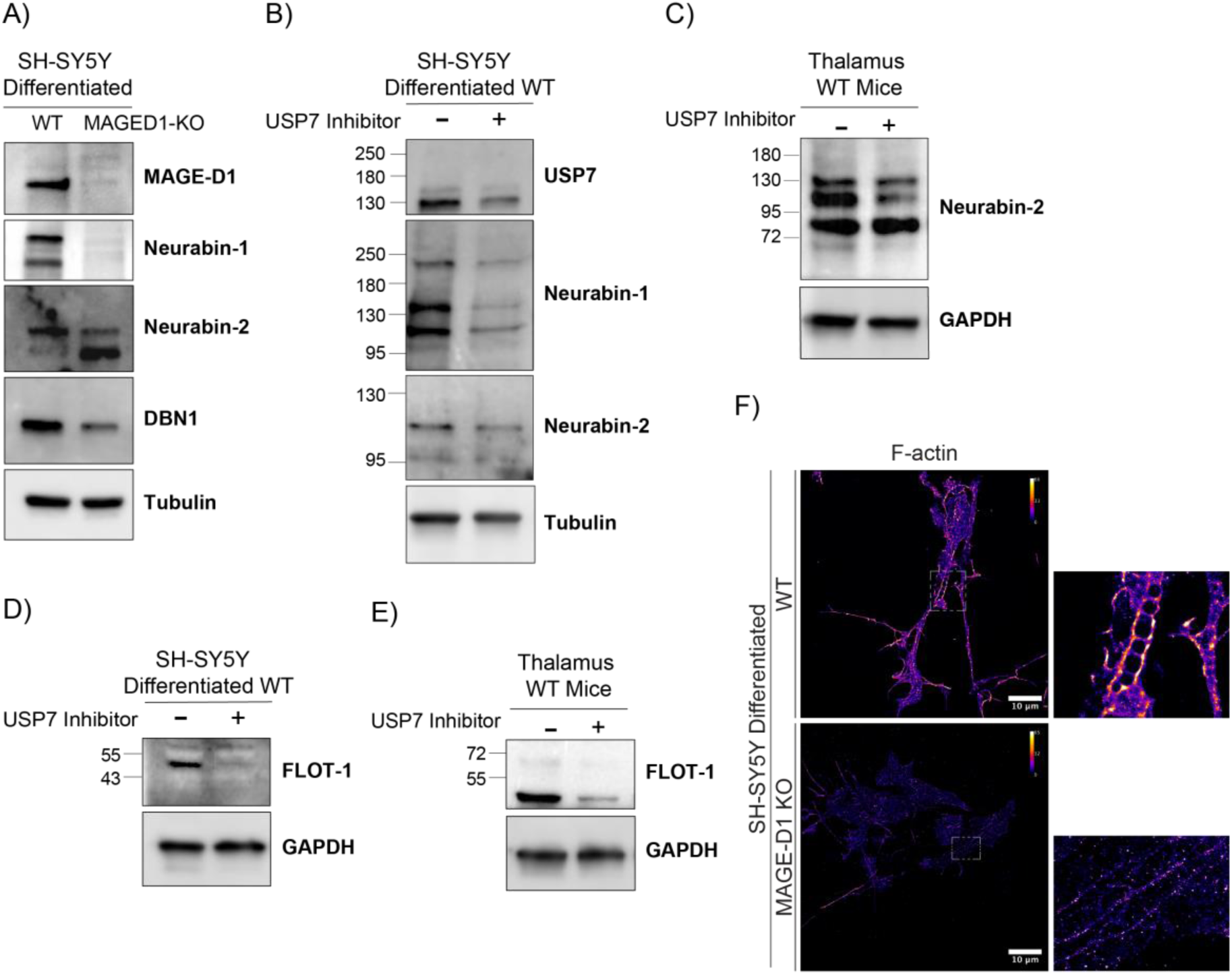
MAGE-D1/Usp7 complex stabilizes synaptic actin-regulatory proteins and preserves cortical F-actin organization. A) Immunoblotting of MAGE-D1, Neurabin-1, Neurabin-2, DBN1 and Tubulin (loading control) from total lysate of differentiated SH-SY5Y WT and MAGE-D1 KO cells. B) Immunoblotting of Usp7, Neurabin-1, Neurabin-2 and Tubulin (loading control) from total lysate of differentiated SH-SY5Y WT cells under vehicle control condition (DMSO) or Usp7 inhibition (HBX 41108, 10uM) for 6 hours. C) Immunoblotting of Neurabin-2 and GAPDH (loading control) from total lysate of mice thalamus under vehicle control condition (5% DMSO + 20% SBE-β-CD in saline) or Usp7 Inhibition (P5091, 10 mg/kg; intraperitoneal injection) for 7 days. D) Immunoblotting of FLOT-1 and GAPDH (loading control) from total lysate of differentiated SH-SY5Y WT cells under vehicle control condition (DMSO) or Usp7 inhibition (HBX 41108, 10uM) for 6 hours. E) Immunoblotting of FLOT-1 and GAPDH (loading control) from total lysate of mice thalamus under vehicle control condition (5% DMSO + 20% SBE-β-CD in saline) or Usp7 Inhibition (P5091, 10 mg/kg; intraperitoneal injection) for 7 days F) STORM images of F-actin stained with Phalloidin in differentiated SH-SY5Y WT and MAGE-D1 KO cells. Images are rendered using the Fire LUT; the accompanying color scale represents relative localization density, with brighter colors indicating regions containing a greater number of detected F-actin localizations. Scale bar 10 µm.

Given the central role of these actin-binding proteins in organizing the cortical actin cytoskeleton and maintaining postsynaptic structure and synaptic transmission, we next examined F-actin organization using phalloidin staining and STORM imaging in differentiated SH-SY5Y cells. In WT cells, actin filaments appeared well polymerized and organized into linear structures. In contrast, MAGE-D1 KO cells displayed a disrupted filament network, with reduced filament continuity and decreased overall phalloidin fluorescence intensity, consistent with impaired actin polymerization (Figure 3F). Our data indicate MAGE-D1/Usp7 mediated ubiquitin signalling regulates an “ensemble” of proteins responsible for F-actin and membrane-raft organization at the neuronal synapse.

### MAGE-D1 scaffolds opposing ubiquitination and deubiquitination activities

MAGE-D1 forms a hetero-trimeric complex with the E3 ubiquitin ligases Praja-1 and Praja-2 (28) and the deubiquitinase Usp7 (17). To define how opposing ubiquitination and deubiquitination activities are coordinated within this complex, we sought to map the interacting regions among Praja-1/2, MAGE-D1, and Usp7. MAGE-D1 comprises a C-terminal MAGE homology domain (MHD, residues 465-703), a conserved domain in MAGE proteins which is involved in both the Ub ligase interaction and substrate recognition (*58*, *59*). MAGE-D1 also contains a large unstructured region at the N-terminus (UNS, residues 1-465).

To map interactions to these individual regions of MAGE-D1, we transiently expressed GFP-MHD and GFP-UNS in HEK cells and performed quantitative proteomics of the GFP-IP fraction (Figure 4A, Supplementary Table 7). As expected, the overexpressed MAGE-D1 MHD interacted with the endogenous Ub ligases Praja-1 and Praja-2 from the HEK cells. Interestingly, BASP1 was also detected as interactors of MAGE-D1 MHD albeit with a lower Log_2_FC compared to the Praja Ub ligases. In contrast, GFP-MAGE-D1 UNS pulled down the endogenous Usp7 from HEK cells. Using AlphaFold, we first mapped the interaction of a poly basic region (PBR) in MAGE-D1 (residues 241-255) residing in the UNS region of MAGE-D1 to a region encompassing Ubl1/Ubl2 of the Tandem Ub-like domain (TUD) of Usp7 (Figure 4B). MAGE-D1 PBR contains a stretch of positively charged residues that form electrostatic interactions with the surface exposed negatively charged residues in Usp7 TUD in a manner that is highly reminiscent of Usp7 interactions with other poly basic region containing substrates RNF169, ICP0, UHRF1 and DNMT1 (Supplementary Figure 4A). Using isothermal titration calorimetry (ITC), we estimated the dissociation constant (Kd) of the binding between MAGE-D1 PBR and the TUD of the Usp7 to be 0.8 µmoles/L (Figure 4C). Next, we sought to map the regions in Praja Ub ligases that interact with MAGE-D1 MHD. AlphaFold predicted structure (*60*) of Praja-1 indicates that except the C-terminal RING domain (residues 565 to 643) most of the protein is unstructured (Supplementary Figure 4B). Intriguingly, adjacent to the C-terminal RING domain, three helices (from now on referred as 3H, residues 511-564) were predicted by AlphaFold although with relatively poor pLDDT scores compared to the RING domain (Supplementary Figure 4B). Based on our recent study (*59*, *61*) where we identified that a lone helix in the Ub ligase RAD18 packs into a ligase-binding cleft in the WH-B subdomain of MAGE-A4 MHD, we reasoned that Praja-1 3H region might interact with MAGE-D1 MHD. To test this, we prepared different deletion constructs of Praja-1 and by co-immunoprecipitation (co-IP) with GFP-MHD in HEK we mapped the interaction of MAGE-D1 MHD to a Praja-1 helical region spanning residues 501–531 (from now on referred to as MAGE-D1 interacting region (MIR)) (Supplementary Figure 4C). To obtain structural insights into the recognition of Praja1/2 Ub ligases through MAGE-D1 MHD, we sought to crystallize MHD in complex with the Praja-1 MIR. Since MAGE-D1 MHD alone was not soluble when expressed in *E. coli*, we fused Praja-1 minimal MIR region (residues 501-520) to MHD which aided in its purification and crystallization (Supplementary Figure 4D). We determined the crystal structure of this fusion construct at 1.9 Å resolution (Figure 4D). In agreement with the published MHD structures of other MAGE proteins (*58*, *61–63*), MAGE-D1 MHD is also composed of two winged helix subdomains: WH-A and WH-B. A part of Praja-1 MIR containing residues 505 to 517 adopts a largely extended conformation and packs into a cleft which is formed by helices α4, α5 and α8 in the WH-B of MAGE-D1. We previously mapped the binding region of Ub ligase RAD18 to a similar cleft in the WH-B of MAGE-A4, which can be termed as the Ligase Binding Cleft (LBC) (*59*). Comparison of the LBCs of MAGE-A4 and MAGE-D1 reveals that Praja-1 also binds to equivalent LBC in MAGE-D1 (Supplementary Figure 4E). However, unlike the extended conformation adopted by the Praja-1 MIR, RAD18 peptide forms one long helix that binds to the LBC of MAGE-A4. Inspecting the interacting residues between MAGE-D1 and Praja-1 reveals extensive hydrophobic contacts including residues V654, F650, V565, V569, M572 and W662 from the MHD of MAGE-D1 and V509, W511, F514 and F517 from Praja-1 MIR (Figure 4E). Substitution of a single hydrophobic residue V569 in the MHD of MAGE-D1 with a charged Aspartic acid results in loss of binding to both Praja-1 and Praja-2 Ub ligases in cells (Figure 4F, Supplementary Table 8). Conversely, substitution of F514 and F517 with charged aspartic acid in Praja-1 and substitution of corresponding F553 and F556 in Praja-2 leads to loss of binding to MAGE-D1 (Supplementary Figure 4F-G, Supplementary Table 9-10). Since Praja-1 and Praja-2 are identical in the MIR region, they are expected to bind MAGE-D1 in a similar manner and therefore would compete for the same LBC of MAGE-D1 MHD. Interestingly, disruption of the MAGE-D1/Praja-1 interaction also abolished Praja-1 association with Usp7 (Supplementary Figure 4F), suggesting that MAGE-D1 acts as a scaffold linking the Ub ligase Praja-1 and the deubiquitinase Usp7. Having mapped the MAGE-D1/Praja1-2/Usp7 interaction interfaces, we next used the minimal interacting domains of MAGE-D1 and Praja-1 to investigate whether this complex can directly ubiquitinate BASP1 *in vitro*. We focused on Praja-1 because it is known to be highly expressed in the brain and has emerged as an important regulator of neuronal proteostasis (*64–66*), with growing evidence linking it to synaptic function and plasticity (*67*, *68*). To test this, we performed an *in vitro* ubiquitination assay using purified BASP1 and a minimal complex of Praja1 RING and MAGE-D1 MHD proteins (Supplementary Figure 4H). Immunoblot analysis showed that BASP1 gets robustly ubiquitinated in the presence of the complex compared with the control condition (Figure 4G). Mass spectrometry analysis of the ubiquitinated BASP1 bands identified the modified lysine residues, several of which have previously been reported in ubiquitination databases (*69*) (Supplementary Figure 4I, Supplementary Table 11). Mass spectrometry also revealed the presence of K48, K11 in the reaction mix (Supplementary table 11) indicating that Praja-1-mediated ubiquitination is degradative consistent with our data in SH-SY5Y cells and in mice.

**Figure 4.**
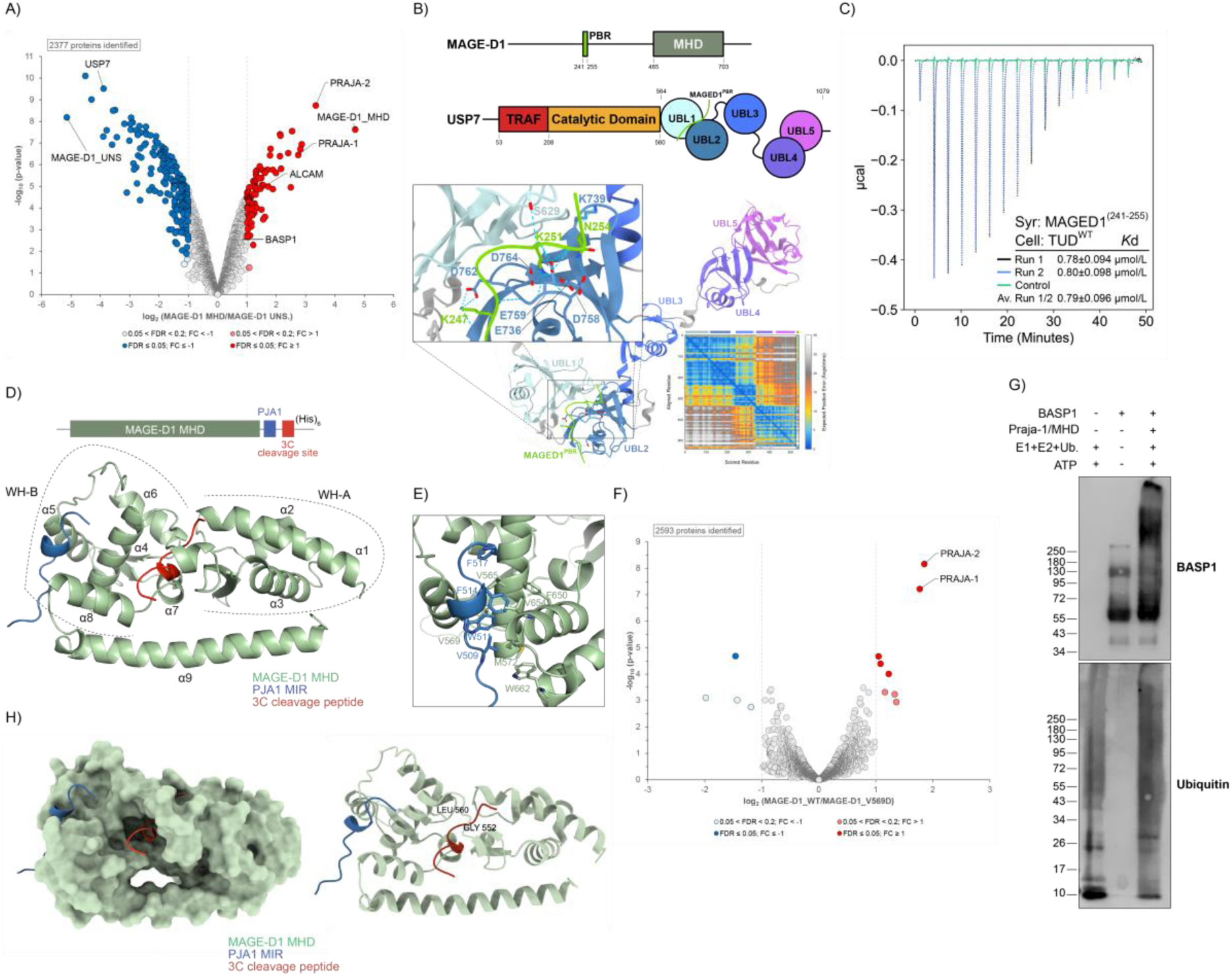
MAGE-D1 interacts with and coordinates Praja-1 and Usp7 to balance BASP1 ubiquitination. A) Volcano plot of total protein levels isolated after GFP immunoprecipitation of GFP_MAGE-D1_MHD and GFP_MAGE-D1_UNS over-expressed in HEK cells. Results are representative of three independent biological replicates. Significantly altered proteins are shown in dark red or blue (FDR ≤ 0.05, log2FC ≥I1I) and light red or blue (0.2 <FDR <0.05, log2FC > I1I) (moderated t-test). B) Schematic representations of MAGE-D1 and Usp7 domains. AlphaFold3 prediction of MAGE-D1 PBR and Usp7 TUD interaction with PAE plot (ipTM=0.76 pTM=0.68). Predicted binding region is enlarged, hydrogen bonding residues are indicated. C) ITC titration curve of MAGE-D1 PBR peptide (syringe) against Usp7 TUD (cell). Values from control and duplicates are indicated in the graph. D) Crystal structure of the fusion construct resolved at 1.9 Å resolution. Schematic representation of MHD/Praja-1 fusion construct composed of the MHD of MAGE-D1 (green), the Praja-1 MIR (blue) and a 3C cleavage site (red) before the His tag (top panel). E) Inset of the binding interface between the MHD and Praja-1 MIR. Hydrophobic contacts including residues V654, F650, V565, V569, M572 and W662 from the MHD of MAGE-D1 and V509, W511, F514 and F517 from Praja-1 MIR are indicated. F) Volcano plot of total protein levels isolated after GFP immunoprecipitation of GFP_MAGE-D1_WT and GFP_MAGE-D1_V569D mutant over-expressed in HEK cells. Results are representative of three independent biological replicates. Significantly altered proteins are shown in dark red or blue (FDR ≤ 0.05, log2FC ≥I1I) and light red or blue (0.2 <FDR <0.05, log2FC > I1I) (moderated t-test). G) *In vitro* ubiquitination assay of BASP1 performed with MAGE-D1 MHD in complex with the Praja-1 minimal RING construct. BASP1 migrated at an apparent molecular weight of 55 kDa. The electrophoretic mobility of BASP1 has been reported to vary depending on multiple factors, including SDS–PAGE conditions (*30*, *32*, *33*). Results are representative of three independent biological replicates. H) Structural representation of the substrate-binding cleft (SBC) of MAGE-D1 with the Praja-1 minimal MIR and the 3C cleavage site. G552 and L560 residues into the SBC involved in BASP1 binding are indicated (Supplementary Figure 4K).

We next performed a cell-based ubiquitination assay in HEK and found that BASP1 protein levels were reduced upon Praja-1, but not Praja-2, overexpression. The reduction of BASP1 is less pronounced in MAGE-D1 KO cells further supporting a specific role for the Praja-1/MAGE-D1 complex in the ubiquitin-dependent degradation of BASP1 (Supplementary Figure 4J). Since we found that BASP1 is ubiquitinated through Praja-1/MAGE-D1 complex and that BASP1 also binds to the MHD of MAGE-D1 (Figure 4A), we next sought to characterize the structural basis of this interaction. Inspection of the MHD/Praja-1 crystal structure revealed a surprising artefact in which a part of the 3C cleavage site of the fusion construct was found interacting with the MHD right at the intersection of WH-A and WH-B subdomains (Figure 4D). Interestingly, this region of MHD was annotated as substrate-binding cleft (SBC) and was found to be responsible for the binding of ubiquitination substrate in the case of MAGE-A11, MAGE-A3 and MAGE-F1 (*58*). The SBC of MAGE-D1 is buried deep into the intersection of WH-A and WH-B subdomains of MHD forming a large pocket and is unlike the surface-exposed LBC (Figure 4H). To test whether this pocket mediates BASP1 binding, we introduced two mutations into MHD, namely, G552N and L560D both of which were designed to obstruct any incoming substrate entering the SBC. Co-IP experiments with BASP1-HA and GFP-MHD WT or mutants in HEK cells showed that while WT MHD pulled down BASP1-HA, both SBC mutants failed to do so to the same extent indicating that the putative binding pocket we identified between WH-A and WH-B of MAGE-D1 MHD indeed forms the SBC and directly interacts with BASP1 (Supplementary Figure 4K).

Together, these findings define the structural basis of the MAGE-D1/Praja-1/2/Usp7 and MAGE-D1/BASP1 interactions. They also establish MAGE-D1 as a scaffold that coordinates Praja-1 E3 ligase and Usp7 deubiquitinase activities towards the ubiquitin-dependent regulation of their substrate BASP1.

## Discussion

Neuronal activity-driven synaptic plasticity requires in part rapid restructuring of synaptic protein composition which is achieved both by local protein synthesis and turnover. There are over 5000 different ubiquitination sites detected in the synaptosomes isolated from the whole brain samples of Wistar rats (*70*). Accordingly, several studies have found activity-dependent ubiquitination of specific synaptic proteins (*1*, *11*).

Here, we found that MAGE-D1 scaffolded ubiquitin signalling module is enriched specifically at the post-synaptic density and regulates a membrane anchored protein called BASP1. Human BASP1 is a 227 amino acid long protein that has unique physicochemical properties likely due to its highly unusual amino acid composition. Over 50% of amino acid residues in BASP1 are highly charged (Aspartates, Glutamates, Lysines, Arginines) and over 25% of the residues are Alanines. It is predicted to be completely disordered, however myristoylated BASP1 purified directly from the bovine brain has been shown to form highly structured oligomers which drive the clustering of negatively charged lipids both *in vitro* and in neurons (*29*, *32*, *33*, *41*, *42*). We have shown that MAGE-D1 complex specifically regulates the oligomeric BASP1 (Figure 1C, 1D), and accordingly, lack of MAGE-D1 function leads to absence of membrane raft bound BASP1 in differentiated human neuroblastoma cells and in the mice thalamus (Figures 2A, 2E). Crucially, we showed that BASP1 downregulation in turn triggers disruption of membrane raft integrity as evident from the diffuse staining of raft markers FLOT1 and CTB (Figure 2B) in differentiated SH-SY5Y cells. It is important to note that a notable fraction of glutamate receptors of both AMPA and NMDA kind are localized to postsynaptic membrane rafts (*71*). Additionally, AMPA receptors have been shown to undergo increased endocytosis upon chronic or acute disruption of neuronal membrane rafts indicating that membrane protein composition could be affected indirectly through the MAGE-D1/BASP1 axis-dependent mechanisms. Accordingly, quantitative proteomics revealed major changes in the membrane proteome of differentiated SH-SY5Y cells in a MAGE-D1 dependent manner (Figure 2E). GO analysis indicates that synaptic membrane proteins are specifically depleted from the MAGE-D1 KO cells (Figure 2F). It would be of high interest to map how the membrane protein composition is altered through the loss of function of MAGE-D1 complex directly at the neuronal synapse *in vivo*. Our results however strongly hint that MAGE-D1 KO mice phenotypes are likely a result of disruption of membrane raft integrity and the resulting anomalies in the membrane protein localization in the postsynaptic compartment. Such a potentially broad effect of MAGE-D1 complex at the postsynaptic compartment is indeed consistent with the pleotropic behavioral phenotypes observed in the MAGE-D1 KO mice. Further corroborating the strong functional interaction between MAGE-D1 and BASP1, a recent study reported that knock-down of BASP1 in mice reduces locomotor activity after cocaine administration (*28*), a phenotype similar to that observed in MAGE-D1 knockout mice (*16*, *19*). Although we find that MAGE-D1 is enriched at the post-synaptic density (PSD) (Figure 1E), its role at the pre-synaptic terminal cannot be ruled out given that BASP1 has been localized both in the pre- and post-synaptic regions (*47*).

Our previous work established that MAGED-1 and Usp7 regulate histone H2A ubiquitination in the nucleus (*19*). In the present study, we show that MAGED1 localizes to the neuronal plasma membrane (Figure 1E) and is enriched within membrane raft microdomains (Figure 2D). We further demonstrate that MAGE-D1/Usp7 complex together with Praja-1 and Praja-2, perform a distinct postsynaptic function, revealing that MAGE-D1 acts in both nuclear and synaptic compartments. These findings uncover an unexpected facet of MAGE-D1 biology and highlight its remarkable functional diversity in neurons.

Finally, we provide detailed structural insights into the MAGE-D1/Praja-1 interaction and define the precise interactions through which Praja-1, BASP1 and Usp7 associate with MAGE-D1. It is important to point that our interaction data does not exclude that MAGE-D1/Praja-1 and MAGE-D1/Usp7 exist as two independent complexes. However, in our HA-Praja-1 IP and proteomics (Supplementary Figure 4F), we detected Praja1-Usp7 interaction which is dependent on MAGE-D1 providing evidence for the MAGE-D1/Usp7/Praja-1 trimeric complex. MAGE-L2 has been previously shown to interact with both the ubiquitin ligase TRIM27 and Usp7 in a similar fashion to MAGE-D1 (*72*). Through its scaffold function, MAGE-D1 promotes Praja-1-mediated ubiquitination and degradation of BASP1 while simultaneously facilitating Usp7-mediated deubiquitination and stabilization of BASP1. How MAGE-D1 precisely coordinates and fine-tunes the balance between these opposing activities will be the focus of future studies. The development of inhibitory peptides or small molecules that disrupt the integrity of MAGE-D1 complex could be a viable strategy to attenuate behavioral responses to cocaine exposure. Collectively, our study reveals that MAGE-D1 regulates the synaptic protein BASP1 through ubiquitin-dependent mechanisms, controlling its stability and membrane localization while preserving membrane raft organization and actin cytoskeleton integrity. These findings establish the MAGE-D1/BASP1 axis as a previously unrecognized pathway linking ubiquitin signalling to the structural organization of synaptic membranes and provide a conceptual framework for understanding how ubiquitin regulatory scaffolds can shape neuronal architecture and synaptic plasticity.

## Acknowledgements

We thank the proteomics core facility at the EMBL for their support in data collection and analysis of mass spectrometry data shown in this manuscript. We also thank the Eukaryotic expression facility (EEF) at EMBL Grenoble for support in creating cell lines and maintaining the cell culture and the light microscopy infrastructure. This work used the platforms of the Grenoble Instruct-ERIC centre (ISBG) and we thank Caroline Mas for biophysics platform support. We thank D. Houtteman, S. Laghmari, P. Hagué, and L. Cuvelier for the technical assistance and the mouse colonies in Brussels. We thank Julian Cheron for his advices and guidance regarding the Usp7 inhibitor treatment in mice. We thank the High-throughput crystallisation facility at EMBL Grenoble for crystallization of MAGE-D1/Praja1 fusion construct. We thank Dr. Andrew McCarthy for supporting X-ray diffraction data collection. We thank the M4D facility at the Institut de Biologie Structurale (IBS) Grenoble, specifically Jean-Philippe Kleman and Oleksandr Glushonkov for their support in confocal and super resolution imaging. We thank Rahul Sharma and Michael Adams and other formers members of the Bhogaraju group in assisting with purification and molecular cloning of MAGE-D1 MHD construct. A.K.E. is a Research Director of the FRS-FNRS and a Welbio investigator. Funding: this work was supported by grants from FRS-FNRS (#23587797,#33659288, and #33659296), Welbio (#30256053, #X), ARC from FWB, Fondation ULB, AXA Research Fund (chaire AXA) to A.K.E. “*A unified approach for identifying ubiquitin ligase substrates in cellulo and in vivo”* project is funded by Impulscience® from Fondation Bettencourt Schueller awarded to S.B. The EMBL EIPOD program funded T.C. position.

## Author contributions

T.C and S.B conceived the study. T.C performed all the biochemical, imaging experiments with SH-SY5Y cells and the mouse brain samples. T.C also prepared samples for all the proteomics experiments. V.K performed purification and crystallization of MAGE-D1 MHD. S.B. determined the crystal structure. W.M. did mice handling, Usp7 inhibitor injections, behavioural recordings and thalamus dissections. S.G. performed BASP1 pulldown experiments, assisted M.P in creating MAGE-D1 KO cell lines and provided general laboratory support. N.A. performed AlphaFold predictions of MAGE-D1 and Usp7 interaction, performed ITC and *in vitro* ubiquitination experiments. M.R. supported the proteomics experiments. A.K.E acquired funding and supervised the mice work. S.B acquired funding and supervised the entire study. S.B and T.C wrote the manuscript with input from all the authors.

## Data availability

The mass spectrometry proteomics data have been deposited to the ProteomeXchange Consortium via the PRIDE partner repository (*73*) with the dataset identifier PXD080982.

## Supplementary Materials

Materials and Methods

Figures S1 to S4

Tables S1 to S12

