## Supplementary Materials for "A conserved ubiquitin signalling module controls membrane raft integrity at the neuronal synapse"

### Materials and Methods

#### Cell culture

Human embryonic kidney epithelial cells (HEK-293; American Type Culture Collection CRL-1573) were cultured in Dulbecco's Modified Eagle Medium (DMEM) supplemented with 10% (v/v) fetal bovine serum (FBS; Gibco™). Human neuroblastoma cells (SH-SY5Y; American Type Culture Collection HTB-11) were maintained in growth medium consisting of DMEM supplemented with 15% (v/v) heat-inactivated fetal bovine serum (hiFBS), 1× penicillin–streptomycin (Pen/Strep), and 6 mM L-glutamine (Gibco™). All cell lines were cultured at 37°C in a humidified incubator with 5% $CO_2$. SH-SY5Y cells were differentiated for 7 days following a previously published protocol (*29*). Briefly, from day 1 cells were cultured in differentiation medium consisting of DMEM supplemented with 2.5% (v/v) hiFBS, 1× Pen/Strep, 2 mM L-glutamine, and 10 μM all-trans retinoic acid (RA). The differentiation medium was replaced every 2 days until day 7. All cell lines used in this study were routinely tested and confirmed to be free of mycoplasma contamination by PCR-based detection.

#### Animals

The MAGE-D1-inactivated (MAGE D1-KO) mice used in the present study have been previously generated and described (*18*). Our experiments were conducted on adult (2-6 months-old) hemizygous MAGE D1 KO males and their wild-type littermates (MAGE D1 WT) generated by crossing heterozygous MAGE D1 KO/WT females with C57Bl6J males. Mice were group-housed (2-4 per cage) with access to food and water ad libitum and nesting materials. Temperature (20-24°C), humidity (45-65%) and light cycle (12h) were controlled in the facility. Animals were manipulated daily to habituate to the experimenter and weighed before the beginning of the behavioral experiment. Mice were randomly assigned to one of the 4 groups. Testing was performed during the light phase of the cycle. All experiments were performed according to the Institutional Animal Care Committee guidelines and were approved by the Local Ethical Committee (Comité d'Ethique et de Bien-Être Animal du pôle santé de l'Université Libre de Bruxelles (ULB), reference no. 872N).

**Reagents and antibodies**

All reagents used in this study are listed in Table 1 and reconstituted according to manufacturers' instructions. Polyclonal and monoclonal antibodies used for this study are listed in Table 2 and were used according to the manufacturers' recommended dilutions.

| **Reagent** | **Manufacturer** | **Catalog no.** | **Concentration and time of treatment** |
|---|---|---|---|
| MG132 | Calbiochem | 474787 | 10μM, 6 hours |
| USP7 Inhibitor HBX 41108 | Tocris (Biotechne) | 4285 | 10μM, 6 hours |
| USP7 inhibitor, P5091 | Selleck Chemicals | S7132 | 10mg/kg, 7 days |
| Cocaine hydrochloride | Fagron | 0244517 | 15mg/kg, 7 days |

Table 1: List of reagents

| **Antibody** | **Catalog no.** | **Manufacturer** |
|---|---|---|
| MAGE-D1 (for human cells) | PA599091 | ThermoFisher |
| MAGE-D1 (for mice) | 74-112 | Cosmo-Bio |
| BASP1 | 5H3L1 | ThermoFisher |
| Tubulin | NB100-690SS | Bio-Techne Ltd. |
| GAPDH | sc-32233 | SantaCruz |
| USP7 | 4833 | Cell signalling |
| PSD-95 | 3450 | Cell signalling |
| Synaptophysin | 17785-1-AP | Proteintech |
| FLOT-1 | 610821 | BD Transduction Laboratories |
| Transferrin-receptor 1 | H68.4 | ThermoFisher |
| Neurabin-1 | PA5-119842 | ThermoFisher |
| Neurabin-2 | PA5-48102 | ThermoFisher |
| DBN1 | 12243 | Cell signalling |
| Ubiquitin | sc-8017 | SantaCruz |
| Ubiquitin | 43124 | Cell signalling |
| HA | 3724 | Cell signalling |
| GFP | sc-9996 | SantaCruz |

Table 2: List of antibodies

**MAGE-D1 knockout cell lines generation**

HEK MAGE-D1 knockout (KO) cell generation it was previously described (*19*). For SH-SY5Y MAGE-D1 knockout cells, the sgRNA sequence was selected based on our previous study in HEK cells (*19*). Two independent sgRNA expression cassettes containing the selected target sequence were assembled by Gibson cloning into psBIG1a Cas9-GFP, generating the psBIG1a Cas9GFP_2xsgRNA construct. All plasmids were verified by restriction digest and Sanger sequencing (Eurofins). The verified psBIG1a Cas9GFP_2xsgRNA plasmid was subsequently used to generate recombinant baculovirus. For baculovirus transduction, 0.5 x 106 Hek293T cells were seeded in one well of a multi-12well plate together with 200 uL concentrated baculovirions loaded with psBIG1a Cas9GFP_2xsgRNA. After 24h cells were trypsinized and seeded in a 10 cm dish. Four days later cells were dissociated and seeded at clonal density in one 10 cm dish (500 cells/dish). After 10 additional days individual clones visible by eye were manually picked and further expanded for genotyping. The day before transduction, 2 x $10^6$ SH-SY5Y cells/well were seeded in two wells of a multi-6well plate. For transduction, medium was exchanged by only 1.5 mL prior addition of 100 uL concentrated baculovirions loaded with psBIG1a Cas9GFP_2xsgRNA. After 8h transduced cells were trypsinized and seeded back in wells of a multi-6well plate. Two days after cells were detached, resuspended in DMEM and GFP+ cells were isolated by fluorescence activated cell sorting (FACS) by flow cytometry on an Aria IIu (BD Biosciences). Cell discrimination was based on fluorescence at 530/30 nm from 488 nm excitation. Single cells were collected in multi-96well plate in DMEM 35% FBS + 25% conditioned medium. Additionally, 1500 GFP+ cells were collected as a pool in one well of a multi-24well plate. Cells in plates, were further expanded for genotyping. Genomic DNA extraction was done on clones growing in multi-12well plate using the Monarch® Genomic DNA Purification Kit (New England Biolabs, cat. no. T3010S) and following the manufacturer recommendation. Genotyping was achieved by running PCRs on extracted gDNA of each individual clone. Couple of primers used for genotyping the deletion were P1/P2 whereas intact gene body (WT band) was amplified using P1/P3. P1: GCACAGTCAGACCACAGTCACC; P2: GAGCCAACCATGGAAAGGAGGG and P3: CTGCGGTGGCCTGGTTAGTAG with P3 annealing within the MAGE-D1 region meant to be deleted. Two homozygous clones were obtained and the abolished expression of MAGE-D1 at protein level was confirmed by WB.

**Endogenous MAGE-D1 Immunoprecipitation in human cells**

For endogenous MAGE-D1 IP in HEK and SH-SY5Y cells, the MAGE-D1 KO cell line was used as control. Wild-type and MAGE-D1 KO cells were grown in triplicates in 150 mm dishes (20 $\times 10^6$ cells). Cells were harvested from the culture dishes by trypsinization and lysed with lysis buffer (50 mM Tris–HCl pH 7.5, 150 mM NaCl, 1% Triton X-100) supplemented with protease inhibitors (Roche, cat. no. 11836170001). The cell lysate was incubated for 30 min at 4 °C, sonicated for 5 min at 0 °C, and then centrifuged at 15,000 × g for 15 min. The total protein concentration was determined by BCA assay (ThermoFisher Scientific) and depending on total protein recovered between 0,5 – 2 mg of total proteins were incubated for 1 h (4 °C, rotary agitation) with 50 µL of Pierce™ Protein A Agarose beads (ThermoFisher, cat. no. 20333), centrifuged at 2,500 g for 3 min at 4 °C and the supernatant recovered as pre-cleared lysate. For the immune complex formation (total proteins + antibody), the pre-cleared lysate was incubated overnight (4 °C, rotary agitation) with 2.5 µg of MAGE-D1 antibody. To reduce non-specific binding and background, 50 µL of protein A Agarose beads were incubated for 1 h with 0.1% of BSA (4 °C, rotary agitation) and washed two times with IP buffer (50 mM Tris–HCl pH 7.5, 150 mM NaCl). The pre-blocked protein A beads were added to the immune complex at 4 °C for 4 h under rotary agitation, centrifuged at 2500 × g for 3 min at 4 °C and washed three times with IP buffer and one time with 10mM Tris-HCl pH 7.5. The immune complex bound to the beads was eluted in 60 µL of Laemmli buffer 2X (Biorad cat.no. 1610747) incubated for 10 min at 95 °C, centrifuged at 2500 × g for 3 min, and analyzed by MS.

**USP7 Inhibitor treatments**

SH-SY5Y cells were differentiated for 7 days. On day 7, the differentiation medium was replaced with fresh medium supplemented with 10 µM of USP7 inhibitor (HBX 41108, 100 mM stock solution in DMSO). Control cells received an equivalent volume of DMSO (10 µL) as a vehicle control. Cells were then incubated for 6 h at 37 °C in a humidified incubator with 5% $CO_2$. Following treatment, cells were harvested by trypsinization and lysed in lysis buffer (50 mM Tris-HCl, pH 7.5, 150 mM NaCl, 1% Triton X-100, 100 mM N-ethylmaleimide [NEM]) supplemented with protease inhibitors (Roche, cat. no. 11836170001). Lysates were incubated for 30 min at 4 °C, sonicated for 5 min on ice, and centrifuged at 15,000 g for 15 min (4 °C). Protein concentrations were determined using the BCA Protein Assay (Thermo Fisher Scientific), and equal amounts of protein were analyzed by Western blot.

For mice treatment, the USP7 inhibitor P5091 was solubilized in 15% DMSO + 20% SBE-β-CD in saline buffer and sonicated for 10 min at 37°C prior to use. P5091 or vehicle (5% DMSO + 20% SBE-β-CD in saline) were injected daily intraperitoneally (i.p.) at 10mg/kg for seven days. After behavioural test recording mice were anesthetized (1.25% Avertin (2,2,2-tribromoethanol) in saline, i.p., 0.2mL/10g) and decapitated. Brains were quickly extracted, transferred to ice-cold PBS where the thalamus was micro dissected. The tissue was mechanically dissociated using Dounce tissue grinder and lysed in lysis buffer (50 mM Tris-HCl pH 8.0, 150 mM NaCl, 1 mM EDTA, 0.5 % NP-40; supplemented by protease inhibitor (Complete, Roche) and phosphatase inhibitor (PhosSTOP, Roche). After centrifugation at 15,000 g for 20min (4 °C), protein concentrations were determined using the BCA Protein Assay and equal amounts of protein were analyzed by Western blot.

**Mouse behavioural test and statistical analysis**

A 2-injection cocaine-induced locomotor sensitization protocol was performed as previously described (*19*, *86*). In total 24 mice were separated into 4 experimental groups as follows: Vehicle + Saline, USP7 inhibitor P5091 + Saline, Vehicle + Cocaine and USP7 inhibitor P5091 + Cocaine (N=6/group, cocaine injected at 15mg/kg, i.p., and P5091 (Selleck Chemicals, S7132) at 10mg/kg, i.p.). Briefly, mice were injected every day for 11 days with the appropriate combination of treatments and their locomotor behavior was recorded on day 1-3 (habituation, saline only) and on day 4 and 11 (test days, saline or cocaine, the same treatment was administered twice, 7 days apart). On the two test days (day 4 and day 11), control mice receiving saline were also tested. No significant differences were observed (data not shown). The test consists of a 30 min recording of the locomotor activity (distance traveled in meters) in an open field arena analyzed using the automated tracking system Ethovision XT 19 (Noldus Information Technology) connected to a top camera in low light conditions (20Lux, to avoid generating an anxiogenic environment). Data are shown as mean ± SEM. Behavior data were analyzed using GraphPad Prism 8 software. Normality has been assessed by two-way repeated-measures ANOVA. Sidak's post-hoc tests was performed, as indicated in the figures. Differences were considered significant when $p < 0.05$. Right after the last behavioral recording (D11), mice were deeply anesthetized, decapitated and the brains were transferred to ice cold PBS to perform thalamus microdissection.

**Synaptosome and PSD isolation from mouse thalamus**

Synaptosomal and PSD fractions were extracted from mouse thalamus as previously described (*87*). In brief, thalamic tissues of USP7 Inhibitor treated mice and littermate controls were dissected and homogenized on ice with a Dounce tissue grinder in 10 volumes of cold sucrose buffer (0.32 M sucrose and 25 mM Hepes pH 7.4) with protease inhibitors. Homogenates were centrifuged at 710g for 10 min at 4°C to isolate supernatant from large debris and nuclei. The supernatant fraction was centrifuged at 10,000g for 15 min at 4°C; then, the supernatant containing light membrane and cytosolic fraction was discarded, and the pellets were washed twice with cold sucrose buffer and resuspended in cold Hepes-buffered saline (HBS) containing 25 mM Hepes and 150 mM NaCl (pH 7.4) to obtain synaptosome fractions. PSD-enriched fractions were prepared by solubilizing synaptosomes in 1% Triton X-100 (HBS) at 4°C for 30 min and, subsequently, centrifuging at 10,000g for 20 min. The pellet was resuspended in 3% SDS in HBS to yield PSD-enriched fractions. Total protein concentration in brain homogenates was measured using BCA assay. Protein lysates were further analyzed by Western Blot.

**Membrane rafts isolation from mouse brain**

Membrane rafts were isolated from total mice brain as previously described (*88*). Briefly, mice were euthanized and whole brain tissue was rinsed with saline solution and homogenized with a Dounce tissue grinder in cold homogenization buffer (10 mM Tris–HCl, pH 7.4, 5 mM ethylenediamine tetra acetic acid (EDTA), 320 mM sucrose) with protease inhibitors. After low-speed centrifugation (800 g, for 10 min) to remove nuclei and residual particulate material, supernatants were spun at 10,000 x g at 4ºC for 15 min and resulting pellets dissolved in cold buffer of 50 mM Tris-HCl pH 7.4, 150 mM NaCl, 5 mM EDTA, 1% Triton X-100 (v/v) and incubated on ice (10 min). Solubilized pellets were adjusted to 40% sucrose and fractionated on discontinuous sucrose gradients by centrifugation at 34,000 rpm (SW40 Ti; Beckman Coulter) for 16 h. Between eleven to thirteen fractions (1 ml each) were collected from each gradient starting from the top and an aliquot of each analyzed by Western blot.

**Cell surface protein isolation**

Cell surface protein isolation was performed by using Pierce™ Cell Surface Protein Biotinylation and Isolation Kit (A44390, ThermoFisher) following manufacturer protocol. SH-SY5Y WT and MAGE-D1 KO cells were grown in triplicates in 150 mm dishes ($20 \times 10^6$ cells) and differentiated for 7 days. In the last day of differentiation, cells were labeled with Thermo

Scientific™EZ-Link™ Sulfo-NHS-SS-Biotin, a thiol-cleavable amine-reactive biotinylation reagent, for 10 min at RT. Cells were subsequently lysed with detergent and the labeled proteins were then isolated with Thermo Scientific™ NeutrAvidin™ Agarose. The bound, labeled proteins were released by reduction of the disulfide bond with 10 mM DTT and analyzed by MS.

**Transient transfection and co-Immunoprecipitations in HEK cells**

Mammalian constructs used for transient co-transfection and co-IP in HEK cells included GFP-MAGED1-MHD, GFP-MHD_G552N, GFP-MHD_L560D, GFP-MAGED1-Unstructured, GFP-MAGED1, GFP-MAGED1_V569D, Praja1-HA, Praja1_F514D_F517D-HA, Praja-1_ΔR-HA (1–563), Praja-1_Δ1HR-HA (1–531), Praja-1_Δ3HR-HA (1–500), Praja2-HA, Praja2_F553D_F556D-HA, GFP-Praja1, GFP-Praja2, Myc-Ubiquitin, and BASP1-HA. All plasmids were transfected into HEK-293 cells using Polyethyleneimine Max (#24765-100, Polysciences), according to the manufacturer's instructions. Briefly, cells were seeded the day before transfection. The following day, equal amounts of two constructs were mixed with PEI MAX in reduced-serum medium, and the transfection mixture was incubated at room temperature for at least 20 min before being added dropwise to the cells. Transfected cells were incubated overnight at 37°C with 5% $CO_2$. For in cell ubiquitination assay cells, after 20 hours of transfection cells were treated with 20 µM of proteasome inhibitor MG132 for 4 hours. For GFP-IP experiments, GFP-Trap® Agarose beads (ChromoTek), while for HA-IP Monoclonal Anti-HA−Agarose antibody (A2095, Millipore) were used following manufacturer protocol. In brief, cell lysates were prepared as previously described for SH-SY5Y cells and 1 mg of total proteins were incubated with 60 µL of beads, previously equilibrated and washed, for 2 hours at +4°C. After three washes, samples were eluted in 2x SDS-sample buffer (Laemmli), and analyzed by Western Blot or MS.

**Immunoblotting**

The cell lysates were run on 4-20% Tris glycine gradient gels (Bio-Rad) by SDS-PAGE followed by transfer to polyvinylidene difluoride (PVDF) or nitrocellulose membranes. After the transfer, the membranes were incubated in blocking buffer (5% BSA in PBS) for 1h at RT followed by overnight incubation with the antibodies listed in table 2 at 4 °C in primary buffer (5% BSA, 0.1% Tween in PBS). Next day, the membranes were incubated in secondary buffer (5% BSA, 0.1% Tween, 0.01% SDS in PBS) with fluorescent tagged secondary antibodies (1:20000) for 1h at RT followed by imaging on a fluorescence scanner imaging system (Biorad or Licor System). For HRP-conjugated secondary antibodies (1:5000), membranes were incubated for 1 hour at RT in primary buffer followed by detection with a Immobilon Crescendo Western HRP substrate (WBLUR0100, Millipore). Imaging was done using Image lab software (v5.2.1) in ChemiDoc™MP Imaging System (Biorad).

**Immunostaining, confocal microscopy and statistical analysis**

Undifferentiated SH-SY5Y cells were seeded onto 22 mm poly-D-lysine-coated coverslips in 35 mm dishes and differentiated for 7 days. On the final day of differentiation, cells were washed once with PBS and fixed using different protocols depending on the target protein. For BASP1 immunostaining, cells were fixed in 3% paraformaldehyde (PFA) and 0.1% glutaraldehyde (GA) in PBS for 10 min at room temperature (RT) in the dark, followed by reduction with 0.1% sodium borohydride ($NaBH_4$) in PBS for 7 min. After three washes with PBS, cells were blocked and permeabilized in Solution 1 (3% BSA, 0.1% Triton X-100 in PBS) for 1 h at RT. Following one wash with PBS, cells were incubated overnight at 4°C with anti-BASP1 primary antibody (1:100 dilution) prepared in Solution 2 (3% BSA in PBS). For Ganglioside GM1 staining cells were incubated with 1 µg/mL Alexa Fluor 647-conjugated recombinant cholera toxin subunit B (AF647-CTB, C-34778 Molecular Probes) in Solution 2 for 30 min at 4°C in the dark. Cells were then washed twice with PBS and fixed with 4% PFA for 20 min at RT. For GM1/MAGE-D1 co-staining, after fixation, cells were subsequently incubated with anti-MAGE-D1 primary antibody (1:100 dilution) in Solution 2 for 2 h at RT. For FLOT-1 immunostaining, cells were fixed in ice-cold 100% methanol for 5 min at −20°C. After three washes with PBS, cells were incubated overnight at 4°C with anti-FLOT-1 primary antibody (1:100 dilution) in Solution 2. For all immunostaining experiments, following primary antibody incubation, cells were washed three times with PBS and incubated with Alexa Fluor 647 (BASP1, FLOT-1) or Alexa Fluor 555 (MAGE-D1) conjugated secondary antibodies (1:200 dilution) for 1 h at 37°C in a humidified chamber protected from light. Cells were then washed three times with PBS and mounted using ProLong™ Antifade Mountant containing DAPI. Spinning-disk confocal microscopy was performed using an Olympus IX81 inverted microscope equipped with an iLaunch laser bench (GATACA Systems) coupled to a Yokogawa CSU-X1 confocal head, using a 60X (CTB, FLOT-1, MAGE-D1) 1.42-NA or 100X (BASP1) 1.49-NA oil-immersion apochromatic objective. Fluorescence excitation was performed at 642 nm for AF647, at 561 nm for AF555 and at 405 nm for DAPI. Fluorescence emission was collected with an Andor iXon Ultra EMCCD camera through a quad-band Semrock™ Di01-T405/488/568/647 dichroic mirror and single-band emission filters (Chroma) adapted to each fluorophore used: ET440/40m for DAPI, ET600/50m for AF555 and ET700/75m for AF647. Emission filters were selected to minimize crosstalk between the AF 555 and AF647 channels, and no bleed-through into the AF 647 channel was detected. Data acquisition was performed using Metamorph 7.10 (Molecular devices). For statistical analysis, total no. of WT cells and the no. of MAGE-D1 KO cells were counted from at least 5 different fields for each condition and percentage of cells showing membrane localization of the relative staining was calculated. For BASP1 three biological replicates, for FLOT-1 two biological replicates and for CTB five technical replicates were used. Percentage difference of membrane localization between WT and MAGE-D1 KO cells were calculated using GraphPad prism 11. Statistical significance of data was calculated by an unpaired T test, and p-values are reported in the figures. To determine colocalization between CTB (red) and MAGE-D1 (orange hot) images were analyzed in FIJI (*89*). Colocalization analysis was done using Coloc2 plugin of ImageJ. The Pearson's correlation coefficient was calculated on a cell-by-cell basis. Results are indicative of 74 cells counted from 12 different field of views.

**Immunostaining, single-molecule localization microscopy (SMLM/dSTORM) and processing**

Undifferentiated SH-SY5Y cells were seeded onto 24 mm high-precision No. 1.5H cover glasses (170 ± 5 µm thickness; Marienfeld) coated with poly-D-lysine in 35 mm dishes and differentiated for 7 days. On the final day of differentiation, BASP1 immunolabelling was performed following the same protocol described for confocal microscopy, using AF647. Alternatively, for F-actin staining, after permeabilization in Solution 1 cells were incubated with Phalloidin-iFluor 647 (ab176759, Abcam) for 20 minutes in a humidified chamber at 37°C, protected from light. The cover glasses with labeled cells were then mounted on cavity slides (475505, Brand Gmbh), containing the STORM buffer, and sealed with a silicon sealant Twinsil (Picodent). The STORM buffer was composed of 50 mM Tris-HCl pH 8.0, 10 mM NaCl, 100 mM MEA (30070, Sigma), GLOX oxygen scavenger (560 µg/ml glucose oxidase (G2133, Sigma), 34 µg/ml catalase (C40, Sigma)) and 10% w/v of D-glucose. dSTORM data were acquired on an Olympus IX83 inverted motorized microscope equipped with a SAFe360 SMLM module (Abbelight), and a UPLXAPO 100x NA 1.5 oil immersion objective. The sample was excited with a 642 nm laser at 400 W/cm$^2$ under continuous HiLo illumination. The emitted signal was filtered with a dichroic mirror Di03-R405/488/532/635-t1, a multiband band-pass FF01-446/510/581/703 and a more specific emission filter FF01-698/70 (Semrock). The data were collected with an Orca Fusion sCMOS camera (Hamamatsu) at a frame time of

50 ms. Typically, 40,000 – 60,000 frames were acquired per dataset under continuous z-drift correction using microscope's Zero Drift Control system (ZDC2 at 830 nm). The temperature of a blind cage incubator (Digital Pixel™) was set to 27°C. SMLM data were processed with the ThunderSTORM (*90*) plugin in Fiji using Integrated Gaussian fit with Weighted Least squares estimator (initial sigma 1.3 px, pixel size 100 nm). Further post-processing of the dataset was performed in ThunderSTORM to correct for possible lateral drift (cross-correlation), and to filter the localizations by sigma and uncertainty values to limit the number of artefacts (noise, double or out-of-focus localizations).

**Mass spectrometry and data analysis**

Protein samples were subjected to the SP3 protocol (*91*) and trypsin was added in a 1:20 ratio (protease: protein) in 50 mM N-2-hydroxyethylpiperazine-N-2-ethanesulfonic acid (HEPES) supplemented with 5 mM Tris(2-carboxyethyl) phosphine hydrochloride (TCEP) and 20 mM 2-chloroacetamide (CAA) for 5 hours at 37°C. Up to 10 µg of peptides were labeled using TMTsixplex™ (TMT6) or TMT10plex™(TMT9) reagent as previously described (*92*, *93*) Offline high-pH reversed-phase fractionation (*94*) was carried out using an Agilent 1200 Infinity high-performance liquid chromatography (HPLC) system, equipped with a Gemini C18 analytical column (3 µm, 110 Å, 100 x 1.0 mm, Phenomenex). An UltiMate 3000 RSLCnano LC system (Thermo Fisher Scientific) equipped with a trapping cartridge (µ-Precolumn C18 PepMap™ 100, 300 µm i.d. × 5 mm, 5 µm particle size, 100 Å pore size; Thermo Fisher Scientific) and an analytical column (nanoEase™ M/Z HSS T3, 75 µm i.d. × 250 mm, 1.8 µm particle size, 100 Å pore size; Waters) was used. Samples were trapped at a constant flow rate of 30 µL/min using 0.05% trifluoroacetic acid (TFA) in water for 6 minutes. After switching in-line with the analytical column, which was pre-equilibrated with solvent A (3% dimethyl sulfoxide [DMSO], 0.1% formic acid in water), the peptides were eluted at a constant flow rate of 0.3 µL/min using a gradient of increasing solvent B concentration (3% DMSO, 0.1% formic acid in acetonitrile. Peptides were introduced into an Orbitrap Fusion™ Lumos™ Tribrid™ mass spectrometer (Thermo Fisher Scientific) via a Pico-Tip emitter (360 µm OD × 20 µm ID; 10 µm tip, CoAnn Technologies) using an applied spray voltage of 2.4 kV. The capillary temperature was maintained at 275 °C. Full MS (MS1) scans were acquired in profile mode over an m/z range of 375–1,500, with a resolution of 60,000 at m/z (TMT6) or 120,000 at m/z (TMT9) 200 in the Orbitrap. The maximum injection time was set to 50 ms, the AGC target limit was set to 'standard'. The instrument was operated in data-dependent acquisition (DDA) mode, with MS/MS scans acquired in the Orbitrap at a resolution of 15,000 (TMT6) or 30,000 (TMT9). The maximum injection time was set to 54 ms (TMT6) or 94 (TMT9), with an AGC target limit of 200%. Fragmentation was performed using higher-energy collisional dissociation (HCD) with a normalized collision energy of 36%. MS2 spectra were acquired in profile mode. The quadrupole isolation window was set to 0.7 m/z, and dynamic exclusion was enabled with a duration of 60 seconds. Only precursor ions with charge states 2–7 were included for fragmentation. Raw files were converted to mzML format using MSConvert from ProteoWizard, using peak picking, 64-bit encoding and zlib compression, and filtering for the 1000 most intense peaks. Files were then searched using MSFragger version 3.7 in FragPipe version 19.1 against FASTA database UP000005640_HomoSapiens_ID9606_20594entries_26102022_dl11012023 containing common contaminants and reversed sequences. For the full scan (MS1) a mass error tolerance of 20 ppm and for MS/MS (MS2) spectra of 20 ppm was set. For protein digestion, 'trypsin' was used as protease with an allowance of maximum 2 missed cleavages requiring a minimum peptide length of 7 amino acids. The false discovery rate on peptide and protein level was set to 0.01. The standard settings of the FragPipe workflow ‘Default’ were used. MSFragger and FragPipe version used could vary for different proteomics dataset. Proteomics raw data were processed using either IsobarQuant (*95*), FragPipe (*96*), or DIA-NN with MaxLFQ quantification on gene level. Downstream statistical analyses were performed in the R programming environment (ISBN 3-900051-07-0) using Bioconductor workflows. Initial data processing included removal of contaminants and reverse protein identifications. For IsobarQuant-derived datasets, proteins quantified with at least two unique peptides (qupm ≥ 2) were retained, whereas FragPipe- and DIA-NN-derived datasets required at least two razor peptides (Razor.Peptides ≥ 2). Depending on the project, between 673 and 6,128 proteins passed quality-control filtering. Protein abundance values (corresponding to signal_sum, channel, or Intensity columns, depending on the software pipeline) were log2-transformed and assembled into either SummarizedExperiment or ExpressionSet objects for downstream analysis. For visualization purposes, protein abundances were normalized either to the median of control replicates or to the global median abundance across conditions. In immunoprecipitation experiments, prey protein intensities were additionally normalized to the bait protein abundance to assess enrichment relative to the bait. Differential protein abundance analysis was performed using moderated t-tests implemented in the limma package (lmFit and eBayes) (*97*). Replicate information was incorporated as a factor in the design matrix.

**Protein purification**

The Praja-1/MHD fusion construct used for crystallization consisted of MAGE-D1 residues 465-703 fused to Praja-1 residues 501-520 with a GSGS linker, a C-terminal 6xHIS tag and 3C protease cleavage site in a pET-21a (+) vector. The complex Praja1/MHD used for *in vitro* ubiquitination assay consisted of two constructs co-expressed: MAGE-D1 residues 465-703 in pCoofy1 vector, and Praja-1 residues 501-643 in a custom made pET vector. Both constructs have a N-terminal 6xHIS tag and 3C protease cleavage site. Full length UBE1 (Mus musculus, Uniprot: Q02053), UBE2D2 (Uniprot: P62837) and $USP7^{TUD}$ (aa556-1083, Uniprot: Q93009) construct were separately cloned into a pET28a (+) vector with a C-terminal 6xHIS tag and 3C protease cleavage site. For all purifications, the constructs were expressed in E. coli T7 Express (NEB) cells by growing the bacteria at 37 °C in lysogeny broth (LB), supplemented with the appropriate antibiotics, to an $OD_{600}$ of 0.6 and inducing the cells with 0.5 mM IPTG at 18 °C for 16 h. Following lysis in 50mM TrisHCl pH 7.5, 150 mM NaCl, 10% Glycerol (lysis buffer), the soluble fraction was incubated with TALON® resin (Takara) for 2 h at 4 °C and eluted with 200 – 400 mM imidazole, following numerous washes with lysis buffer and low amounts of imidazole. $USP7^{TUD}$ was subjected to anion exchange chromatography (AIEX) and subsequently to size-exclusion chromatography (SEC). Protein fractions were eluted in 20 mM HEPES pH 7.5, 150mM NaCl, and 1 mM DTT (SEC buffer). The fusion construct was subjected to SEC in the buffer 20 mM MES, 250 mM NaCl, 50 mM KCl and 5% Glycerol, pH 6.0. The complex Praja1/MHD was subjected to SEC in the buffer 10mM HEPES, 150mM NaCl, 5% Glycerol and 1mM TCEP at pH 7.5.

BASP1 was purified as previously described (*98*). Construct of full length BASP1 (Uniprot: P80723) was cloned into a pET-21a (+) vector (GenScript) and pNMT cloned into a pRSF-1 vector (Addgene, #42578). The two constructs were co-expressed in E. coli T7 Express (NEB) cells by growing the bacteria at 37 °C in lysogeny broth (LB), supplemented with the appropriate antibiotic, to an $OD_{600}$ of 0.6. 50μM of Myristic acid (Sigma Aldrich) was added to the growth media for 10 min before induction with 0.8 mM IPTG at 18 °C for 4 h. Following lysis in 10mM HEPES, 0.2 mM EGTA, pH 7.5 (lysis buffer), and sonication on ice the supernatant was recovered by centrifugation at 40,000 g at 4°C for 30 min. 2.5% of Perchloric acid was added to the supernatant, incubated for 5 min on ice and recovered by centrifugation at 40,000 g at 4°C for 30 minutes. The supernatant was then neutralized using NaOH and adjusted to pH 7.5 before concentration in a Amicon® Ultra Centrifugal Filter, 10 kDa MWCO.

**AlphaFold prediction**

To understand the interaction between MAGED1 and $USP7^{TUD}$, various peptides of different lengths (between 12-14aa) along the $MAGED1^{PBR}$ region containing the KxKxxxK TUD binding motif were modelled against our $USP7^{TUD}$ domain construct using AlphaFold3 (AF3) (*99*). While all AF3 outputs were screened, only the highest confidence prediction of the interaction (using MAGED1 peptide aa243-256, Uniprot: Q9Y5V3) as determined by ipTM and pTM scores was then used as a structural basis for the design of a peptide for ITC validation.

**Isothermal Titration Calorimetry**

$MAGE\text{-}D1^{PBR}$ peptide (aa241-255) was synthetized by GenScript with a HPLC purity of 95.8%. $USP7^{TUD}$ and $MAGE\text{-}D1^{PBR}$ were dialyzed in the same buffer containing 10mM HEPES, pH8.0, 100mM NaCl, and 1mM TCEP. The protein concentration was determined post-dialysis using BCA Assay. 200µl of wild-type $USP7^{TUD}$ (25µM) were titrated against 38µl of wild-type $MAGE\text{-}D1^{PBR}$ (250µM) using a MicroCal PEAQ-ITC (Malvern Panalytical) at 18°C. Injections per ITC run were 16, one 0.5µl injection at the start, followed by 15 2.5µl injections, with 60s initial delay, 5s injection duration, and 180s injection spacing. Reference power was 5 µcal/s with the stir speed set to 750rpm. Duplicate runs were done of each condition with statistical tests being employed to decide whether all the runs could be used to determine the binding affinity. Normally distributed synthetic data was generated based of the mean and standard deviation of each run, the normality of which was confirmed by Shapiro–Wilk test ($p > 0.05$), before undergoing the Fischer test ($p > 0.05$) to check for the homogeneity of variance between runs. Student t-test ($p > 0.05$) was employed to determine that the data from each run belonged to the same population.

**Crystallization, data collection and processing**

The fusion protein of Praja-1/MHD was purified as described above and concentrated to 3.5 mg/ml. Protein was used to set up 2 µL crystallization drops by sitting drop vapor diffusion method using 1 µL of purified protein and 1 µL of mother liquor which consisted of 0.1 M HEPES MOPS, 0.06 M Morpheus Divalents, 30% v/v Glycerol PEG 4000 along with 1X Hampton additives Sodium Acetate and Ammonium Chloride. The drops were incubated for 7 days at 20°C, which led to the formation of needle like crystals of considerable size. The crystals were then harvested and transferred to a cryo-preservant buffer which consisted of 0.1 M HEPES MOPS, 0.06 M Morpheus Divalents, 50 % v/v Glycerol PEG 4000 along with 1X Hampton additives Sodium Acetate and Ammonium Chloride. The crystals were diffracted at the European Synchrotron Research Facility (ESRF) at EMBL beamline ID23 operated with MXCuBE3 to obtain diffraction data (Supplementary Table 12). The data was processed using AutoPROC (*100*) and STARANISO and indexed in space group P 42212. The structure was determined with molecular replacement (MR) using a AlphaFold3 predicted model of MAGE-D1 MHD. MR was performed using the PHASER module as available in PHENIX (*101*) package. Iterative model building and refinement were done using COOT (*102*) and Phenix refine as available in Phenix package.

**In vitro Ubiquitination assay**

For examining BASP1 ubiquitination in the presence of Praja-1/MHD, 10μM BASP1, 10 μM Praja-1/MHD, 0.6 μM E1, 6 μM E2 (UBE2D2), 80μM Ubiquitin (R&D Systems, #U-100H) and 5mM ATP were incubated in presence or absence of Praja-1/MHD complex in the buffer 50 mM Tris pH 6.5, 50 mM NaCl, 0.5 mM DTT and 5 mM MgCl2. Concentration of proteins was determined by BCA assay. The reaction was incubated for 1 h at 35°C. Samples were run on a SDS PAGE gel, followed by Western blot or in-gel digestion for MS analysis as described below.

**BASP1 band in gel-digestion for MS analysis**

The protein bands of interest were excised from the gel as previously described (*103*). Gel pieces were first dehydrated, reduced, alkylated and again dehydrated with 100% acetonitrile. Proteins were enzymatically digested overnight at 37°C using trypsin at a concentration of 2 ng/μL in 50 mM ammonium bicarbonate. Peptides were extracted by sonication for 15 minutes, followed by centrifugation and collection of the supernatant. For optimal peptide recovery, gel pieces were subjected to a second extraction step using a solution composed of 50% acetonitrile and 1% formic acid in ultrapure water. Samples were again sonicated for 15 minutes, centrifuged, and the supernatants combined with the initial extracts. The pooled peptide extracts were dried using vacuum centrifugation. Dried peptides were reconstituted in a solution containing 4% acetonitrile and 1% formic acid prior to LC-MS/MS analysis.

### Supplementary figures

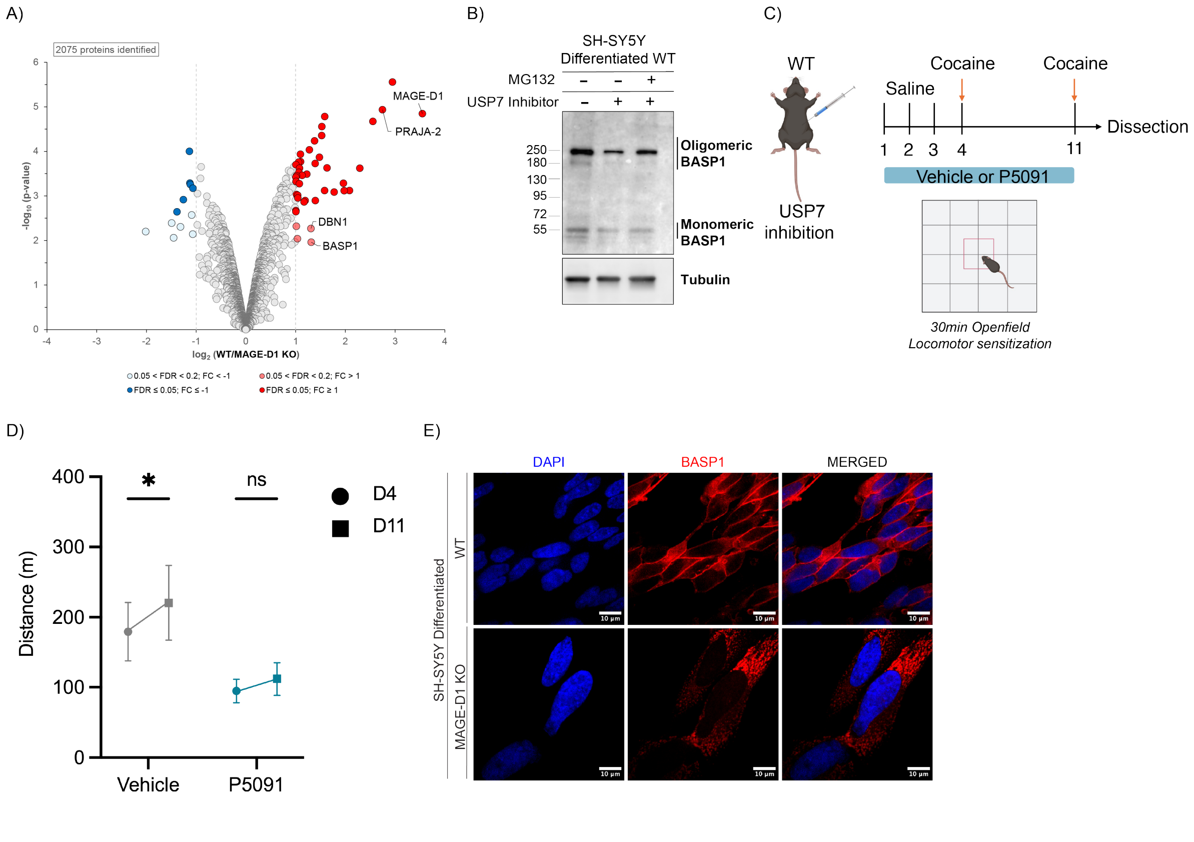


Supplementary figure 1

### Supplementary figure 1:

A) Volcano plot of total protein levels isolated after endogenous immunoprecipitation of MAGE-D1 from WT and MAGE-D1 KO HEK cells. Results are representative of three independent biological replicates. Significantly altered proteins are shown in dark red or blue (FDR ≤ 0.05, log2FC ≥I1I) and light red or blue (0.2 <FDR <0.05, log2FC > I1I) (moderated t-test).

B) Immunoblotting of BASP1, and Tubulin (loading control) from total lysate of differentiated SH-SY5Y WT cells under vehicle control condition (DMSO) or USP7 inhibition (HBX 41108, 10uM) or USP7 inhibition (HBX 41108, 10uM) and the proteasome inhibitor MG-132 (10uM) for 6 hours. Oligomeric and monomeric BASP1 forms are indicated.

C) Schematic of the protocol used for the cocaine-induced locomotor sensitization behavioral test. Images were created with Biorender. Experimental procedures are described in Materials and Methods.

D) Cocaine-induced locomotor sensitization. Cocaine administration 15 mg/kg, intraperitoneal injection. Vehicle (5% DMSO + 20% SBE-β-CD in saline) or USP7 Inhibition (P5091) administration 10 mg/kg; intraperitoneal injection. Vehicle treated mice n=6, USP7 Inhibitor treated mice n=5, two-way repeated measures ANOVA: Day Effect: $p$=0,0218 (*); Treatment Effect: $p$= 0,1118 (ns); Interaction Day x Treatment: $p$=0,2857 (ns). Posthoc (Sidak's multiple comparison), D4 vs D11: $p$=0,0352 (*) for Vehicle and $p$=0,5064 (ns) for P5091 group, respectively.

E) Confocal microscopy images of differentiated SH-SY5Y WT and MAGE-D1 KO cells stained with DAPI (nuclei, blue) and immunostained with anti-BASP1 antibody. Scale bar 10 µm.

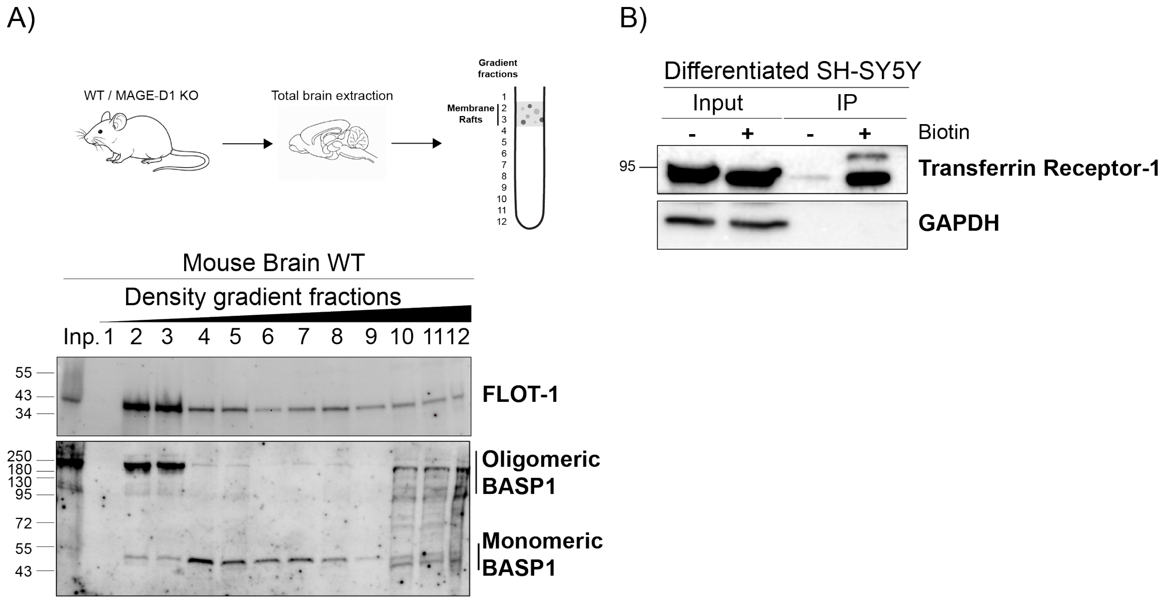


Supplementary figure 2

**Supplementary figure 2:**

A) Schematic representation of the protocol used for membrane raft isolation from total brain of mice (top panel). Immunoblotting of BASP1 and FLOT1 (membrane raft marker) from fractions collected during membrane raft isolation from total brain WT mice (bottom panel). Fractions 2 and 3 were pooled and used as the membrane raft fraction shown in Figure 2A. Oligomeric and monomeric BASP1 forms are indicated.

B) Immunoblotting of Transferrin receptor 1 and GAPDH from total lysate and enriched biotinylated protein fractions in presence (+) or absence (-) of biotin treatment in differentiated SH-SY5Y WT cells.

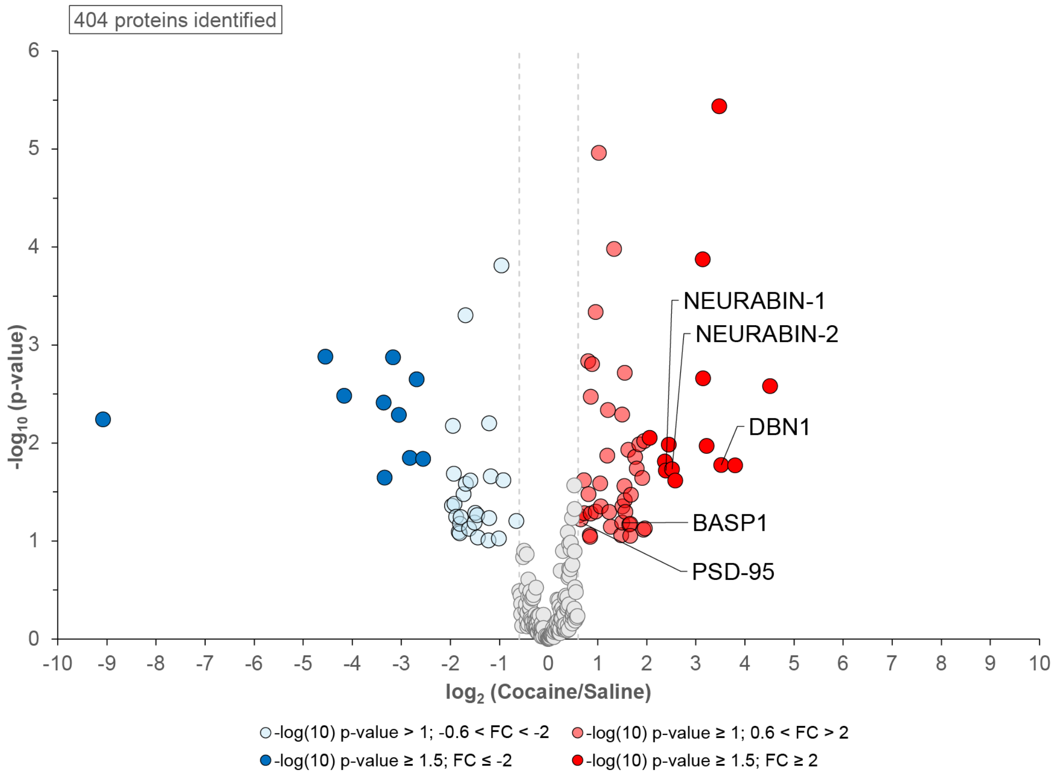


Supplementary figure 3

**Supplementary figure 3:**

A) Volcano plot of total protein levels isolated after endogenous immunoprecipitation of MAGE-D1 from thalamus of control mice treated with saline (n=5) and mice treated with cocaine (n=7) (20mg/kg, intraperitoneal). Results are representative of three independent biological replicates. Significantly altered proteins are shown in dark red or blue ($-\log_{10}$p-value≥1.5, $\log_2$FC ≥I2I) and light red or blue ($-\log_{10}$p-value≥1, I0.6I<$\log_2$FC > I2I) (moderated t-test). Data re-analyzed from reference 19.

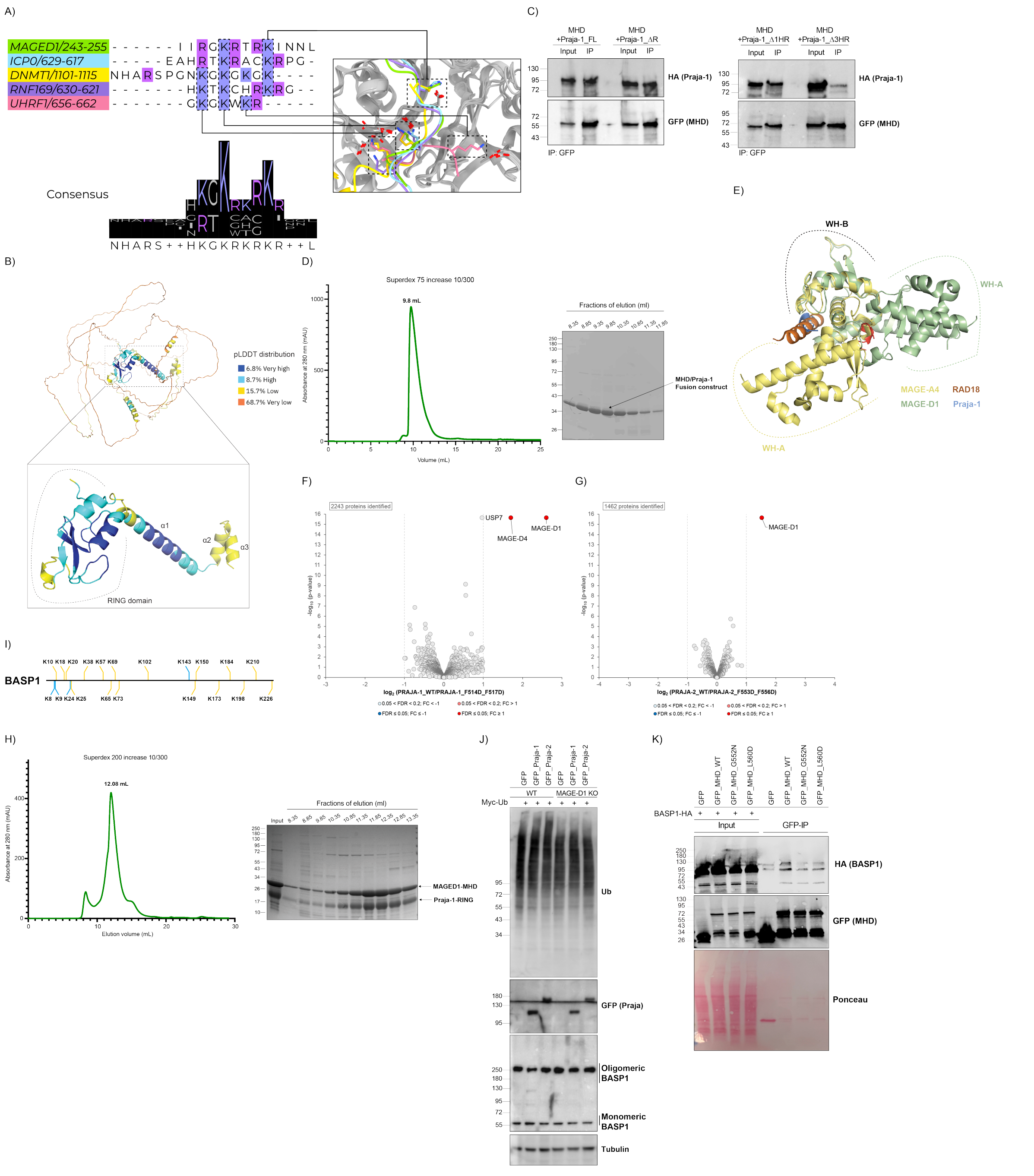


Supplementary Figure 4

### Supplementary figure 4:

A) Multiple Sequence Alignment of MAGE-D1 PBR against solved crystal structures of Usp7 TUD. Aligned lysine residues are indicated on overlayed crystal structure and AlphaFold3 prediction (Figure 4B). PBD: 5C56 (ICP0), 4YOC (DNMT1), 5GG4 (RNF169), 5C6D (UHRF1).

B) AlphaFold predicted structure of Praja-1. Inset of the C-terminal RING domain and the three helices adjacent to it. Structure colored based on pLDTT score.

C) Immunoblotting of Praja-1_HA deletion constructs and GFP-MHD following transient co-expression in HEK293 cells. GFP-MHD was immunoprecipitated from total cell lysates, and associated Praja-1_HA deletion constructs were detected by immunoblotting. The Praja-1 deletion constructs used are Praja-1 full length (FL), Praja-1_ **Δ**R (1-563), Praja-1_ **Δ**1HR (1-531), Praja-1_ **Δ**3HR (1-500).

D) Purification of MHD/Praja-1 fusion construct. Size-exclusion chromatography (SEC) profile of the MHD–Praja1 minimal MIR complex on a Superdex 75 Increase 10/300 GL column (left panel). Fractions corresponding to the elution peak were analyzed by SDS–PAGE followed by Coomassie Brilliant Blue staining (right panel). The expected MHD–Praja1 complex is indicated by the arrow.

E) Comparison of the ligase-binding clefts (LBC) of MAGE-A4 and MAGE-D1 by superimposition of the two structures: MHD-MAGED1/Praja-1 MIR and MHD-MAGEA4/RAD18 peptide (PDB: 9BD3).

F) Volcano plot of total protein levels isolated after HA immunoprecipitation of Praja-1_WT_HA and Praja-1_F514D_F517D_HA mutant over-expressed in HEK cells. Results are representative of three independent biological replicates. Significantly altered proteins are shown in dark red or blue (FDR ≤ 0.05, log2FC ≥I1I) and light red or blue (0.2 <FDR <0.05, log2FC > I1I) (moderated t-test).

G) Volcano plot of total protein levels isolated after HA immunoprecipitation of Praja-2_WT_HA and Praja-2_F553D_F556D_HA mutant over-expressed in HEK cells. Results are representative of three independent biological replicates. Significantly altered proteins are shown in dark red or blue (FDR ≤ 0.05, log2FC ≥I1I) and light red or blue (0.2 <FDR <0.05, log2FC > I1I) (moderated t-test).

H) Purification of MHD/Praja-1 minimal RING complex. Size-exclusion chromatography (SEC) profile of the MHD and Praja1-RING complex on a Superdex 200 Increase 10/300 GL column (left panel). Fractions corresponding to the elution peak were analyzed by SDS–PAGE followed by Coomassie Brilliant Blue staining (right panel). The expected MHD–Praja1 complex is indicated by the arrows.

I) BASP1 Ubiquitination sites. Newly identified ubiquitinated lysine residues are shown in blue. Ubiquitinated lysine residues previously reported in PhosphoSitePlus (*72*) and also identified in this study are shown in orange.

J) Immunoblotting of Ubiquitin, GFP_Praja-1, GFP_Praja-2, BASP1 and Tubulin (loading control) following transient co-expression of GFP_Praja-1 or GFP_Praja-2 with Myc-Ubiquitin in HEK WT and MAGE-D1 KO cells treated with 20 µM of proteasome inhibitor MG132 for 4 hours. Results are representative of two independent biological replicates.

K) Immunoblotting of BASP1_HA and GFP_MHD following transient co-expression of BASP1_HA and GFP_MHD_WT or GFP_MHD_G552N or GFP_MHD_L560D in HEK cells. GFP-MHD was immunoprecipitated from total cell lysates, and associated BASP1_HA was detected by immunoblotting. BASP1 migrated at an apparent molecular weight of >95 kDa. The electrophoretic mobility of BASP1 has been reported to vary depending on multiple factors, including SDS–PAGE conditions (*33*, *35*, *36*). Ponceau S staining was used as a loading control. Results are representative of three independent biological replicates.

**Supplementary tables**

| Gene | Protein.ID | Protein.Description | log2(FC) | -log10pvalue | fdr | Comparison |
|---|---|---|---|---|---|---|
| **DNMT3A** | Q9Y6K1 | DNA (cytosine-5)-methyltransferase 3A | 2,404129511 | 6,199324106 | 0,002140182 | WT - MAGE-D1_KO |
| **C7** | P10643 | Complement component C7 | 2,097359734 | 5,805062576 | 0,002140182 | WT - MAGE-D1_KO |
| **POGZ** | Q7Z3K3 | Pogo transposable element with ZNF domain | 1,880653698 | 5,757898125 | 0,002140182 | WT - MAGE-D1_KO |
| **ARID3B** | Q8IVW6 | AT-rich interactive domain-containing protein 3B | 1,901181364 | 5,623691638 | 0,002348309 | WT - MAGE-D1_KO |
| **GAL** | P22466 | Galanin peptides | 1,596778654 | 5,530048173 | 0,002348309 | WT - MAGE-D1_KO |
| **BASP1** | P80723 | Brain acid soluble protein 1 | 2,273163712 | 5,510558326 | 0,002348309 | WT - MAGE-D1_KO |
| **VGF** | O15240 | Neurosecretory protein VGF | 1,524986754 | 5,376362966 | 0,002348309 | WT - MAGE-D1_KO |
| **ZNF326** | Q5BKZ1 | DBIRD complex subunit ZNF326 | 1,559303505 | 5,376272813 | 0,002348309 | WT - MAGE-D1_KO |
| **H4C16** | P62805 | Histone H4 | 3,525093368 | 5,26123586 | 0,002348309 | WT - MAGE-D1_KO |
| **RSRC1** | Q96IZ7 | Serine/Arginine-related protein 53 | 1,30941798 | 5,254057506 | 0,002348309 | WT - MAGE-D1_KO |
| **SCG2** | P13521 | Secretogranin-2 | 1,986243734 | 5,226200544 | 0,002348309 | WT - MAGE-D1_KO |
| **PARVA** | Q9NVD7 | Alpha-parvin | 1,465107715 | 5,181301844 | 0,002348309 | WT - MAGE-D1_KO |
| **MACROH2A2** | Q9P0M6 | Core histone macro-H2A.2 | 1,773020093 | 5,168869599 | 0,002348309 | WT - MAGE-D1_KO |
| **SAFB** | Q15424 | Scaffold attachment factor B1 | 1,46453838 | 5,107061204 | 0,002348309 | WT - MAGE-D1_KO |
| **NRCAM** | Q92823 | Neuronal cell adhesion molecule | 1,556183845 | 5,094198074 | 0,002348309 | WT - MAGE-D1_KO |
| **RSF1** | Q96T23 | Remodeling and spacing factor 1 | 2,330068125 | 5,087452843 | 0,002348309 | WT - MAGE-D1_KO |
| **NAV2** | Q8IVL1 | Neuron navigator 2 | 1,597519374 | 5,084545383 | 0,002348309 | WT - MAGE-D1_KO |
| **SCGN** | O76038 | Secretagogin | 1,653849374 | 5,081774989 | 0,002348309 | WT - MAGE-D1_KO |
| **H3-3B** | P84243 | Histone H3.3 | 2,534586469 | 5,074761109 | 0,002348309 | WT - MAGE-D1_KO |
| **MATR3** | P43243 | Matrin-3 | 1,202126068 | 5,074140823 | 0,002348309 | WT - MAGE-D1_KO |
| **PCDH17** | O14917 | Protocadherin-17 | 1,614403498 | 5,029425147 | 0,002489808 | WT - MAGE-D1_KO |
| **BAZ1B** | Q9UIG0 | Tyrosine-protein kinase BAZ1B | 1,232713791 | 4,985231181 | 0,002641656 | WT - MAGE-D1_KO |
| **HNRNPH3** | P31942 | Heterogeneous nuclear ribonucleoprotein H3 | 1,180963866 | 4,866246281 | 0,003142651 | WT - MAGE-D1_KO |
| **CHGB** | P05060 | Secretogranin-1 | 1,040471763 | 4,826100467 | 0,003142651 | WT - MAGE-D1_KO |
| **HNRNPA1** | P09651 | Heterogeneous nuclear ribonucleoprotein A1 | 1,184032541 | 4,81073317 | 0,003142651 | WT - MAGE-D1_KO |
| **HNRNPA2B1** | P22626 | Heterogeneous nuclear ribonucleoproteins A2/B1 | 1,432733687 | 4,802838153 | 0,003142651 | WT - MAGE-D1_KO |
| **HNRNPR** | O43390 | Heterogeneous nuclear ribonucleoprotein R | 1,105700555 | 4,784719664 | 0,003142651 | WT - MAGE-D1_KO |
| **CHTOP** | Q9Y3Y2 | Chromatin target of PRMT1 protein | 1,188640207 | 4,771508723 | 0,003142651 | WT - MAGE-D1_KO |
| **PSD-93** | Q15700 | Disks large homolog 2 | 1,121151075 | 4,753929347 | 0,003176219 | WT - MAGE-D1_KO |
| **SAFB2** | Q14151 | Scaffold attachment factor B2 | 1,33346995 | 4,707930994 | 0,003430202 | WT - MAGE-D1_KO |
| **WDR43** | Q15061 | WD repeat-containing protein 43 | 1,033088855 | 4,636183443 | 0,003488651 | WT - MAGE-D1_KO |
| **HNRNPC** | P07910 | Heterogeneous nuclear ribonucleoproteins C1/C2 | 2,21585971 | 4,631877317 | 0,003488651 | WT - MAGE-D1_KO |
| **COCH** | O43405 | Cochlin | 1,486385276 | 4,595924398 | 0,003503075 | WT - MAGE-D1_KO |
| **GATA2** | P23769 | Endothelial transcription factor GATA-2 | 1,836711829 | 4,588248989 | 0,003503075 | WT - MAGE-D1_KO |
| **HMGA1** | P17096 | High mobility group protein HMG-I/HMG-Y | 1,18552087 | 4,563553538 | 0,003503075 | WT - MAGE-D1_KO |
| **FDFT1** | P37268 | Squalene synthase | 1,041058818 | 4,558150083 | 0,003503075 | WT - MAGE-D1_KO |
| **SAP18** | O00422 | Histone deacetylase complex subunit SAP18 | 1,548148682 | 4,517790491 | 0,003559241 | WT - MAGE-D1_KO |
| **SMARCA5** | O60264 | SWI/SNF-related matrix-associated actin-dependent regulator of chromatin subfamily A member 5 | 1,161712982 | 4,463145128 | 0,003559241 | WT - MAGE-D1_KO |
| **ANXA2** | P07355 | Annexin A2 | 1,077358441 | 4,448965037 | 0,003559241 | WT - MAGE-D1_KO |
| **RBMXL1** | Q96E39 | RNA binding motif protein, X-linked-like-1 | 1,475387044 | 4,443582288 | 0,003559241 | WT - MAGE-D1_KO |
| **TOP2B** | Q02880 | DNA topoisomerase 2-beta | 1,507555366 | 4,442701001 | 0,003559241 | WT - MAGE-D1_KO |
| **NEURABIN-1** | Q9ULJ8 | Neurabin-1 | 1,596231208 | 4,434614605 | 0,003559241 | WT - MAGE-D1_KO |
| **ELAVL1** | Q15717 | ELAV-like protein 1 | 1,321765583 | 4,430151603 | 0,003559241 | WT - MAGE-D1_KO |
| **RAP1GAP2** | Q684P5 | Rap1 GTPase-activating protein 2 | 1,097465432 | 4,407278325 | 0,003690856 | WT - MAGE-D1_KO |
| **ZMYND8** | Q9ULU4 | Protein kinase C-binding protein 1 | 1,183245589 | 4,398392126 | 0,003710075 | WT - MAGE-D1_KO |
| **ZNF384** | Q8TF68 | Zinc finger protein 384 | 1,329112626 | 4,386761541 | 0,003753898 | WT - MAGE-D1_KO |
| **HNRNPDL** | O14979 | Heterogeneous nuclear ribonucleoprotein D-like | 1,308310551 | 4,327099111 | 0,0038595 | WT - MAGE-D1_KO |
| **ACIN1** | Q9UKV3 | Apoptotic chromatin condensation inducer in the nucleus | 1,117357178 | 4,31997412 | 0,0038595 | WT - MAGE-D1_KO |
| **WDR75** | Q8IWA0 | WD repeat-containing protein 75 | 1,056911806 | 4,276441583 | 0,004055899 | WT - MAGE-D1_KO |
| **PNN** | Q9H307 | Pinin | 1,180410981 | 4,255872875 | 0,004055899 | WT - MAGE-D1_KO |
| **SMARCA1** | P28370 | Probable global transcription activator SNF2L1 | 1,183580206 | 4,225209276 | 0,004085657 | WT - MAGE-D1_KO |
| **H2AX** | P16104 | Histone H2AX | 1,299810878 | 4,225024707 | 0,004085657 | WT - MAGE-D1_KO |
| **MACROH2A1** | O75367 | Core histone macro-H2A.1 | 1,189662862 | 4,212386086 | 0,004129441 | WT - MAGE-D1_KO |
| **PSIP1** | O75475 | PC4 and SFRS1-interacting protein | 1,133626321 | 4,169267936 | 0,004322944 | WT - MAGE-D1_KO |
| **TOP2A** | P11388 | DNA topoisomerase 2-alpha | 1,308023841 | 4,118942053 | 0,004438001 | WT - MAGE-D1_KO |
| **CDC40** | O60508 | Pre-mRNA-processing factor 17 | 1,26896825 | 4,111893774 | 0,004468061 | WT - MAGE-D1_KO |
| **RBMX** | P38159 | RNA-binding motif protein, X chromosome | 1,565492717 | 4,090528013 | 0,004529178 | WT - MAGE-D1_KO |
| **ERI1** | Q8IV48 | 3'-5' exoribonuclease 1 | 1,052177341 | 4,089906696 | 0,004529178 | WT - MAGE-D1_KO |
| **RALY** | Q9UKM9 | RNA-binding protein Raly | 1,883549975 | 4,060775351 | 0,004632821 | WT - MAGE-D1_KO |
| **NSD3** | Q9BZ95 | Histone-lysine N-methyltransferase NSD3 | 1,061133477 | 4,032506323 | 0,004818706 | WT - MAGE-D1_KO |
| **MECP2** | P51608 | Methyl-CpG-binding protein 2 | 1,154530154 | 4,000714337 | 0,005024289 | WT - MAGE-D1_KO |
| **VTN** | Q3ZBS7 | Uncharacterized protein | 1,085589956 | 3,942304924 | 0,005327761 | WT - MAGE-D1_KO |
| **PPHLN1** | Q8NEY8 | Periphilin-1 | 1,139632947 | 3,9235941 | 0,005327761 | WT - MAGE-D1_KO |
| **CADM3** | Q8N126 | Cell adhesion molecule 3 | 2,273101622 | 3,843583519 | 0,00605853 | WT - MAGE-D1_KO |
| **BRD1** | O95696 | Bromodomain-containing protein 1 | 1,49604179 | 3,663450533 | 0,007389171 | WT - MAGE-D1_KO |
| **MOSPD2** | Q8NHP6 | Motile sperm domain-containing protein 2 | 1,191349366 | 3,602197275 | 0,007955649 | WT - MAGE-D1_KO |
| **PTPRN** | Q16849 | Receptor-type tyrosine-protein phosphatase-like N | 1,096664926 | 3,596472712 | 0,007958137 | WT - MAGE-D1_KO |
| **RAD21** | O60216 | Double-strand-break repair protein rad21 homolog | 1,028997256 | 3,556623999 | 0,008338052 | WT - MAGE-D1_KO |
| **HNRNPA3** | P51991 | Heterogeneous nuclear ribonucleoprotein A3 | 1,313949265 | 3,530222893 | 0,008583886 | WT - MAGE-D1_KO |
| **MBD1** | Q9UIS9 | Methyl-CpG-binding domain protein 1 | 1,274867346 | 3,223037047 | 0,012001166 | WT - MAGE-D1_KO |
| **NPTX2** | P47972 | Neuronal pentraxin-2 | 1,063209216 | 3,175060669 | 0,012630095 | WT - MAGE-D1_KO |
| **C11orf96** | Q7Z7L8 | Uncharacterized protein C11orf96 | 1,108068835 | 3,10875991 | 0,013637995 | WT - MAGE-D1_KO |
| **PLCXD3** | Q63HM9 | PI-PLC X domain-containing protein 3 | 1,25879199 | 3,102924535 | 0,013774857 | WT - MAGE-D1_KO |
| **CBLL1** | Q75N03 | E3 ubiquitin-protein ligase Hakai | 1,315642748 | 3,086912257 | 0,014171089 | WT - MAGE-D1_KO |
| **URB1** | O60287 | Nucleolar pre-ribosomal-associated protein 1 | 1,033640748 | 3,065049188 | 0,014453622 | WT - MAGE-D1_KO |
| **INTS5** | Q6P9B9 | Integrator complex subunit 5 | 1,080208102 | 3,031460241 | 0,014867142 | WT - MAGE-D1_KO |
| **TRA2B** | P62995 | Transformer-2 protein homolog beta | 1,157450159 | 2,931729618 | 0,01707418 | WT - MAGE-D1_KO |
| **OTUD7B** | Q6GQQ9 | OTU domain-containing protein 7B | 1,315803201 | 2,785062553 | 0,020415645 | WT - MAGE-D1_KO |
| **PRPF38A** | Q8NAV1 | Pre-mRNA-splicing factor 38A | 1,036579998 | 2,662281 | 0,023562498 | WT - MAGE-D1_KO |
| **DKK 2,00** | Q9UBU2 | Dickkopf-related protein 2 | 1,444847704 | 2,472708439 | 0,029394983 | WT - MAGE-D1_KO |
| **APOH** | P17690 | Beta-2-glycoprotein 1 | 1,0273054 | 2,342888943 | 0,033831685 | WT - MAGE-D1_KO |
| **PHIP** | Q8WWQ0 | PH-interacting protein | 1,338014781 | 2,323637443 | 0,03437227 | WT - MAGE-D1_KO |
| **CELF6** | Q96J87 | CUGBP Elav-like family member 6 | 1,194281313 | 2,15506903 | 0,042531903 | WT - MAGE-D1_KO |
| **AMBP** | P00978 | Protein AMBP | 1,186990871 | 2,095364137 | 0,046066304 | WT - MAGE-D1_KO |
| **FBXO11** | Q86XK2 | F-box only protein 11 | 1,355327252 | 1,52467908 | 0,095802974 | WT - MAGE-D1_KO |
| **HMG20B** | Q9P0W2 | SWI/SNF-related matrix-associated actin-dependent regulator of chromatin subfamily E member 1-related | 1,030236707 | 1,353209944 | 0,122458708 | WT - MAGE-D1_KO |
| **FABP3** | P05413 | Fatty acid-binding protein, heart | -2,028719382 | 5,984379938 | 0.0021401816051564 | WT - MAGE-D1_KO |
| **NOS1** | P29475 | Nitric oxide synthase, brain | -1,770291083 | 5,803133584 | 0.0021401816051564 | WT - MAGE-D1_KO |
| **ASS1** | P00966 | Argininosuccinate synthase | -1,468380148 | 4,941879996 | 0.00280219757181476 | WT - MAGE-D1_KO |
| **ZNF532** | Q9HCE3 | Zinc finger protein 532 | -1,424223349 | 4,826968525 | 0.00314265056036175 | WT - MAGE-D1_KO |
| **GULP1** | Q9UBP9 | PTB domain-containing engulfment adapter protein 1 | -1,037062364 | 4,773288946 | 0.00314265056036175 | WT - MAGE-D1_KO |
| **CARD9** | Q9H257 | Caspase recruitment domain-containing protein 9 | -1,046303267 | 4,689362047 | 0.00348060074904399 | WT - MAGE-D1_KO |
| **PDZRN3** | Q9UPQ7 | E3 ubiquitin-protein ligase PDZRN3 | -1,23605104 | 4,675218192 | 0.00348865146401235 | WT - MAGE-D1_KO |
| **ALDH1L2** | Q3SY69 | Mitochondrial 10-formyltetrahydrofolate dehydrogenase | -1,545365649 | 4,645739053 | 0.00348865146401235 | WT - MAGE-D1_KO |
| **MT-CO2** | P00403 | Cytochrome c oxidase subunit 2 | -1,130429742 | 4,330327311 | 0.00385949998765663 | WT - MAGE-D1_KO |
| **ARHGAP36** | Q6ZRI8 | Rho GTPase-activating protein 36 | -1,463351731 | 4,101306654 | 0.00452917786263547 | WT - MAGE-D1_KO |
| **NKTR** | P30414 | NK-tumor recognition protein | -1,222111782 | 3,992970637 | 0.00506340898618628 | WT - MAGE-D1_KO |
| **SYTL4** | Q96C24 | Synaptotagmin-like protein 4 | -1,127164889 | 3,978120065 | 0.00507452002737587 | WT - MAGE-D1_KO |
| **MAGEA10** | P43363 | Melanoma-associated antigen 10 | -1,799743791 | 3,784675807 | 0.00666294152515313 | WT - MAGE-D1_KO |
| **NDUFA4** | O00483 | Cytochrome c oxidase subunit NDUFA4 | -1,503388324 | 3,736692013 | 0.00689348983483195 | WT - MAGE-D1_KO |
| **FTL** | P02792 | Ferritin light chain | -1,078004086 | 3,625218353 | 0.0077257838674718 | WT - MAGE-D1_KO |
| **CPVL** | Q9H3G5 | Probable serine carboxypeptidase CPVL | -1,412604802 | 3,560121977 | 0.00831190827602605 | WT - MAGE-D1_KO |
| **ZMYM1** | Q5SVZ6 | Zinc finger MYM-type protein 1 | -1,200914949 | 3,247832183 | 0.0117946365207255 | WT - MAGE-D1_KO |
| **PNPLA4** | P41247 | Patatin-like phospholipase domain-containing protein 4 | -1,77063239 | 2,606549299 | 0.0253142796740076 | WT - MAGE-D1_KO |
| **NEBL** | O76041 | Nebulette | -1,011293659 | 2,305815965 | 0.0353607908447414 | WT - MAGE-D1_KO |
| **JPH4** | Q96JJ6 | Junctophilin-4 | -1,000755835 | 2,207292651 | 0.0402764367971766 | WT - MAGE-D1_KO |
| **ECD** | O95905 | Protein ecdysoneless homolog | -1,157705359 | 1,950823834 | 0,055077838 | WT - MAGE-D1_KO |
| **HJURP** | Q8NCD3 | Holliday junction recognition protein | -1,170191079 | 1,334165552 | 0,125827166 | WT - MAGE-D1_KO |
| **TWNK** | Q96RR1 | Twinkle mtDNA helicase | -1,08416095 | 1,194451641 | 0,154913778 | WT - MAGE-D1_KO |

532

**Supplementary table 1:** Proteomics with reported hits and candidates of total proteome of differentiated SH-SY5Y WT versus MAGE-D1 KO cells. Related to Figure 1A.

| Gene | Protein.ID | Protein.Description | log2(FC) | pvalue | -log10pvalue | fdr | Comparison |
|---|---|---|---|---|---|---|---|
| **BASP1** | P80723\|P80723-2 | Brain acid soluble protein 1 OS=Homo sapiens GN=BASP1 PE=1 SV=2\|Isoform 2 of Brain acid solub | 3,23160575 | 2,94E-06 | 5,53162166 | 0,0019788 | WT - MAGE-D1_KO |
| **MAGE-D1** | Q9Y5V3\|Q9Y5V3-2 | Melanoma-associated antigen D1 OS=Homo sapiens GN=MAGED1 PE=1 SV=3\|Isoform 2 of Melano | 2,91164295 | 7,50E-06 | 5,12495813 | 0,0025236 | WT - MAGE-D1_KO |
| **DBH** | P09172 | Dopamine beta-hydroxylase OS=Homo sapiens GN=DBH PE=1 SV=3 | 1,51494214 | 0,000131 | 3,88262016 | 0,0181542 | WT - MAGE-D1_KO |
| **ALCAM** | F5GXJ9\|Q13740\|Q13740-2 | CD166 antigen OS=Homo sapiens GN=ALCAM PE=1 SV=1\|CD166 antigen OS=Homo sapiens GN= | 1,82684657 | 0,0001349 | 3,87006843 | 0,0181542 | WT - MAGE-D1_KO |
| **GNAS** | P63092\|P63092-2\|P63092-3\|P63092-4\|Q5JWF2\|Q5J\ | Guanine nucleotide-binding protein G(s) subunit alpha isoforms short OS=Homo sapiens GN=GNAS | 1,46117826 | 0,0002031 | 3,69238745 | 0,0227759 | WT - MAGE-D1_KO |
| **RTL1** | A6NKG5 | Retrotransposon-like protein 1 OS=Homo sapiens GN=RTL1 PE=3 SV=3 | 1,23429739 | 0,0003338 | 3,47653451 | 0,0320909 | WT - MAGE-D1_KO |
| **HK2** | E9PB90\|P52789 | Hexokinase-2 OS=Homo sapiens GN=HK2 PE=1 SV=1\|Hexokinase-2 OS=Homo sapiens GN=HK2 F | 1,18363785 | 0,0004388 | 3,35769462 | 0,0328154 | WT - MAGE-D1_KO |
| **IGSF3** | O75054\|O75054-2 | Immunoglobulin superfamily member 3 OS=Homo sapiens GN=IGSF3 PE=2 SV=3\|Isoform 2 of Immu | 1,08313447 | 0,0021233 | 2,67298784 | 0,0752097 | WT - MAGE-D1_KO |
| **LBR** | Q14739 | Lamin-B receptor OS=Homo sapiens GN=LBR PE=1 SV=2 | 1,13561961 | 0,0085168 | 2,06972425 | 0,1364714 | WT - MAGE-D1_KO |
| **APOB** | A0A087WTM7\|P04114 | Apolipoprotein B-100 OS=Homo sapiens GN=APOB PE=1 SV=1\|Apolipoprotein B-100 OS=Homo sa | -1,4999505 | 6,63E-05 | 4,17871334 | 0,0148655 | WT - MAGE-D1_KO |
| **GULP1** | Q9UBP9\|Q9UBP9-3\|Q9UBP9-4 | PTB domain-containing engulfment adapter protein 1 OS=Homo sapiens GN=GULP1 PE=1 SV=1\|Iso | -1,4561652 | 0,0004301 | 3,36643463 | 0,0328154 | WT - MAGE-D1_KO |
| **MT-CO2** | P00403 | Cytochrome c oxidase subunit 2 OS=Homo sapiens GN=MT-CO2 PE=1 SV=1 | -1,223913 | 0,0005537 | 3,25675995 | 0,0372611 | WT - MAGE-D1_KO |
| **COX7C** | D6R9Z7\|P15954 | Cytochrome c oxidase subunit 7C, mitochondrial OS=Homo sapiens GN=COX7C PE=1 SV=1\|Cytoch | -1,0321246 | 0,000735 | 3,13369102 | 0,0438075 | WT - MAGE-D1_KO |
| **LGALS3BP** | Q08380 | Galectin-3-binding protein OS=Homo sapiens GN=LGALS3BP PE=1 SV=1 | -1,7836725 | 0,0007811 | 3,10728562 | 0,0438075 | WT - MAGE-D1_KO |
| **COX4I1** | P13073 | Cytochrome c oxidase subunit 4 isoform 1, mitochondrial OS=Homo sapiens GN=COX4I1 PE=1 SV=' | -1,0421385 | 0,0013934 | 2,85593375 | 0,055161 | WT - MAGE-D1_KO |
| **CACNA2D'** | P54289\|P54289-2\|P54289-3\|P54289-4\|P54289-5 | Voltage-dependent calcium channel subunit alpha-2/delta-1 OS=Homo sapiens GN=CACNA2D1 PE | -1,0346605 | 0,0038768 | 2,41152216 | 0,1003505 | WT - MAGE-D1_KO |
| **LMNA** | P02545\|P02545-2\|P02545-3\|P02545-6\|Q3BDU5 | Prelamin-A/C OS=Homo sapiens GN=LMNA PE=1 SV=1\|Isoform C of Prelamin-A/C OS=Homo sapie | -1,2772409 | 0,0043335 | 2,36316123 | 0,1041588 | WT - MAGE-D1_KO |
| **RPS8** | P62241\|Q5JR95 | 40S ribosomal protein S8 OS=Homo sapiens GN=RPS8 PE=1 SV=2\|40S ribosomal protein S8 OS=H | -1,3186352 | 0,013352 | 1,87445273 | 0,1523037 | WT - MAGE-D1_KO |

**Supplementary table 2:** Proteomics with reported hits and candidates of endogenous MAGE-D1 immunoprecipitation from undifferentiated SH-SY5Y WT versus MAGE-D1 KO cells. Related to Figure 1B.

| Gene | Protein.ID | Protein.Description | log2(FC) | pvalue | -log10pvalue | fdr | Comparison |
|---|---|---|---|---|---|---|---|
| **MAGED2** | Q9UNF1 | Melanoma-associated antigen D2 | 2,949743422 | 2,79E-06 | 5,554577898 | 0,005786823 | WT - MAGE-D1_KO |
| **PJA2** | O43164 | E3 ubiquitin-protein ligase Praja-2 | 2,74841575 | 1,16E-05 | 4,935342991 | 0,008549467 | WT - MAGE-D1_KO |
| **MAGED1** | Q9Y5V3 | Melanoma-associated antigen D1 | 3,550960084 | 1,43E-05 | 4,845716478 | 0,008549467 | WT - MAGE-D1_KO |
| **SVIL** | O95425 | Supervillin | 1,587491408 | 1,65E-05 | 4,783019047 | 0,008549467 | WT - MAGE-D1_KO |
| **PJA1** | Q8NG27 | E3 ubiquitin-protein ligase Praja-1 | 2,555463454 | 2,12E-05 | 4,67428403 | 0,008785451 | WT - MAGE-D1_KO |
| **USP7** | Q93009 | Ubiquitin carboxyl-terminal hydrolase 7 | 1,52749904 | 2,78E-05 | 4,556469945 | 0,009602778 | WT - MAGE-D1_KO |
| **SMC5** | Q8IY18 | Structural maintenance of chromosomes protein 5 | 1,525915374 | 4,40E-05 | 4,356776816 | 0,013035967 | WT - MAGE-D1_KO |
| **GSN** | P06396 | Gelsolin | 1,382061417 | 5,79E-05 | 4,237347996 | 0,015016894 | WT - MAGE-D1_KO |
| **ASPM** | Q8IZT6 | Abnormal spindle-like microcephaly-associated protein | 1,278757208 | 9,22E-05 | 4,035305004 | 0,020749394 | WT - MAGE-D1_KO |
| **SLC25A13** | Q9UJS0 | Electrogenic aspartate/glutamate antiporter SLC25A13, mitochondrial | 1,096804114 | 0,000115808 | 3,936262467 | 0,021845548 | WT - MAGE-D1_KO |
| **SMTN** | P53814 | Smoothelin | 1,478995982 | 0,00013608 | 3,866206324 | 0,023530466 | WT - MAGE-D1_KO |
| **SLC25A12** | O75746 | Electrogenic aspartate/glutamate antiporter SLC25A12, mitochondrial | 1,101481991 | 0,000170692 | 3,767786135 | 0,023650416 | WT - MAGE-D1_KO |
| **MYO5A** | Q9Y4I1 | Unconventional myosin-Va | 1,062613368 | 0,000173926 | 3,759635132 | 0,023650416 | WT - MAGE-D1_KO |
| **AP2B1** | P63010 | AP-2 complex subunit beta | 1,395592568 | 0,0001874 | 3,727229868 | 0,023650416 | WT - MAGE-D1_KO |
| **CLINT1** | Q14677 | Clathrin interactor 1 | 1,006600852 | 0,000198457 | 3,702334301 | 0,023650416 | WT - MAGE-D1_KO |
| **TPM4** | P67936 | Tropomyosin alpha-4 chain | 1,63096714 | 0,000235803 | 3,627451389 | 0,023650416 | WT - MAGE-D1_KO |
| **NSMCE3** | Q96MG7 | Non-structural maintenance of chromosomes element 3 homolog | 1,07840245 | 0,000238115 | 3,62321363 | 0,023650416 | WT - MAGE-D1_KO |
| **MYH9** | P35579 | Myosin-9 | 2,290686298 | 0,000239354 | 3,620960023 | 0,023650416 | WT - MAGE-D1_KO |
| **AKAP8L** | Q9ULX6 | A-kinase anchor protein 8-like | 1,068628436 | 0,000295915 | 3,528832354 | 0,025584357 | WT - MAGE-D1_KO |
| **RPL22L1** | Q6P5R6 | 60S ribosomal protein L22-like 1 | 1,225039124 | 0,00032355 | 3,49005847 | 0,026283114 | WT - MAGE-D1_KO |
| **AP2M1** | Q96CW1 | AP-2 complex subunit mu | 1,13108493 | 0,000346382 | 3,460444151 | 0,026283114 | WT - MAGE-D1_KO |
| **SPECC1L** | Q69YQ0 | Cytospin-A | 1,019920968 | 0,000354664 | 3,450183246 | 0,026283114 | WT - MAGE-D1_KO |
| **P4HA2** | O15460 | Prolyl 4-hydroxylase subunit alpha-2 | 1,007796651 | 0,000378725 | 3,421675508 | 0,026990941 | WT - MAGE-D1_KO |
| **HSPB1** | P04792 | Heat shock protein beta-1 | 1,008352885 | 0,000472833 | 3,325292423 | 0,029251124 | WT - MAGE-D1_KO |
| **TPM3** | P06753 | Tropomyosin alpha-3 chain | 1,962155385 | 0,000518421 | 3,285317015 | 0,029251124 | WT - MAGE-D1_KO |
| **METTL7A** | Q9H8H3 | Putative methyltransferase-like protein 7A | 1,075460728 | 0,000533072 | 3,27321412 | 0,029251124 | WT - MAGE-D1_KO |
| **MYL6** | P60660 | Myosin light polypeptide 6 | 1,583981617 | 0,00075668 | 3,121087815 | 0,029251124 | WT - MAGE-D1_KO |
| **TPM1** | P09493 | Tropomyosin alpha-1 chain | 2,083741139 | 0,000759321 | 3,119574678 | 0,029251124 | WT - MAGE-D1_KO |
| **MYH14** | Q7Z406 | Myosin-14 | 1,978135671 | 0,000761234 | 3,118481788 | 0,029251124 | WT - MAGE-D1_KO |
| **MYH10** | P35580 | Myosin-10 | 1,775843863 | 0,000818907 | 3,086765212 | 0,030786514 | WT - MAGE-D1_KO |
| **PPP1R12A** | O14974 | Protein phosphatase 1 regulatory subunit 12A | 1,017542199 | 0,00092586 | 3,033454634 | 0,032245012 | WT - MAGE-D1_KO |
| **AP1B1** | Q10567 | AP-1 complex subunit beta-1 | 1,041797847 | 0,000963148 | 3,01630718 | 0,032762806 | WT - MAGE-D1_KO |
| **AP2S1** | P53680 | AP-2 complex subunit sigma | 1,046034391 | 0,001114356 | 2,952976226 | 0,035318354 | WT - MAGE-D1_KO |
| **OBI1** | Q5W0B1 | ORC ubiquitin ligase 1 | 1,193718097 | 0,001273758 | 2,894913218 | 0,035763287 | WT - MAGE-D1_KO |
| **RAB3GAP2** | Q9H2M9 | Rab3 GTPase-activating protein non-catalytic subunit | 1,39773751 | 0,001275414 | 2,894348956 | 0,035763287 | WT - MAGE-D1_KO |
| **MYO1E** | Q12965 | Unconventional myosin-Ie | 1,170156851 | 0,001369093 | 2,863567024 | 0,037878242 | WT - MAGE-D1_KO |
| **UACA** | Q9BZF9 | Uveal autoantigen with coiled-coil domains and ankyrin repeats | 1,006344431 | 0,002134542 | 2,670695262 | 0,044762461 | WT - MAGE-D1_KO |
| **TTK** | P33981 | Dual specificity protein kinase TTK | 1,004790949 | 0,002259481 | 2,6459914 | 0,045845145 | WT - MAGE-D1_KO |
| **ARPC2** | O15144 | Actin-related protein 2/3 complex subunit 2 | 1,014671036 | 0,004789602 | 2,319700586 | 0,070306271 | WT - MAGE-D1_KO |
| **DBN1** | Q16643 | Drebrin | 1,308767477 | 0,005395771 | 2,267946504 | 0,07177067 | WT - MAGE-D1_KO |
| **LIMA1** | Q9UHB6 | LIM domain and actin-binding protein 1 | 1,03628758 | 0,009136103 | 2,039238997 | 0,090309621 | WT - MAGE-D1_KO |
| **BASP1** | P80723 | Brain acid soluble protein 1 | 1,313230811 | 0,010911128 | 1,962130354 | 0,096871866 | WT - MAGE-D1_KO |
| **WDR91** | A4D1P6 | WD repeat-containing protein 91 | -1,134901999 | 1,00E-04 | 4,000012689 | 0,020749394 | WT - MAGE-D1_KO |
| **CEP250** | Q9BV73 | Centrosome-associated protein CEP250 | -1,12474811 | 0,00051358 | 3,289392183 | 0,029251124 | WT - MAGE-D1_KO |
| **HMBOX1** | Q6NT76 | Homeobox-containing protein 1 | -1,116349243 | 0,000550901 | 3,258926711 | 0,029251124 | WT - MAGE-D1_KO |
| **RBM20** | Q5T481 | RNA-binding protein 20 | -1,06492685 | 0,000675273 | 3,170520615 | 0,029251124 | WT - MAGE-D1_KO |
| **SNX33** | Q8WV41 | Sorting nexin-33 | -1,256822271 | 0,001217315 | 2,914597194 | 0,035763287 | WT - MAGE-D1_KO |
| **NEFL** | P07196 | Neurofilament light polypeptide | -1,38469432 | 0,002275687 | 2,642887526 | 0,045845145 | WT - MAGE-D1_KO |
| **HSPG2** | P98160 | Basement membrane-specific heparan sulfate proteoglycan core protein | -1,089885531 | 0,002689557 | 2,570319235 | 0,051101346 | WT - MAGE-D1_KO |
| **NEFM** | P07197 | Neurofilament medium polypeptide | -1,490679709 | 0,004073451 | 2,39003752 | 0,064979929 | WT - MAGE-D1_KO |
| **PHC1** | P78364 | Polyhomeotic-like protein 1 | -1,313863681 | 0,004960175 | 2,304503008 | 0,070439771 | WT - MAGE-D1_KO |
| **TP53** | P04637 | Cellular tumor antigen p53 | -2,012754931 | 0,006304888 | 2,200322606 | 0,078050756 | WT - MAGE-D1_KO |
| **CYP20A1** | Q6UW02 | Cytochrome P450 20A1 | -1,068658987 | 0,007227573 | 2,141007512 | 0,079772415 | WT - MAGE-D1_KO |
| **VIM** | P08670 | Vimentin | -1,453057835 | 0,008734523 | 2,058760783 | 0,089472762 | WT - MAGE-D1_KO |

**Supplementary table 3:** Proteomics with reported hits and candidates of endogenous MAGE-D1 immunoprecipitation from HEK WT versus MAGE-D1 KO cells. Related to Supplementary Figure 1A.

| | | | D4 | | | | | | D11 | | | |
|---|---|---|---|---|---|---|---|---|---|---|---|---|
| Vehicle | 93,3 | 76,3 | 100,1 | 240 | 253,5 | 314,1 | 108,7 | 101,4 | 142 | 287,9 | 247,2 | 436,7 |
| P5091 | *284,5** | 66,2 | 156,3 | 79,8 | 103,4 | 68,5 | *254,1** | 80,8 | 197,3 | 83,4 | 127,9 | 70,8 |
| **Two way anova** | | | | | | | | | | | | |
| Table Analyzed | D4/D11 Coke only | | | | | | | | | | | |
| **Two-way RM ANOVA** | Matching: Across row | | | | | | | | | | | |
| Assume sphericity? | Yes | | | | | | | | | | | |
| Alpha | 0,05 | | | | | | | | | | | |
| **Source of Variation** | **% of total variation** | **P value** | **P value summary** | **Significant?** | | | | | | | | |
| Treatment x Days | 0,3712 | 0,2857 | ns | No | | | | | | | | |
| Treatment | 24,28 | 0,1118 | ns | No | | | | | | | | |
| Days | 2,209 | 0,0218 | * | Yes | | | | | | | | |
| Subject | 70,36 | <0,0001 | **** | Yes | | | | | | | | |
| **ANOVA table** | **SS** | **DF** | **MS** | **F (DFn, DFd)** | **P value** | | | | | | | |
| Treatment x Days | 778,9 | 1 | 778,9 | F (1, 9) = 1,288 | P=0,2857 | | | | | | | |
| Treatment | 50963 | 1 | 50963 | F (1, 9) = 3,106 | P=0,1118 | | | | | | | |
| Days | 4635 | 1 | 4635 | F (1, 9) = 7,664 | P=0,0218 | | | | | | | |
| Subject | 147648 | 9 | 16405 | F (9, 9) = 27,13 | P<0,0001 | | | | | | | |
| Residual | 5443 | 9 | 604,8 | | | | | | | | | |
| **Difference between row means** | | | | | | | | | | | | |
| Mean of Vehicle | 200,1 | | | | | | | | | | | |
| Mean of P5091 | 103,4 | | | | | | | | | | | |
| Difference between means | 96,66 | | | | | | | | | | | |
| SE of difference | 54,84 | | | | | | | | | | | |
| 95% CI of difference | -27,40 to 220,7 | | | | | | | | | | | |
| **Difference between column means** | | | | | | | | | | | | |
| Mean of D4 | 137,2 | | | | | | | | | | | |
| Mean of D11 | 166,3 | | | | | | | | | | | |
| Difference between means | -29,15 | | | | | | | | | | | |
| SE of difference | 10,53 | | | | | | | | | | | |
| 95% CI of difference | -52,97 to -5,330 | | | | | | | | | | | |
| **Interaction CI** | | | | | | | | | | | | |
| Mean diff, A1 - B1 | -41,1 | | | | | | | | | | | |
| Mean diff, A2 - B2 | -17,2 | | | | | | | | | | | |
| (A1 -B1) - (A2 - B2) | -23,9 | | | | | | | | | | | |
| 95% CI of difference | -71,54 to 23,74 | | | | | | | | | | | |
| (B1 - A1) - (B2 - A2) | 23,9 | | | | | | | | | | | |
| 95% CI of difference | -23,74 to 71,54 | | | | | | | | | | | |
| **Data summary** | | | | | | | | | | | | |
| Number of columns (Days) | 2 | | | | | | | | | | | |
| Number of rows (Treatment) | 2 | | | | | | | | | | | |
| Number of subjects (Subject) | 11 | | | | | | | | | | | |
| Number of missing values | 0 | | | | | | | | | | | |
| **Two way anova multicomparison** | | | | | | | | | | | | |
| Compare each cell mean with the other cell mean in that row | | | | | | | | | | | | |
| Number of families | 1 | | | | | | | | | | | |
| Number of comparisons per family | 2 | | | | | | | | | | | |
| Alpha | 0,05 | | | | | | | | | | | |
| Šídák's multiple comparisons test | Predicted (LS) Mean diff, | 95,00% CI of diff, | Below threshold? | Summary | Adjusted P Value | | | | | | | |
| D4 - D11 | | | | | | | | | | | | |
| Vehicle | -41,1 | -79,11 to -3,088 | Yes | * | 0,0352 | | | | | | | |
| P5091 | -17,2 | -58,84 to 24,44 | No | ns | 0,5064 | | | | | | | |
| Test details | Predicted (LS) Mean 1 | Predicted (LS) Mean 2 | Predicted (LS) Mean diff, | SE of diff, | N1 | N2 | t | DF | | | | |
| D4 - D11 | | | | | | | | | | | | |
| Vehicle | 179,6 | 220,7 | -41,1 | 14,2 | 6 | 6 | 2,895 | 9 | | | | |
| P5091 | 94,84 | 112 | -17,2 | 15,55 | 5 | 5 | 1,106 | 9 | | | | |

**Supplementary table 4:** Statistical test of cocaine induced locomotor sensitization in mice. Related to Supplementary Figure 2A-B.

| Gene | Protein ID | Protein Description | log2(FC) | pvalue | -log10(pvalue) | fdr | Comparison |
|---|---|---|---|---|---|---|---|
| **DPP6** | P42658 | Dipeptidyl aminopeptidase-like protein 6 | 6,507314974 | 1,98291E-05 | 4,70269697 | 0,003540596 | WT -MAGE-D1 KO |
| **GRM8** | O00222 | Metabotropic glutamate receptor 8 | 4,308807288 | 7,52894E-06 | 5,123266361 | 0,002717318 | WT -MAGE-D1 KO |
| **DBH** | P09172 | Dopamine beta-hydroxylase | 3,803250593 | 8,45463E-06 | 5,072905368 | 0,002717318 | WT -MAGE-D1 KO |
| **DPP10** | Q8N608 | Inactive dipeptidyl peptidase 10 | 3,765840252 | 7,93496E-06 | 5,10045534 | 0,002717318 | WT -MAGE-D1 KO |
| **ELFN1** | P0C7U0 | Protein ELFN1 | 3,33215605 | 4,98633E-06 | 5,302218594 | 0,002717318 | WT -MAGE-D1 KO |
| **ACHE** | P22303 | Acetylcholinesterase | 3,24868838 | 2,43619E-05 | 4,613288725 | 0,003793784 | WT -MAGE-D1 KO |
| **CADM3** | Q8N126 | Cell adhesion molecule 3 | 2,933936532 | 6,3705E-05 | 4,195826544 | 0,005849938 | WT -MAGE-D1 KO |
| **ACVRL1** | P37023 | Serine/threonine-protein kinase receptor R3 | 2,905235063 | 1,11122E-05 | 4,954201767 | 0,003246769 | WT -MAGE-D1 KO |
| **CNTNAP5** | Q8WYK1 | Contactin-associated protein-like 5 | 2,790138725 | 1,62283E-05 | 4,789727711 | 0,003540596 | WT -MAGE-D1 KO |
| **CNTN2** | Q02246 | Contactin-2 | 2,686752349 | 3,58885E-05 | 4,445045224 | 0,00384485 | WT -MAGE-D1 KO |
| **SDK1** | Q7Z5N4 | Protein sidekick-1 | 2,61898094 | 2,68477E-05 | 4,571092653 | 0,003793784 | WT -MAGE-D1 KO |
| **NRXN1** | Q9ULB1 | Neurexin-1 | 2,599811763 | 1,82165E-05 | 4,739535906 | 0,003540596 | WT -MAGE-D1 KO |
| **GRM7** | Q14831 | Metabotropic glutamate receptor 7 | 2,59377037 | 5,09255E-05 | 4,293065121 | 0,005114825 | WT -MAGE-D1 KO |
| **PRNP** | P04156 | Major prion protein | 2,562744781 | 9,99405E-05 | 4,000258407 | 0,0069828 | WT -MAGE-D1 KO |
| **PDGFRA** | P16234 | Platelet-derived growth factor receptor alpha | 2,459406598 | 2,91424E-05 | 4,535474244 | 0,003793784 | WT -MAGE-D1 KO |
| **FAT1** | Q14517 | Protocadherin Fat 1 | 2,416037073 | 3,26775E-05 | 4,485751322 | 0,00384485 | WT -MAGE-D1 KO |
| **CELSR3** | Q9NYQ7 | Cadherin EGF LAG seven-pass G-type receptor 3 | 2,264767871 | 3,48641E-05 | 4,45762191 | 0,00384485 | WT -MAGE-D1 KO |
| **PLXNA2** | O75051 | Plexin-A2 | 2,258623288 | 2,82236E-05 | 4,549387244 | 0,003793784 | WT -MAGE-D1 KO |
| **LRFN2** | Q9ULH4 | Leucine-rich repeat and fibronectin type-III domain- | 2,234584698 | 0,000422706 | 3,373961722 | 0,012696978 | WT -MAGE-D1 KO |
| **ITM2C** | Q9NQX7 | Integral membrane protein 2C | 2,173673073 | 0,001105405 | 2,95647871 | 0,021531943 | WT -MAGE-D1 KO |
| **BASP1** | P80723 | Brain acid soluble protein 1 | 2,081934741 | 0,000129303 | 3,888391895 | 0,00784112 | WT -MAGE-D1 KO |
| **ITGA7** | Q13683 | Integrin alpha-7 | 2,071234164 | 0,000143859 | 3,842063414 | 0,00825647 | WT -MAGE-D1 KO |
| **C7** | P10643 | Complement component C7 | 2,037348927 | 0,000300338 | 3,52238963 | 0,011078435 | WT -MAGE-D1 KO |
| **TENM4** | Q6N022 | Teneurin-4 | 2,034493075 | 0,000623543 | 3,205133598 | 0,015068174 | WT -MAGE-D1 KO |
| **MSRB2** | Q9Y3D2 | Methionine-R-sulfoxide reductase B2, mitochondria | 2,019874417 | 0,000112746 | 3,947898174 | 0,0075254 | WT -MAGE-D1 KO |
| **IGSF3** | O75054 | Immunoglobulin superfamily member 3 | 1,970066632 | 7,97535E-05 | 4,098250114 | 0,006572508 | WT -MAGE-D1 KO |
| **RTL1** | A6NKG5 | Retrotransposon-like protein 1 | 1,949263252 | 0,001058856 | 2,975163036 | 0,020750998 | WT -MAGE-D1 KO |
| **PLG** | P00747 | Plasminogen | 1,934323462 | 0,000764003 | 3,116905197 | 0,016849564 | WT -MAGE-D1 KO |
| **SEMA4B** | Q9NPR2 | Semaphorin-4B | 1,91858235 | 0,001496631 | 2,824885204 | 0,02642952 | WT -MAGE-D1 KO |
| **GPC2** | Q8N158 | Glypican-2 | 1,897707126 | 0,000115992 | 3,935572839 | 0,0075254 | WT -MAGE-D1 KO |
| **NRCAM** | Q92823 | Neuronal cell adhesion molecule | 1,894441574 | 7,83674E-05 | 4,105864784 | 0,006572508 | WT -MAGE-D1 KO |
| **TNR** | Q92752 | Tenascin-R | 1,890122644 | 0,000110688 | 3,955899782 | 0,0075254 | WT -MAGE-D1 KO |
| **NRXN2** | Q9P2S2 | Neurexin-2 | 1,863577072 | 8,76934E-05 | 4,05703294 | 0,006823157 | WT -MAGE-D1 KO |
| **ISLR2** | Q6UXK2 | Immunoglobulin superfamily containing leucine-rich | 1,848858948 | 8,47325E-05 | 4,071950059 | 0,006808255 | WT -MAGE-D1 KO |
| **ALK** | Q9UM73 | ALK tyrosine kinase receptor | 1,832854283 | 0,000211827 | 3,674017937 | 0,009725902 | WT -MAGE-D1 KO |
| **SEMA4C** | Q9C0C4 | Semaphorin-4C | 1,828612412 | 9,58527E-05 | 4,018395853 | 0,00684601 | WT -MAGE-D1 KO |
| **CACNG4** | Q9UBN1 | Voltage-dependent calcium channel gamma-4 subu | 1,800687316 | 0,000469621 | 3,328252321 | 0,013240022 | WT -MAGE-D1 KO |
| **RAB3A** | P20336 | Ras-related protein Rab-3A | 1,763627962 | 0,000367965 | 3,434193548 | 0,012067748 | WT -MAGE-D1 KO |
| **XPNPEP2** | O43895 | Xaa-Pro aminopeptidase 2 | 1,73583882 | 0,001280776 | 2,892526824 | 0,024072597 | WT -MAGE-D1 KO |
| **ITGA9** | Q13797 | Integrin alpha-9 | 1,715899575 | 0,000283493 | 3,547457303 | 0,010594735 | WT -MAGE-D1 KO |
| **SEMA7A** | O75326 | Semaphorin-7A | 1,709587367 | 0,000187071 | 3,727993988 | 0,008961411 | WT -MAGE-D1 KO |
| **SEPTIN6** | Q14141 | Septin-6 | 1,699129519 | 0,001524512 | 2,816869124 | 0,026576581 | WT -MAGE-D1 KO |
| **IGF2R** | P11717 | Cation-independent mannose-6-phosphate recepto | 1,682289631 | 0,000140886 | 3,851132201 | 0,00825647 | WT -MAGE-D1 KO |
| **PTCH1** | Q13635 | Protein patched homolog 1 | 1,670603577 | 0,000775898 | 3,110195485 | 0,016849564 | WT -MAGE-D1 KO |
| **SEMA4D** | Q92854 | Semaphorin-4D | 1,619125042 | 0,001553714 | 2,808628989 | 0,026847505 | WT -MAGE-D1 KO |
| **PCLAF** | Q15004 | PCNA-associated factor | 1,587853083 | 0,00044936 | 3,347405983 | 0,01312947 | WT -MAGE-D1 KO |
| **LRP4** | O75096 | Low-density lipoprotein receptor-related protein 4 | 1,58641473 | 0,000879694 | 3,055668443 | 0,018240878 | WT -MAGE-D1 KO |
| **CHL1** | O00533 | Neural cell adhesion molecule L1-like protein | 1,580931557 | 0,000408548 | 3,388756733 | 0,012696978 | WT -MAGE-D1 KO |
| **LIMD2** | Q9BT23 | LIM domain-containing protein 2 | 1,567527672 | 0,000315205 | 3,501406802 | 0,011086327 | WT -MAGE-D1 KO |
| **L1CAM** | P32004 | Neural cell adhesion molecule L1 | 1,565150452 | 0,00030839 | 3,510899439 | 0,011086327 | WT -MAGE-D1 KO |
| **SEZ6** | Q53EL9 | Seizure protein 6 homolog | 1,552806041 | 0,000486682 | 3,31275494 | 0,013601697 | WT -MAGE-D1 KO |
| **IGSF9** | Q9P2J2 | Protein turtle homolog A | 1,54774504 | 0,000411749 | 3,385367321 | 0,012696978 | WT -MAGE-D1 KO |
| **INSR** | P06213 | Insulin receptor | 1,543398225 | 0,000252145 | 3,598348908 | 0,01004851 | WT -MAGE-D1 KO |
| **ECEL1** | O95672 | Endothelin-converting enzyme-like 1 | 1,531509048 | 0,000328447 | 3,483534423 | 0,011350853 | WT -MAGE-D1 KO |
| **KALRN** | O60229 | Kalirin | 1,523103989 | 0,000222682 | 3,652314364 | 0,009804121 | WT -MAGE-D1 KO |
| **SH3PXD2B** | A1X283 | SH3 and PX domain-containing protein 2B | 1,510974799 | 0,002074387 | 2,683110269 | 0,032208111 | WT -MAGE-D1 KO |
| **TENM3** | Q9P273 | Teneurin-3 | 1,499025962 | 0,000461965 | 3,335390553 | 0,013240022 | WT -MAGE-D1 KO |
| **ALCAM** | Q13740 | CD166 antigen | 1,451338156 | 0,000885949 | 3,052591464 | 0,018252813 | WT -MAGE-D1 KO |
| **SLC8A3** | P57103 | Sodium/calcium exchanger 3 | 1,449644737 | 0,00245424 | 2,610083012 | 0,035371868 | WT -MAGE-D1 KO |
| **CNTFR** | P26992 | Ciliary neurotrophic factor receptor subunit alpha | 1,44176182 | 0,000559169 | 3,252457297 | 0,014530042 | WT -MAGE-D1 KO |
| **ISLR** | O14498 | Immunoglobulin superfamily containing leucine-rich | 1,41575793 | 0,000468862 | 3,32895459 | 0,013240022 | WT -MAGE-D1 KO |
| **ADGRA3** | Q8IWK6 | Adhesion G protein-coupled receptor A3 | 1,388000926 | 0,000681989 | 3,166222477 | 0,015999368 | WT -MAGE-D1 KO |
| **B4GALNT1** | Q00973 | Beta-1,4 N-acetylgalactosaminyltransferase 1 | 1,368163862 | 0,001115581 | 2,952499005 | 0,021599255 | WT -MAGE-D1 KO |
| **UNC5A** | Q6ZN44 | Netrin receptor UNC5A | 1,356530024 | 0,00059402 | 3,226198792 | 0,014853467 | WT -MAGE-D1 KO |
| **APP** | P05067 | Amyloid-beta precursor protein | 1,352564344 | 0,000591115 | 3,228328286 | 0,014853467 | WT -MAGE-D1 KO |
| **CHGA** | P10645 | Chromogranin-A | 1,329499013 | 0,001709704 | 2,767078957 | 0,02857227 | WT -MAGE-D1 KO |
| **RAB3D** | O95716 | Ras-related protein Rab-3D | 1,316153206 | 0,000722788 | 3,140988819 | 0,016446668 | WT -MAGE-D1 KO |
| **C11orf96** | Q7Z7L8 | Uncharacterized protein C11orf96 | 1,315183296 | 0,000829704 | 3,081076738 | 0,017543876 | WT -MAGE-D1 KO |
| **SEMA4G** | Q9NTN9 | Semaphorin-4G | 1,313607115 | 0,004567968 | 2,340276948 | 0,047393897 | WT -MAGE-D1 KO |
| **YBX2** | Q9Y2T7 | Y-box-binding protein 2 | 1,309372161 | 0,001574287 | 2,802916073 | 0,027035086 | WT -MAGE-D1 KO |
| **LAMP5** | Q9UJQ1 | Lysosome-associated membrane glycoprotein 5 | 1,305139141 | 0,002662842 | 2,574654577 | 0,03683901 | WT -MAGE-D1 KO |
| **IGF1R** | P08069 | Insulin-like growth factor 1 receptor | 1,299165839 | 0,000523203 | 3,28133017 | 0,014285819 | WT -MAGE-D1 KO |
| **VPS13A** | Q96RL7 | Intermembrane lipid transfer protein VPS13A | 1,283271027 | 0,000759322 | 3,119574032 | 0,016849564 | WT -MAGE-D1 KO |
| **ITGB5** | P18084 | Integrin beta-5 | 1,271176901 | 0,000958534 | 3,018392392 | 0,019254556 | WT -MAGE-D1 KO |
| **SORCS1** | Q8WY21 | VPS10 domain-containing receptor SorCS1 | 1,261079912 | 0,001240365 | 2,906450511 | 0,023450194 | WT -MAGE-D1 KO |
| **DCX** | O43602 | Neuronal migration protein doublecortin | 1,258511625 | 0,001337824 | 2,873600869 | 0,024430499 | WT -MAGE-D1 KO |
| **ZC2HC1A** | Q96GY0 | Zinc finger C2HC domain-containing protein 1A | 1,258216628 | 0,004465389 | 2,350140732 | 0,047054948 | WT -MAGE-D1 KO |
| **PRPH** | P41219 | Peripherin | 1,243173502 | 0,002121179 | 2,673422782 | 0,032310274 | WT -MAGE-D1 KO |
| **KIF1A** | Q12756 | Kinesin-like protein KIF1A | 1,220628412 | 0,001321752 | 2,878849919 | 0,024413953 | WT -MAGE-D1 KO |
| **HDLBP** | Q00341 | Vigilin | 1,210140027 | 0,000912691 | 3,039676365 | 0,018684 | WT -MAGE-D1 KO |
| **IRS2** | Q9Y4H2 | Insulin receptor substrate 2 | 1,205680472 | 0,001711153 | 2,766711195 | 0,02857227 | WT -MAGE-D1 KO |
| **RET** | P07949 | Proto-oncogene tyrosine-protein kinase receptor Re | 1,200198546 | 0,000792 | 3,101274813 | 0,016907222 | WT -MAGE-D1 KO |

| | | | | | | | |
|---|---|---|---|---|---|---|---|
| **SNRNP27** | Q8WVK2 | U4/U6,U5 small nuclear ribonucleoprotein 27 kDa p | 1,200130838 | 0,001483013 | 2,828855162 | 0,026333715 | WT -MAGE-D1 KO |
| **FANCD2** | Q9BXW9 | Fanconi anemia group D2 protein | 1,198294612 | 0,000720876 | 3,142139621 | 0,016446668 | WT -MAGE-D1 KO |
| **UNC5B** | Q8IZJ1 | Netrin receptor UNC5B | 1,196287151 | 0,002308055 | 2,636753859 | 0,034502737 | WT -MAGE-D1 KO |
| **SEPTIN7** | Q16181 | Septin-7 | 1,180252332 | 0,000792229 | 3,101149342 | 0,016907222 | WT -MAGE-D1 KO |
| **SEPTIN2** | Q15019 | Septin-2 | 1,165764036 | 0,001715759 | 2,76554382 | 0,02857227 | WT -MAGE-D1 KO |
| **SHTN1** | A0MZ66 | Shootin-1 | 1,159386615 | 0,00281458 | 2,550586412 | 0,036891657 | WT -MAGE-D1 KO |
| **KNTC1** | P50748 | Kinetochore-associated protein 1 | 1,151691762 | 0,004289407 | 2,367602709 | 0,045649521 | WT -MAGE-D1 KO |
| **STXBP1** | P61764 | Syntaxin-binding protein 1 | 1,123361834 | 0,000977854 | 3,009725992 | 0,019520638 | WT -MAGE-D1 KO |
| **ERI1** | Q8IV48 | 3'-5' exoribonuclease 1 | 1,120470206 | 0,001581393 | 2,800960273 | 0,027035086 | WT -MAGE-D1 KO |
| **NDRG1** | Q92597 | Protein NDRG1 | 1,11412199 | 0,001968972 | 2,705760361 | 0,030865688 | WT -MAGE-D1 KO |
| **ANXA6** | P08133 | Annexin A6 | 1,092633001 | 0,003346093 | 2,475461963 | 0,039978973 | WT -MAGE-D1 KO |
| **CRACD** | Q6ZU35 | Capping protein-inhibiting regulator of actin dynami | 1,076517827 | 0,001314545 | 2,881224524 | 0,024413953 | WT -MAGE-D1 KO |
| **DPYSL5** | Q9BPU6 | Dihydropyrimidinase-related protein 5 | 1,072305234 | 0,002497236 | 2,602540458 | 0,035723293 | WT -MAGE-D1 KO |
| **NUDT3** | O95989 | Diphosphoinositol polyphosphate phosphohydrolas | 1,046600567 | 0,003748625 | 2,426128036 | 0,042581113 | WT -MAGE-D1 KO |
| **FNBP1L** | Q5T0N5 | Formin-binding protein 1-like | 1,045068134 | 0,001398484 | 2,85434239 | 0,024970715 | WT -MAGE-D1 KO |
| **SYNE2** | Q8WXH0 | Nesprin-2 | 1,035604716 | 0,002908068 | 2,536395487 | 0,037088297 | WT -MAGE-D1 KO |
| **LEPR** | P48357 | Leptin receptor | 1,034408457 | 0,001529766 | 2,815375027 | 0,026576581 | WT -MAGE-D1 KO |
| **DCAF7** | P61962 | DDB1- and CUL4-associated factor 7 | 1,027344406 | 0,001764074 | 2,753483199 | 0,028693571 | WT -MAGE-D1 KO |
| **NOTCH3** | Q9UM47 | Neurogenic locus notch homolog protein 3 | 1,025828334 | 0,00462062 | 2,335299731 | 0,047393897 | WT -MAGE-D1 KO |
| **DIAPH1** | O60610 | Protein diaphanous homolog 1 | 1,020221846 | 0,001767681 | 2,752596084 | 0,028693571 | WT -MAGE-D1 KO |
| **DISP2** | A7MBM2 | Protein dispatched homolog 2 | 1,014013367 | 0,002374814 | 2,624370365 | 0,034948652 | WT -MAGE-D1 KO |
| **KIF13B** | Q9NQT8 | Kinesin-like protein KIF13B | 1,012862353 | 0,003819548 | 2,417988017 | 0,042795316 | WT -MAGE-D1 KO |
| **NCAPG** | Q9BPX3 | Condensin complex subunit 3 | 1,010767726 | 0,00217609 | 2,66232314 | 0,032990346 | WT -MAGE-D1 KO |
| **BRINP1** | O60477 | BMP/retinoic acid-inducible neural-specific protein 1 | 1,009183155 | 0,003319415 | 2,4789385 | 0,039837414 | WT -MAGE-D1 KO |
| **BCHE** | P06276 | Cholinesterase | 5,685178872 | 0,018800682 | 1,725826393 | 0,097775716 | WT -MAGE-D1 KO |
| **TENM1** | Q9UKZ4 | Teneurin-1 | 4,164291957 | 0,052799084 | 1,277373611 | 0,167779946 | WT -MAGE-D1 KO |
| **PLD5** | Q8N7P1 | Inactive phospholipase D5 | 3,77606249 | 0,069922064 | 1,155385761 | 0,195928085 | WT -MAGE-D1 KO |
| **HBB** | P68871 | Hemoglobin subunit beta | 2,150095778 | 0,018145789 | 1,741224153 | 0,095540673 | WT -MAGE-D1 KO |
| **SEMA4F** | O95754 | Semaphorin-4F | 1,723338853 | 0,01054052 | 1,977137959 | 0,071320488 | WT -MAGE-D1 KO |
| **IGFBPL1** | Q8WX77 | Insulin-like growth factor-binding protein-like 1 | 1,571396722 | 0,008813772 | 2,054838187 | 0,064143899 | WT -MAGE-D1 KO |
| **COA5** | Q86WW8 | Cytochrome c oxidase assembly factor 5 | 1,474758614 | 0,011418013 | 1,942409455 | 0,074061626 | WT -MAGE-D1 KO |
| **DYNC1H1** | Q14204 | Cytoplasmic dynein 1 heavy chain 1 | 1,449557366 | 0,008181277 | 2,08717892 | 0,061869702 | WT -MAGE-D1 KO |
| **ULBP2** | Q9BZM5 | UL16-binding protein 2 | 1,415536764 | 0,00901576 | 2,044997653 | 0,064571789 | WT -MAGE-D1 KO |
| **GGT7** | Q9UJ14 | Glutathione hydrolase 7 | 1,342963978 | 0,007039788 | 2,152440391 | 0,057719082 | WT -MAGE-D1 KO |
| **PRKDC** | P78527 | DNA-dependent protein kinase catalytic subunit | 1,310678473 | 0,009001043 | 2,045707182 | 0,064571789 | WT -MAGE-D1 KO |
| **STRA6** | Q9BX79 | Receptor for retinol uptake STRA6 | 1,271177949 | 0,005859635 | 2,23212945 | 0,052613817 | WT -MAGE-D1 KO |
| **SVOP** | Q8N4V2 | Synaptic vesicle 2-related protein | 1,239735362 | 0,005073442 | 2,294697328 | 0,05006797 | WT -MAGE-D1 KO |
| **PIN4** | Q9Y237 | Peptidyl-prolyl cis-trans isomerase NIMA-interacting | 1,215133033 | 0,005300381 | 2,275692876 | 0,051157435 | WT -MAGE-D1 KO |
| **TMEFF1** | Q8IYR6 | Tomoregulin-1 | 1,211534535 | 0,023626389 | 1,62660265 | 0,108634069 | WT -MAGE-D1 KO |
| **ERVMER34-1** | Q9H9K5 | Endogenous retroviral envelope protein HEMO | 1,148095049 | 0,034720971 | 1,459408137 | 0,131751123 | WT -MAGE-D1 KO |
| **APOB** | P04114 | Apolipoprotein B-100 | 1,146829809 | 0,006364687 | 2,196222921 | 0,054549614 | WT -MAGE-D1 KO |
| **C1QL1** | O75973 | C1q-related factor | 1,14250827 | 0,044277451 | 1,35381739 | 0,149955455 | WT -MAGE-D1 KO |
| **ICAM2** | P13598 | Intercellular adhesion molecule 2 | 1,122380418 | 0,005668667 | 2,246519067 | 0,052609089 | WT -MAGE-D1 KO |
| **GGT5** | P36269 | Glutathione hydrolase 5 proenzyme | 1,118118416 | 0,0061185 | 2,213355035 | 0,053582722 | WT -MAGE-D1 KO |
| **PRRT1** | Q99946 | Proline-rich transmembrane protein 1 | 1,107097951 | 0,048900714 | 1,310684799 | 0,159823118 | WT -MAGE-D1 KO |
| **TMEM63B** | Q5T3F8 | CSC1-like protein 2 | 1,104774899 | 0,014704288 | 1,832556011 | 0,085424606 | WT -MAGE-D1 KO |
| **STX1A** | Q16623 | Syntaxin-1A | 1,104392979 | 0,030018239 | 1,522614784 | 0,122124837 | WT -MAGE-D1 KO |
| **MAGI3** | Q5TCQ9 | Membrane-associated guanylate kinase, WW and P | 1,100231777 | 0,033233746 | 1,4784207 | 0,128845911 | WT -MAGE-D1 KO |
| **HECTD4** | Q9Y4D8 | Probable E3 ubiquitin-protein ligase HECTD4 | 1,087586568 | 0,023381829 | 1,631121526 | 0,107972985 | WT -MAGE-D1 KO |
| **ARHGEF2** | Q92974 | Rho guanine nucleotide exchange factor 2 | 1,08140416 | 0,005725301 | 2,2422017 | 0,052609089 | WT -MAGE-D1 KO |
| **RADIL** | Q96JH8 | Ras-associating and dilute domain-containing prote | 1,079736279 | 0,016180861 | 1,790998362 | 0,089356166 | WT -MAGE-D1 KO |
| **RGMB** | Q6NW40 | Repulsive guidance molecule B | 1,063891684 | 0,014353234 | 1,843050233 | 0,084181194 | WT -MAGE-D1 KO |
| **GPT2** | Q8TD30 | Alanine aminotransferase 2 | 1,051828389 | 0,007269589 | 2,138490156 | 0,058034665 | WT -MAGE-D1 KO |
| **SMC4** | Q9NTJ3 | Structural maintenance of chromosomes protein 4 | 1,046588153 | 0,007803265 | 2,107723632 | 0,060352899 | WT -MAGE-D1 KO |
| **POLDIP3** | Q9BY77 | Polymerase delta-interacting protein 3 | 1,044519146 | 0,007578815 | 2,120398691 | 0,059410516 | WT -MAGE-D1 KO |
| **APLP1** | P51693 | Amyloid beta precursor like protein 1 | 1,032846266 | 0,019206003 | 1,716563018 | 0,098792654 | WT -MAGE-D1 KO |
| **NETO2** | Q8NC67 | Neuropilin and tolloid-like protein 2 | 1,032501955 | 0,013393979 | 1,873090392 | 0,080464015 | WT -MAGE-D1 KO |
| **SLC12A6** | Q9UHW9 | Solute carrier family 12 member 6 | 1,01747463 | 0,040333882 | 1,394329981 | 0,143399442 | WT -MAGE-D1 KO |
| **FBLN1** | P23142 | Fibulin-1 | 1,001066123 | 0,018162835 | 1,740816366 | 0,095540673 | WT -MAGE-D1 KO |
| **KIRREL1** | Q96J84 | Kin of IRRE-like protein 1 | -6,594751879 | 0,001808158 | 2,742763519 | 0,029053766 | WT -MAGE-D1 KO |
| **MMP14** | P50281 | Matrix metalloproteinase-14 | -6,04900697 | 1,83991E-05 | 4,735202383 | 0,003540596 | WT -MAGE-D1 KO |
| **NPR 1,00** | P16066 | Atrial natriuretic peptide receptor 1 | -5,864001 | 0,003137234 | 2,5034531 | 0,038632451 | WT -MAGE-D1 KO |
| **PROM1** | O43490 | Prominin-1 | -5,827703091 | 0,003211215 | 2,493330575 | 0,03924276 | WT -MAGE-D1 KO |
| **NTNG1** | Q9Y2I2 | Netrin-G1 | -5,796587365 | 0,003382301 | 2,470787699 | 0,040261913 | WT -MAGE-D1 KO |
| **SYNM** | O15061 | Synemin | -5,616612476 | 0,003759866 | 2,424827662 | 0,042581113 | WT -MAGE-D1 KO |
| **CA12** | O43570 | Carbonic anhydrase 12 | -5,22264804 | 5,09991E-06 | 5,292437436 | 0,002717318 | WT -MAGE-D1 KO |
| **EGFR** | P00533 | Epidermal growth factor receptor | -4,88350666 | 5,90935E-06 | 5,228460489 | 0,002717318 | WT -MAGE-D1 KO |
| **GFRA2** | O00451 | GDNF family receptor alpha-2 | -4,878335463 | 3,73529E-06 | 5,4276755 | 0,002717318 | WT -MAGE-D1 KO |
| **ENPEP** | Q07075 | Glutamyl aminopeptidase | -4,809482703 | 1,63879E-05 | 4,785476959 | 0,003540596 | WT -MAGE-D1 KO |
| **MXRA8** | Q9BRK3 | Matrix remodeling-associated protein 8 | -4,660371119 | 3,35919E-06 | 5,473766007 | 0,002717318 | WT -MAGE-D1 KO |
| **CD44** | P16070 | CD44 antigen | -4,383455645 | 9,34097E-05 | 4,029607857 | 0,006823157 | WT -MAGE-D1 KO |
| **ITGA4** | P13612 | Integrin alpha-4 | -4,295933039 | 1,88227E-05 | 4,725317198 | 0,003540596 | WT -MAGE-D1 KO |
| **CD274** | Q9NZQ7 | Programmed cell death 1 ligand 1 | -4,285956092 | 0,000232773 | 3,633067907 | 0,009843836 | WT -MAGE-D1 KO |
| **TSPAN8** | P19075 | Tetraspanin-8 | -4,224427467 | 4,143E-05 | 4,38268459 | 0,00429536 | WT -MAGE-D1 KO |
| **ICAM1** | P05362 | Intercellular adhesion molecule 1 | -4,177753956 | 3,46738E-05 | 4,459998129 | 0,00384485 | WT -MAGE-D1 KO |
| **MGP** | P08493 | Matrix Gla protein | -3,974039273 | 1,64828E-06 | 5,782967907 | 0,002717318 | WT -MAGE-D1 KO |
| **LINGO2** | Q7L985 | Leucine-rich repeat and immunoglobulin-like doma | -3,951490724 | 0,000119414 | 3,922946079 | 0,0075254 | WT -MAGE-D1 KO |
| **ADGRL3** | Q9HAR2 | Adhesion G protein-coupled receptor L3 | -3,854602799 | 2,63167E-05 | 4,57976789 | 0,003793784 | WT -MAGE-D1 KO |
| **B2M** | P61769 | Beta-2-microglobulin | -3,765494135 | 0,000362131 | 3,441134681 | 0,011998845 | WT -MAGE-D1 KO |
| **HLA-A** | P04439 | HLA class I histocompatibility antigen, A alpha chair | -3,724191852 | 0,000146952 | 3,832825005 | 0,008286021 | WT -MAGE-D1 KO |
| **NTNG2** | Q96CW9 | Netrin-G2 | -3,709756367 | 5,81427E-05 | 4,235504483 | 0,005662751 | WT -MAGE-D1 KO |
| **FAP** | Q12884 | Prolyl endopeptidase FAP | -3,687136347 | 6,22767E-05 | 4,20567447 | 0,005849938 | WT -MAGE-D1 KO |
| **ENPP6** | Q6UWR7 | Glycerophosphocholine cholinephosphodiesterase | -3,628572078 | 3,58488E-05 | 4,445525609 | 0,00384485 | WT -MAGE-D1 KO |
| **SIRPA** | P78324 | Tyrosine-protein phosphatase non-receptor type su | -3,569415897 | 0,000338317 | 3,470676269 | 0,011508804 | WT -MAGE-D1 KO |
| **LAYN** | Q6UX15 | Layilin | -3,556854793 | 0,000231451 | 3,63554063 | 0,009843836 | WT -MAGE-D1 KO |
| **RDH10** | Q8IZV5 | Retinol dehydrogenase 10 | -3,506610016 | 1,92109E-05 | 4,71645162 | 0,003540596 | WT -MAGE-D1 KO |

| | | | | | | | |
|---|---|---|---|---|---|---|---|
| **FAS** | P25445 | Tumor necrosis factor receptor superfamily member | -3,483417578 | 0,000663067 | 3,178442532 | 0,015785908 | WT -MAGE-D1 KO |
| **FCGRT** | P55899 | IgG receptor FcRn large subunit p51 | -3,469041431 | 0,000218088 | 3,661368306 | 0,009804121 | WT -MAGE-D1 KO |
| **NEGR1** | Q7Z3B1 | Neuronal growth regulator 1 | -3,407794557 | 0,000253779 | 3,59554484 | 0,01004851 | WT -MAGE-D1 KO |
| **LYPD6** | Q86Y78 | Ly6/PLAUR domain-containing protein 6 | -3,394990507 | 0,000222155 | 3,653343402 | 0,009804121 | WT -MAGE-D1 KO |
| **GAS1** | P54826 | Growth arrest-specific protein 1 | -3,368433441 | 9,04196E-05 | 4,043737262 | 0,006823157 | WT -MAGE-D1 KO |
| **CNTN3** | Q9P232 | Contactin-3 | -3,249520324 | 6,67361E-06 | 5,175638968 | 0,002717318 | WT -MAGE-D1 KO |
| **PLAUR** | Q03405 | Urokinase plasminogen activator surface receptor | -3,212467751 | 0,003007476 | 2,521797876 | 0,037757917 | WT -MAGE-D1 KO |
| **NPR 2,00** | P20594 | Atrial natriuretic peptide receptor 2 | -3,068312796 | 0,000250031 | 3,602005542 | 0,01004851 | WT -MAGE-D1 KO |
| **SUSD5** | O60279 | Sushi domain-containing protein 5 | -2,937854207 | 7,78652E-05 | 4,108656795 | 0,006572508 | WT -MAGE-D1 KO |
| **NOTCH1** | P46531 | Neurogenic locus notch homolog protein 1 | -2,880913823 | 0,000186129 | 3,730185168 | 0,008961411 | WT -MAGE-D1 KO |
| **PTPRG** | P23470 | Receptor-type tyrosine-protein phosphatase gamma | -2,838166873 | 2,95098E-05 | 4,530033265 | 0,003793784 | WT -MAGE-D1 KO |
| **VSIR** | Q9H7M9 | V-type immunoglobulin domain-containing suppres | -2,800874254 | 0,000311978 | 3,505875538 | 0,011086327 | WT -MAGE-D1 KO |
| **BMPR1B** | O00238 | Bone morphogenetic protein receptor type-1B | -2,760797367 | 0,000416908 | 3,379959375 | 0,012696978 | WT -MAGE-D1 KO |
| **MCAM** | P43121 | Cell surface glycoprotein MUC18 | -2,680582079 | 0,000159228 | 3,797980747 | 0,00852931 | WT -MAGE-D1 KO |
| **IL13RA1** | P78552 | Interleukin-13 receptor subunit alpha-1 | -2,610831193 | 0,000256371 | 3,591130364 | 0,01004851 | WT -MAGE-D1 KO |
| **TSPAN4** | O14817 | Tetraspanin-4 | -2,608481874 | 0,000170207 | 3,76902144 | 0,008547605 | WT -MAGE-D1 KO |
| **RTN3** | O95197 | Reticulon-3 | -2,520426027 | 0,000155867 | 3,807246689 | 0,008490772 | WT -MAGE-D1 KO |
| **ITGA5** | P08648 | Integrin alpha-5 | -2,487844005 | 0,002099363 | 2,67791245 | 0,032235172 | WT -MAGE-D1 KO |
| **ITGB3** | P05106 | Integrin beta-3 | -2,484114002 | 0,000199539 | 3,69997128 | 0,009294489 | WT -MAGE-D1 KO |
| **ANTXR2** | P58335 | Anthrax toxin receptor 2 | -2,479188657 | 0,000611588 | 3,2135412 | 0,014891236 | WT -MAGE-D1 KO |
| **NRP1** | O14786 | Neuropilin-1 | -2,475280585 | 0,000129029 | 3,88931108 | 0,00784112 | WT -MAGE-D1 KO |
| **NOS1** | P29475 | Nitric oxide synthase, brain | -2,461990856 | 2,90624E-05 | 4,536669247 | 0,003793784 | WT -MAGE-D1 KO |
| **BST2** | Q10589 | Bone marrow stromal antigen 2 | -2,438608482 | 0,000265869 | 3,575332762 | 0,010295205 | WT -MAGE-D1 KO |
| **SLC38A1** | Q9H2H9 | Sodium-coupled neutral amino acid symporter 1 | -2,426589877 | 0,00071998 | 3,142679536 | 0,016446668 | WT -MAGE-D1 KO |
| **SNAP23** | O00161 | Synaptosomal-associated protein 23 | -2,391321965 | 9,29426E-05 | 4,031785218 | 0,006823157 | WT -MAGE-D1 KO |
| **ADGRA2** | Q96PE1 | Adhesion G protein-coupled receptor A2 | -2,390758197 | 0,00011719 | 3,931109928 | 0,0075254 | WT -MAGE-D1 KO |
| **TLR2** | O60603 | Toll-like receptor 2 | -2,336209577 | 0,00030333 | 3,5180848 | 0,011078435 | WT -MAGE-D1 KO |
| **FTH1** | P02794 | Ferritin heavy chain | -2,331048161 | 0,001383166 | 2,85912556 | 0,024835178 | WT -MAGE-D1 KO |
| **CD9** | P21926 | CD9 antigen | -2,305350778 | 0,000282984 | 3,54823843 | 0,010594735 | WT -MAGE-D1 KO |
| **CMTM4** | Q8IZR5 | CKLF-like MARVEL transmembrane domain-contai | -2,300594238 | 0,000548875 | 3,260526254 | 0,014459717 | WT -MAGE-D1 KO |
| **IL13RA2** | Q14627 | Interleukin-13 receptor subunit alpha-2 | -2,291667345 | 0,000543762 | 3,264591219 | 0,014443395 | WT -MAGE-D1 KO |
| **SYPL1** | Q16563 | Synaptophysin-like protein 1 | -2,253474819 | 0,000838491 | 3,076501564 | 0,017613793 | WT -MAGE-D1 KO |
| **DLK2** | Q6UY11 | Protein delta homolog 2 | -2,175283539 | 0,00238138 | 2,623171328 | 0,034948652 | WT -MAGE-D1 KO |
| **ERBB4** | Q15303 | Receptor tyrosine-protein kinase erbB-4 | -2,145912312 | 0,000533385 | 3,272959488 | 0,014285819 | WT -MAGE-D1 KO |
| **NT5E** | P21589 | 5'-nucleotidase | -2,114859611 | 0,000661444 | 3,179507237 | 0,015785908 | WT -MAGE-D1 KO |
| **GFRA1** | P56159 | GDNF family receptor alpha-1 | -2,10684014 | 0,000775367 | 3,110492689 | 0,016849564 | WT -MAGE-D1 KO |
| **KIT** | P10721 | Mast/stem cell growth factor receptor Kit | -2,072866433 | 0,001041879 | 2,982182685 | 0,020543554 | WT -MAGE-D1 KO |
| **PPL** | O60437 | Periplakin | -2,068665345 | 0,001031268 | 2,986628257 | 0,020459857 | WT -MAGE-D1 KO |
| **DNASE1L1** | P49184 | Deoxyribonuclease-1-like 1 | -2,037079652 | 0,00130335 | 2,884939091 | 0,02435445 | WT -MAGE-D1 KO |
| **SLC8A1** | P32418 | Sodium/calcium exchanger 1 | -1,987512057 | 0,000688557 | 3,162060066 | 0,016036394 | WT -MAGE-D1 KO |
| **COQ9** | O75208 | Ubiquinone biosynthesis protein COQ9, mitochondr | -1,986449429 | 0,000167663 | 3,775563368 | 0,008547605 | WT -MAGE-D1 KO |
| **AKAP12** | Q02952 | A-kinase anchor protein 12 | -1,975691836 | 7,47376E-05 | 4,126460598 | 0,006572508 | WT -MAGE-D1 KO |
| **BST1** | Q10588 | ADP-ribosyl cyclase/cyclic ADP-ribose hydrolase 2 | -1,973685046 | 0,000168616 | 3,773102127 | 0,008547605 | WT -MAGE-D1 KO |
| **PRDX1** | Q06830 | Peroxiredoxin-1 | -1,940241083 | 0,000151193 | 3,820467078 | 0,008378201 | WT -MAGE-D1 KO |
| **PLS3** | P13797 | Plastin-3 | -1,926406208 | 0,000726642 | 3,138679598 | 0,016446668 | WT -MAGE-D1 KO |
| **NTM** | Q9P121 | Neurotrimin | -1,900437499 | 0,004276189 | 2,368943139 | 0,045649521 | WT -MAGE-D1 KO |
| **ITGA3** | P26006 | Integrin alpha-3 | -1,878155067 | 0,000402958 | 3,394740648 | 0,012696978 | WT -MAGE-D1 KO |
| **PSAP** | P07602 | Prosaposin | -1,867824337 | 0,0001896 | 3,722160584 | 0,008961411 | WT -MAGE-D1 KO |
| **PYCR3** | Q53H96 | Pyrroline-5-carboxylate reductase 3 | -1,842267389 | 0,000248768 | 3,604205563 | 0,01004851 | WT -MAGE-D1 KO |
| **FOLR1** | P15328 | Folate receptor alpha | -1,840729168 | 0,002724757 | 2,564672161 | 0,036891657 | WT -MAGE-D1 KO |
| **GPC4** | O75487 | Glypican-4 | -1,823424531 | 0,000142415 | 3,846442801 | 0,00825647 | WT -MAGE-D1 KO |
| **CD46** | P15529 | Membrane cofactor protein | -1,813281307 | 0,002734793 | 2,563075537 | 0,036891657 | WT -MAGE-D1 KO |
| **ART4** | Q93070 | Ecto-ADP-ribosyltransferase 4 | -1,799756606 | 0,000768748 | 3,11421626 | 0,016849564 | WT -MAGE-D1 KO |
| **NRP2** | O60462 | Neuropilin-2 | -1,77646862 | 0,00023109 | 3,636219356 | 0,009843836 | WT -MAGE-D1 KO |
| **EPHA2** | P29317 | Ephrin type-A receptor 2 | -1,768903183 | 0,002223856 | 2,652893359 | 0,033521085 | WT -MAGE-D1 KO |
| **MELTF** | P08582 | Melanotransferrin | -1,74442963 | 0,000388488 | 3,410622344 | 0,012486006 | WT -MAGE-D1 KO |
| **LORICRIN** | P23490 | Loricrin | -1,744327385 | 0,002869606 | 2,542177705 | 0,036891657 | WT -MAGE-D1 KO |
| **PRDX2** | P32119 | Peroxiredoxin-2 | -1,728374994 | 0,001664616 | 2,77868597 | 0,028214835 | WT -MAGE-D1 KO |
| **SCARB1** | Q8WTV0 | Scavenger receptor class B member 1 | -1,727276109 | 0,004728909 | 2,325239068 | 0,047981152 | WT -MAGE-D1 KO |
| **GPNMB** | Q14956 | Transmembrane glycoprotein NMB | -1,726357316 | 0,000164059 | 3,784999807 | 0,008547605 | WT -MAGE-D1 KO |
| **A2M** | P01023 | Alpha-2-macroglobulin | -1,713188985 | 0,000610641 | 3,214213776 | 0,014891236 | WT -MAGE-D1 KO |
| **COQ7** | Q99807 | 5-demethoxyubiquinone hydroxylase, mitochondria | -1,705722397 | 0,000794334 | 3,099996663 | 0,016907222 | WT -MAGE-D1 KO |
| **RPS6KA3** | P51812 | Ribosomal protein S6 kinase alpha-3 | -1,679966324 | 0,000269814 | 3,568935424 | 0,0103236 | WT -MAGE-D1 KO |
| **LY6H** | O94772 | Lymphocyte antigen 6H | -1,679071207 | 0,002106218 | 2,676496592 | 0,032235172 | WT -MAGE-D1 KO |
| **TAGLN** | Q01995 | Transgelin | -1,678154718 | 0,000254646 | 3,594062821 | 0,01004851 | WT -MAGE-D1 KO |
| **PLAU** | P00749 | Urokinase-type plasminogen activator | -1,669935167 | 0,004871926 | 2,312299278 | 0,048628483 | WT -MAGE-D1 KO |
| **CD151** | P48509 | CD151 antigen | -1,65737438 | 0,00181612 | 2,740855541 | 0,029053766 | WT -MAGE-D1 KO |
| **TIMP1** | P01033 | Metalloproteinase inhibitor 1 | -1,651917141 | 0,000578017 | 3,238059345 | 0,014853467 | WT -MAGE-D1 KO |
| **FLNC** | Q14315 | Filamin-C | -1,64947416 | 0,000179391 | 3,7462004 | 0,008870174 | WT -MAGE-D1 KO |
| **FLRT3** | Q9NZU0 | Leucine-rich repeat transmembrane protein FLRT3 | -1,628790987 | 0,00138263 | 2,859293975 | 0,024835178 | WT -MAGE-D1 KO |
| **GPC6** | Q9Y625 | Glypican-6 | -1,628336272 | 0,00043095 | 3,365572847 | 0,012707102 | WT -MAGE-D1 KO |
| **EPDR1** | Q9UM22 | Mammalian ependymin-related protein 1 | -1,61897839 | 0,00034376 | 3,463744449 | 0,011508804 | WT -MAGE-D1 KO |
| **ATP1B3** | P54709 | Sodium/potassium-transporting ATPase subunit bet | -1,616405879 | 0,003900716 | 2,408855674 | 0,043082134 | WT -MAGE-D1 KO |
| **LRP1B** | Q9NZR2 | Low-density lipoprotein receptor-related protein 1B | -1,60352182 | 0,003861432 | 2,413251638 | 0,042795316 | WT -MAGE-D1 KO |
| **VCAM1** | P19320 | Vascular cell adhesion protein 1 | -1,584496592 | 0,000596172 | 3,224628318 | 0,014853467 | WT -MAGE-D1 KO |
| **ATP2B3** | Q16720 | Plasma membrane calcium-transporting ATPase 3 | -1,583179298 | 0,000594618 | 3,225762301 | 0,014853467 | WT -MAGE-D1 KO |
| **TM9SF2** | Q99805 | Transmembrane 9 superfamily member 2 | -1,573406716 | 0,000317344 | 3,498470356 | 0,011086327 | WT -MAGE-D1 KO |
| **FLNB** | O75369 | Filamin-B | -1,573201136 | 0,000422205 | 3,374476117 | 0,012696978 | WT -MAGE-D1 KO |
| **PLXNC1** | O60486 | Plexin-C1 | -1,539795688 | 0,001128307 | 2,947572719 | 0,021714842 | WT -MAGE-D1 KO |
| **SLC7A5** | Q01650 | Large neutral amino acids transporter small subunit | -1,534668062 | 0,004035726 | 2,394078373 | 0,043820345 | WT -MAGE-D1 KO |
| **AHNAK** | Q09666 | Neuroblast differentiation-associated protein AHNA | -1,523698683 | 0,000455614 | 3,341403372 | 0,01319227 | WT -MAGE-D1 KO |
| **BAK1** | Q16611 | Bcl-2 homologous antagonist/killer | -1,507521685 | 0,000936745 | 3,028378501 | 0,018937518 | WT -MAGE-D1 KO |
| **SCARF2** | Q96GP6 | Scavenger receptor class F member 2 | -1,49049764 | 0,000378525 | 3,421905806 | 0,01228867 | WT -MAGE-D1 KO |
| **ATP11A** | P98196 | Phospholipid-transporting ATPase IH | -1,470714702 | 0,002342384 | 2,63034198 | 0,034853801 | WT -MAGE-D1 KO |
| **THY1** | P04216 | Thy-1 membrane glycoprotein | -1,466025721 | 0,000560587 | 3,251357312 | 0,014530042 | WT -MAGE-D1 KO |
| **HEATR5A** | Q86XA9 | HEAT repeat-containing protein 5A | -1,461839535 | 0,000506058 | 3,295799649 | 0,014021298 | WT -MAGE-D1 KO |

| | | | | | | | |
|---|---|---|---|---|---|---|---|
| **BCL2** | P10415 | Apoptosis regulator Bcl-2 | -1,442257192 | 0,000409808 | 3,38741998 | 0,012696978 | WT -MAGE-D1 KO |
| **CDK6** | Q00534 | Cyclin-dependent kinase 6 | -1,438855848 | 0,000340403 | 3,468006307 | 0,011508804 | WT -MAGE-D1 KO |
| **MAOA** | P21397 | Amine oxidase [flavin-containing] A | -1,428168734 | 0,000855827 | 3,067613947 | 0,017861224 | WT -MAGE-D1 KO |
| **TGM2** | P21980 | Protein-glutamine gamma-glutamyltransferase 2 | -1,41548417 | 0,000427773 | 3,368786957 | 0,012707102 | WT -MAGE-D1 KO |
| **MYADM** | Q96S97 | Myeloid-associated differentiation marker | -1,403702857 | 0,003952334 | 2,403146357 | 0,043502745 | WT -MAGE-D1 KO |
| **DOCK4** | Q8N1I0 | Dedicator of cytokinesis protein 4 | -1,38593372 | 0,001174201 | 2,930257392 | 0,022330671 | WT -MAGE-D1 KO |
| **TM7SF2** | O76062 | Delta(14)-sterol reductase TM7SF2 | -1,376300958 | 0,0005314 | 3,274578352 | 0,014285819 | WT -MAGE-D1 KO |
| **CNTNAP1** | P78357 | Contactin-associated protein 1 | -1,371992871 | 0,000525263 | 3,279623557 | 0,014285819 | WT -MAGE-D1 KO |
| **COMT** | P21964 | Catechol O-methyltransferase | -1,342232899 | 0,003246256 | 2,488617187 | 0,039520712 | WT -MAGE-D1 KO |
| **CNPY2** | Q9Y2B0 | Protein canopy homolog 2 | -1,334373174 | 0,001380179 | 2,860064543 | 0,024835178 | WT -MAGE-D1 KO |
| **MAOB** | P27338 | Amine oxidase [flavin-containing] B | -1,313434871 | 0,002441369 | 2,612366492 | 0,035371868 | WT -MAGE-D1 KO |
| **ATPAF1** | Q5TC12 | ATP synthase mitochondrial F1 complex assembly f | -1,302365612 | 0,002821505 | 2,549519208 | 0,036891657 | WT -MAGE-D1 KO |
| **NOTCH2** | Q04721 | Neurogenic locus notch homolog protein 2 | -1,291679917 | 0,000604255 | 3,218780102 | 0,014891236 | WT -MAGE-D1 KO |
| **SIPA1L2** | Q9P2F8 | Signal-induced proliferation-associated 1-like prote | -1,28258971 | 0,000773937 | 3,111294619 | 0,016849564 | WT -MAGE-D1 KO |
| **NPTN** | Q9Y639 | Neuroplastin | -1,258929439 | 0,003711117 | 2,43049539 | 0,042556412 | WT -MAGE-D1 KO |
| **EFNB1** | P98172 | Ephrin-B1 | -1,258023622 | 0,003849992 | 2,414540158 | 0,042795316 | WT -MAGE-D1 KO |
| **VIM** | P08670 | Vimentin | -1,256958329 | 0,001766056 | 2,75299542 | 0,028693571 | WT -MAGE-D1 KO |
| **ABCA1** | O95477 | Phospholipid-transporting ATPase ABCA1 | -1,254893831 | 0,00151353 | 2,820008871 | 0,026576581 | WT -MAGE-D1 KO |
| **BPNT2** | Q9NX62 | Golgi-resident adenosine 3',5'-bisphosphate 3'-pho | -1,254519205 | 0,000671591 | 3,172895033 | 0,015871279 | WT -MAGE-D1 KO |
| **KRT85** | P78386 | Keratin, type II cuticular Hb5 | -1,230682916 | 0,00278107 | 2,555788134 | 0,036891657 | WT -MAGE-D1 KO |
| **POGLUT1** | Q8NBL1 | Protein O-glucosyltransferase 1 | -1,225957548 | 0,002639492 | 2,578479598 | 0,036724365 | WT -MAGE-D1 KO |
| **MCCC1** | Q96RQ3 | Methylcrotonoyl-CoA carboxylase subunit alpha, mi | -1,213593782 | 0,001138557 | 2,943645272 | 0,021781677 | WT -MAGE-D1 KO |
| **CNTN1** | Q12860 | Contactin-1 | -1,205451904 | 0,001329322 | 2,876369727 | 0,024413953 | WT -MAGE-D1 KO |
| **FRY** | Q5TBA9 | Protein furry homolog | -1,205366625 | 0,001978324 | 2,703702684 | 0,030865688 | WT -MAGE-D1 KO |
| **CACNA2D1** | P54289 | Voltage-dependent calcium channel subunit alpha- | -1,205176049 | 0,00181699 | 2,740647382 | 0,029053766 | WT -MAGE-D1 KO |
| **RAB27B** | O00194 | Ras-related protein Rab-27B | -1,201657714 | 0,002634362 | 2,579324577 | 0,036724365 | WT -MAGE-D1 KO |
| **SLC3A2** | P08195 | 4F2 cell-surface antigen heavy chain | -1,199541756 | 0,001884759 | 2,724744238 | 0,029840466 | WT -MAGE-D1 KO |
| **SEMA6D** | Q8NFY4 | Semaphorin-6D | -1,190171577 | 0,004505748 | 2,346233071 | 0,047170929 | WT -MAGE-D1 KO |
| **EPHX2** | P34913 | Bifunctional epoxide hydrolase 2 | -1,189093167 | 0,004644056 | 2,333102559 | 0,047393897 | WT -MAGE-D1 KO |
| **DCBLD2** | Q96PD2 | Discoidin, CUB and LCCL domain-containing prote | -1,184398878 | 0,000936859 | 3,028325692 | 0,018937518 | WT -MAGE-D1 KO |
| **TTC27** | Q6P3X3 | Tetratricopeptide repeat protein 27 | -1,178567805 | 0,002744499 | 2,561536992 | 0,036891657 | WT -MAGE-D1 KO |
| **S100A7** | P31151 | Protein S100-A7 | -1,176618644 | 0,002557077 | 2,592256205 | 0,036364802 | WT -MAGE-D1 KO |
| **PTPRR** | Q15256 | Receptor-type tyrosine-protein phosphatase R | -1,151425424 | 0,001742711 | 2,758774547 | 0,028693571 | WT -MAGE-D1 KO |
| **COTL1** | Q14019 | Coactosin-like protein | -1,145853203 | 0,00332185 | 2,478619938 | 0,039837414 | WT -MAGE-D1 KO |
| **COL3A1** | P02461 | Collagen alpha-1(III) chain | -1,135966914 | 0,003538352 | 2,451198913 | 0,041495519 | WT -MAGE-D1 KO |
| **LAMP2** | P13473 | Lysosome-associated membrane glycoprotein 2 | -1,125086924 | 0,002097249 | 2,678350008 | 0,032235172 | WT -MAGE-D1 KO |
| **ERLIN1** | O75477 | Erlin-1 | -1,123306167 | 0,003102748 | 2,50825346 | 0,038417226 | WT -MAGE-D1 KO |
| **GLIPR2** | Q9H4G4 | Golgi-associated plant pathogenesis-related protein | -1,112701863 | 0,002500853 | 2,601911869 | 0,035723293 | WT -MAGE-D1 KO |
| **VDAC3** | Q9Y277 | Voltage-dependent anion-selective channel protein | -1,099152417 | 0,002443647 | 2,611961567 | 0,035371868 | WT -MAGE-D1 KO |
| **RYK** | P34925 | Tyrosine-protein kinase RYK | -1,093065568 | 0,002670656 | 2,573382004 | 0,03683901 | WT -MAGE-D1 KO |
| **MPDU1** | O75352 | Mannose-P-dolichol utilization defect 1 protein | -1,08944704 | 0,00316225 | 2,500003837 | 0,038791873 | WT -MAGE-D1 KO |
| **VASN** | Q6EMK4 | Vasorin | -1,088375961 | 0,003834223 | 2,416322646 | 0,042795316 | WT -MAGE-D1 KO |
| **GAA** | P10253 | Lysosomal alpha-glucosidase | -1,087749399 | 0,004017286 | 2,396067267 | 0,043767989 | WT -MAGE-D1 KO |
| **DHRS3** | O75911 | Short-chain dehydrogenase/reductase 3 | -1,087652152 | 0,001849265 | 2,73300094 | 0,029423448 | WT -MAGE-D1 KO |
| **NDST1** | P52848 | Bifunctional heparan sulfate N-deacetylase/N-sulfot | -1,071694192 | 0,001667958 | 2,777814763 | 0,028214835 | WT -MAGE-D1 KO |
| **VDAC2** | P45880 | Voltage-dependent anion-selective channel protein | -1,053341694 | 0,001765201 | 2,753205895 | 0,028693571 | WT -MAGE-D1 KO |
| **PCMTD2** | Q9NV79 | Protein-L-isoaspartate O-methyltransferase domain | -1,036023661 | 0,004641119 | 2,33337725 | 0,047393897 | WT -MAGE-D1 KO |
| **ANXA4** | P09525 | Annexin A4 | -1,024598328 | 0,003720707 | 2,429374548 | 0,042556412 | WT -MAGE-D1 KO |
| **NLGN1** | Q8N2Q7 | Neuroligin-1 | -1,016127695 | 0,003308352 | 2,480388327 | 0,039837414 | WT -MAGE-D1 KO |
| **ALOX12B** | O75342 | Arachidonate 12-lipoxygenase, 12R-type | -1,011963626 | 0,002630972 | 2,579883752 | 0,036724365 | WT -MAGE-D1 KO |
| **OSMR** | Q99650 | Oncostatin-M-specific receptor subunit beta | -5,181939919 | 0,005458012 | 2,26296552 | 0,052053562 | WT -MAGE-D1 KO |
| **HGF** | P14210 | Hepatocyte growth factor | -4,87051459 | 0,00718772 | 2,14340883 | 0,057849355 | WT -MAGE-D1 KO |
| **TMEM192** | Q8IY95 | Transmembrane protein 192 | -4,485440628 | 0,01009861 | 1,995738391 | 0,069203978 | WT -MAGE-D1 KO |
| **MYRF** | Q9Y2G1 | Myelin regulatory factor | -4,413571871 | 0,011040089 | 1,957027408 | 0,072860056 | WT -MAGE-D1 KO |
| **CSF1** | P09603 | Macrophage colony-stimulating factor 1 | -4,367628399 | 0,011436487 | 1,941707364 | 0,074061626 | WT -MAGE-D1 KO |
| **SDK2** | Q58EX2 | Protein sidekick-2 | -4,241387119 | 0,013312124 | 1,87575264 | 0,080122036 | WT -MAGE-D1 KO |
| **EPPK1** | P58107 | Epiplakin | -4,166455904 | 0,013522789 | 1,868933714 | 0,08093528 | WT -MAGE-D1 KO |
| **CA10** | Q9NS85 | Carbonic anhydrase-related protein 10 | -4,122455305 | 0,01502677 | 1,823134351 | 0,085702032 | WT -MAGE-D1 KO |
| **COLEC12** | Q5KU26 | Collectin-12 | -4,096218363 | 0,014781063 | 1,83029434 | 0,085424606 | WT -MAGE-D1 KO |
| **IL4R** | P24394 | Interleukin-4 receptor subunit alpha | -4,068319423 | 0,015052188 | 1,822400365 | 0,085702032 | WT -MAGE-D1 KO |
| **CA4** | P22748 | Carbonic anhydrase 4 | -3,957772085 | 0,017446467 | 1,758292508 | 0,092989958 | WT -MAGE-D1 KO |
| **HLA-E** | P13747 | HLA class I histocompatibility antigen, alpha chain E | -3,926026598 | 0,017089487 | 1,767270974 | 0,092378338 | WT -MAGE-D1 KO |
| **PDE5A** | O76074 | cGMP-specific 3',5'-cyclic phosphodiesterase | -3,907332345 | 0,018313917 | 1,73721875 | 0,095933923 | WT -MAGE-D1 KO |
| **LAMA5** | O15230 | Laminin subunit alpha-5 | -3,793517287 | 0,021948212 | 1,658600856 | 0,10435141 | WT -MAGE-D1 KO |
| **TNFRSF1A** | P19438 | Tumor necrosis factor receptor superfamily member | -3,775262154 | 0,020175943 | 1,69516616 | 0,100070186 | WT -MAGE-D1 KO |
| **LGI1** | O95970 | Leucine-rich glioma-inactivated protein 1 | -3,769463608 | 0,020416512 | 1,69001846 | 0,100397488 | WT -MAGE-D1 KO |
| **FGFR2** | P21802 | Fibroblast growth factor receptor 2 | -3,492353084 | 0,026737094 | 1,572885792 | 0,115816617 | WT -MAGE-D1 KO |
| **HLA-C** | P10321 | HLA class I histocompatibility antigen, C alpha chai | -3,442201088 | 0,028597144 | 1,543677334 | 0,120262655 | WT -MAGE-D1 KO |
| **HBEGF** | Q99075 | Proheparin-binding EGF-like growth factor | -3,389622766 | 0,030802174 | 1,511418625 | 0,123652047 | WT -MAGE-D1 KO |
| **EIF2B5** | Q13144 | Translation initiation factor eIF-2B subunit epsilon | -3,202645416 | 0,037595749 | 1,424861257 | 0,13809622 | WT -MAGE-D1 KO |
| **VIPR2** | P41587 | Vasoactive intestinal polypeptide receptor 2 | -3,195608859 | 0,037425499 | 1,426832398 | 0,1379958 | WT -MAGE-D1 KO |
| **SGCD** | Q92629 | Delta-sarcoglycan | -3,171893713 | 0,038385053 | 1,415837852 | 0,140033554 | WT -MAGE-D1 KO |
| **NTRK3** | Q16288 | NT-3 growth factor receptor | -3,159578019 | 0,039571116 | 1,4026217 | 0,14210231 | WT -MAGE-D1 KO |
| **ENTPD1** | P49961 | Ectonucleoside triphosphate diphosphohydrolase 1 | -3,027842498 | 0,044125941 | 1,355306017 | 0,149955455 | WT -MAGE-D1 KO |
| **MTCH1** | Q9NZJ7 | Mitochondrial carrier homolog 1 | -2,993547535 | 0,045949221 | 1,337721851 | 0,153994573 | WT -MAGE-D1 KO |
| **GLIPR1** | P48060 | Glioma pathogenesis-related protein 1 | -2,977044558 | 0,047106916 | 1,326915327 | 0,15624523 | WT -MAGE-D1 KO |
| **CRIM1** | Q9NZV1 | Cysteine-rich motor neuron 1 protein | -2,948084577 | 0,048666468 | 1,312770169 | 0,159660954 | WT -MAGE-D1 KO |
| **CORIN** | Q9Y5Q5 | Atrial natriuretic peptide-converting enzyme | -2,92246169 | 0,051156607 | 1,291098266 | 0,164398106 | WT -MAGE-D1 KO |
| **MTMR14** | Q8NCE2 | Myotubularin-related protein 14 | -2,89424133 | 0,061894471 | 1,208348142 | 0,181670165 | WT -MAGE-D1 KO |
| **KRT31** | Q15323 | Keratin, type I cuticular Ha1 | -2,837327089 | 0,060126174 | 1,220936433 | 0,178990063 | WT -MAGE-D1 KO |
| **TMEM132C** | Q8N3T6 | Transmembrane protein 132C | -2,830896781 | 0,055043981 | 1,259290163 | 0,171544263 | WT -MAGE-D1 KO |
| **ALOXE3** | Q9BYJ1 | Hydroperoxide isomerase ALOXE3 | -2,808548544 | 0,058221403 | 1,234917336 | 0,176365305 | WT -MAGE-D1 KO |
| **FZD6** | O60353 | Frizzled-6 | -2,80123136 | 0,056760747 | 1,245951895 | 0,173707322 | WT -MAGE-D1 KO |
| **WNT5A** | P41221 | Protein Wnt-5a | -2,798554708 | 0,058312272 | 1,234240035 | 0,17642167 | WT -MAGE-D1 KO |
| **PLPPR4** | Q7Z2D5 | Phospholipid phosphatase-related protein type 4 | -2,731767132 | 0,06176048 | 1,209289339 | 0,181608584 | WT -MAGE-D1 KO |
| **SUSD2** | Q9UGT4 | Sushi domain-containing protein 2 | -2,71889905 | 0,067701948 | 1,169398837 | 0,192215751 | WT -MAGE-D1 KO |
| **NRG1** | Q02297 | Pro-neuregulin-1, membrane-bound isoform | -2,711387199 | 0,064437403 | 1,190861968 | 0,187762298 | WT -MAGE-D1 KO |
| [illegible] | [illegible] | [illegible] | [illegible] | [illegible] | [illegible] | [illegible] | [illegible] |

| | | | | | | | |
|---|---|---|---|---|---|---|---|
| **SDSL** | Q96GA7 | Serine dehydratase-like | -2,682627806 | 0,065231046 | 1,185545658 | 0,188882625 | WT -MAGE-D1 KO |
| **CAMK2G** | Q13555 | Calcium/calmodulin-dependent protein kinase type | -2,65514085 | 0,067982821 | 1,167600817 | 0,192677943 | WT -MAGE-D1 KO |
| **MAGEA10** | P43363 | Melanoma-associated antigen 10 | -2,621315778 | 0,07018599 | 1,153749573 | 0,196154583 | WT -MAGE-D1 KO |
| **SLC38A3** | Q99624 | Sodium-coupled neutral amino acid transporter 3 | -1,861846785 | 0,031341304 | 1,503882941 | 0,124518184 | WT -MAGE-D1 KO |
| **XP32** | Q5T750 | Skin-specific protein 32 | -1,701391769 | 0,00721767 | 2,14160298 | 0,057849355 | WT -MAGE-D1 KO |
| **SFRP1** | Q8N474 | Secreted frizzled-related protein 1 | -1,615546715 | 0,006004868 | 2,221496568 | 0,05311247 | WT -MAGE-D1 KO |
| **KPRP** | Q5T749 | Keratinocyte proline-rich protein | -1,566431175 | 0,009682351 | 2,014019174 | 0,067650166 | WT -MAGE-D1 KO |
| **TYRO3** | Q06418 | Tyrosine-protein kinase receptor TYRO3 | -1,56404305 | 0,017101777 | 1,766958763 | 0,092378338 | WT -MAGE-D1 KO |
| **CCNY** | Q8ND76 | Cyclin-Y | -1,56149475 | 0,027932854 | 1,553884686 | 0,119066423 | WT -MAGE-D1 KO |
| **FKBP7** | Q9Y680 | Peptidyl-prolyl cis-trans isomerase FKBP7 | -1,558758073 | 0,013796266 | 1,860238434 | 0,081509558 | WT -MAGE-D1 KO |
| **BVES** | Q8NE79 | Blood vessel epicardial substance | -1,495739467 | 0,013749434 | 1,861715193 | 0,081382467 | WT -MAGE-D1 KO |
| **TNFRSF10B** | O14763 | Tumor necrosis factor receptor superfamily member | -1,457967526 | 0,016755742 | 1,775836323 | 0,091225305 | WT -MAGE-D1 KO |
| **PIANP** | Q8IYJ0 | PILR alpha-associated neural protein | -1,4495215 | 0,020312861 | 1,692228896 | 0,100397488 | WT -MAGE-D1 KO |
| **LGALS8** | O00214 | Galectin-8 | -1,416010403 | 0,010141279 | 1,993907252 | 0,069203978 | WT -MAGE-D1 KO |
| **ADAM11** | O75078 | Disintegrin and metalloproteinase domain-containir | -1,41334577 | 0,008852965 | 2,052911272 | 0,064143899 | WT -MAGE-D1 KO |
| **IL10RB** | Q08334 | Interleukin-10 receptor subunit beta | -1,404743332 | 0,008621384 | 2,064423001 | 0,063993369 | WT -MAGE-D1 KO |
| **MVP** | Q14764 | Major vault protein | -1,342049992 | 0,007074906 | 2,15027935 | 0,057849355 | WT -MAGE-D1 KO |
| **ACVR1** | Q04771 | Activin receptor type-1 | -1,32916515 | 0,007901328 | 2,102299935 | 0,060608274 | WT -MAGE-D1 KO |
| **TGFB1** | P01137 | Transforming growth factor beta-1 proprotein | -1,328985769 | 0,009827462 | 2,007558618 | 0,067925728 | WT -MAGE-D1 KO |
| **ACVR1B** | P36896 | Activin receptor type-1B | -1,309076616 | 0,017730356 | 1,751282544 | 0,093725928 | WT -MAGE-D1 KO |
| **PTPRE** | P23469 | Receptor-type tyrosine-protein phosphatase epsilor | -1,307526847 | 0,035672856 | 1,44766212 | 0,133634441 | WT -MAGE-D1 KO |
| **SLC5A6** | Q9Y289 | Sodium-dependent multivitamin transporter | -1,304269096 | 0,023669606 | 1,625808965 | 0,108677307 | WT -MAGE-D1 KO |
| **ETFDH** | Q16134 | Electron transfer flavoprotein-ubiquinone oxidoredu | -1,298290739 | 0,007468438 | 2,126770206 | 0,058832256 | WT -MAGE-D1 KO |
| **ECHDC1** | Q9NTX5 | Ethylmalonyl-CoA decarboxylase | -1,296584475 | 0,012742796 | 1,894735264 | 0,078430183 | WT -MAGE-D1 KO |
| **FRRS1L** | Q9P0K9 | DOMON domain-containing protein FRRS1L | -1,277606177 | 0,013267322 | 1,877216721 | 0,080002202 | WT -MAGE-D1 KO |
| **PCDHB6** | Q9Y5E3 | Protocadherin beta-6 | -1,27599754 | 0,005738241 | 2,241221219 | 0,052609089 | WT -MAGE-D1 KO |
| **NGFR** | P08138 | Tumor necrosis factor receptor superfamily member | -1,271282697 | 0,047077747 | 1,327184329 | 0,15624523 | WT -MAGE-D1 KO |
| **DEGS1** | O15121 | Sphingolipid delta(4)-desaturase DES1 | -1,264097764 | 0,025463815 | 1,594076535 | 0,113039642 | WT -MAGE-D1 KO |
| **ERBB3** | P21860 | Receptor tyrosine-protein kinase erbB-3 | -1,237819283 | 0,025879095 | 1,587050916 | 0,113918887 | WT -MAGE-D1 KO |
| **TXNDC12** | O95881 | Thioredoxin domain-containing protein 12 | -1,221780457 | 0,035285185 | 1,452407604 | 0,133262731 | WT -MAGE-D1 KO |
| **BDH1** | Q02338 | D-beta-hydroxybutyrate dehydrogenase, mitochond | -1,212024248 | 0,012035116 | 1,919549721 | 0,075878639 | WT -MAGE-D1 KO |
| **WBP11** | Q9Y2W2 | WW domain-binding protein 11 | -1,208787483 | 0,031017608 | 1,508391697 | 0,123804046 | WT -MAGE-D1 KO |
| **TPM1** | P09493 | Tropomyosin alpha-1 chain | -1,204512432 | 0,021805916 | 1,661425663 | 0,104258377 | WT -MAGE-D1 KO |
| **SDC2** | P34741 | Syndecan-2 | -1,194917947 | 0,00593234 | 2,226773953 | 0,052815904 | WT -MAGE-D1 KO |
| **KRT82** | Q9NSB4 | Keratin, type II cuticular Hb2 | -1,190341255 | 0,006733998 | 2,171727048 | 0,056070124 | WT -MAGE-D1 KO |
| **GSPT2** | Q8IYD1 | Eukaryotic peptide chain release factor GTP-binding | -1,17941947 | 0,021888028 | 1,659793371 | 0,104258377 | WT -MAGE-D1 KO |
| **DAG1** | Q14118 | Dystroglycan 1 | -1,175199086 | 0,005543656 | 2,25620376 | 0,052355867 | WT -MAGE-D1 KO |
| **NCR3LG1** | Q68D85 | Natural cytotoxicity triggering receptor 3 ligand 1 | -1,166916761 | 0,010291012 | 1,987541902 | 0,069926667 | WT -MAGE-D1 KO |
| **GHDC** | Q8N2G8 | GH3 domain-containing protein | -1,165575557 | 0,04812033 | 1,317671404 | 0,158624349 | WT -MAGE-D1 KO |
| **SLC20A1** | Q8WUM9 | Sodium-dependent phosphate transporter 1 | -1,165562561 | 0,009805674 | 2,008522532 | 0,067921202 | WT -MAGE-D1 KO |
| **MRPS15** | P82914 | 28S ribosomal protein S15, mitochondrial | -1,156559931 | 0,01015941 | 1,993131527 | 0,069203978 | WT -MAGE-D1 KO |
| **ANPEP** | P15144 | Aminopeptidase N | -1,151331895 | 0,009020763 | 2,044756715 | 0,064571789 | WT -MAGE-D1 KO |
| **ARHGEF17** | Q96PE2 | Rho guanine nucleotide exchange factor 17 | -1,144565435 | 0,014804451 | 1,829607694 | 0,085424606 | WT -MAGE-D1 KO |
| **FNDC10** | F2Z333 | Fibronectin type III domain-containing protein 10 | -1,10117272 | 0,051252925 | 1,29028134 | 0,164398106 | WT -MAGE-D1 KO |
| **PKD2** | Q13563 | Polycystin-2 | -1,09601489 | 0,022941383 | 1,639380411 | 0,106676706 | WT -MAGE-D1 KO |
| **GMPPA** | Q96IJ6 | Mannose-1-phosphate guanyltransferase alpha | -1,095370233 | 0,012762597 | 1,894060955 | 0,078430183 | WT -MAGE-D1 KO |
| **DDR1** | Q08345 | Epithelial discoidin domain-containing receptor 1 | -1,090353123 | 0,021525274 | 1,667051305 | 0,10374662 | WT -MAGE-D1 KO |
| **HBA2** | P69905 | Hemoglobin subunit alpha | -1,088476093 | 0,028479948 | 1,545460804 | 0,120262655 | WT -MAGE-D1 KO |
| **SLC30A9** | Q6PML9 | Zinc transporter 9 | -1,076461607 | 0,03667462 | 1,435634381 | 0,136557821 | WT -MAGE-D1 KO |
| **STARD3NL** | O95772 | STARD3 N-terminal-like protein | -1,075769045 | 0,005764154 | 2,239264437 | 0,052609089 | WT -MAGE-D1 KO |
| **CACHD1** | Q5VU97 | VWFA and cache domain-containing protein 1 | -1,075665423 | 0,008835834 | 2,053752449 | 0,064143899 | WT -MAGE-D1 KO |
| **APH1A** | Q96BI3 | Gamma-secretase subunit APH-1A | -1,072407721 | 0,029425716 | 1,531272958 | 0,121219127 | WT -MAGE-D1 KO |
| **HAL** | P42357 | Histidine ammonia-lyase | -1,070573128 | 0,016555323 | 1,781062333 | 0,090774587 | WT -MAGE-D1 KO |
| **RELL1** | Q8IUW5 | RELT-like protein 1 | -1,069523872 | 0,011452591 | 1,941096242 | 0,074061626 | WT -MAGE-D1 KO |
| **KEAP1** | Q14145 | Kelch-like ECH-associated protein 1 | -1,066338178 | 0,016774784 | 1,77534307 | 0,091225305 | WT -MAGE-D1 KO |
| **NELFE** | P18615 | Negative elongation factor E | -1,06372389 | 0,02167115 | 1,664118038 | 0,103956831 | WT -MAGE-D1 KO |
| **DPY19L3** | Q6ZPD9 | Probable C-mannosyltransferase DPY19L3 | -1,063297569 | 0,015393593 | 1,812660008 | 0,086561052 | WT -MAGE-D1 KO |
| **ROBO1** | Q9Y6N7 | Roundabout homolog 1 | -1,063020537 | 0,005478163 | 2,261365024 | 0,052055644 | WT -MAGE-D1 KO |
| **TM9SF3** | Q9HD45 | Transmembrane 9 superfamily member 3 | -1,061560434 | 0,005759396 | 2,239623045 | 0,052609089 | WT -MAGE-D1 KO |
| **LGR4** | Q9BXB1 | Leucine-rich repeat-containing G-protein coupled re | -1,059659652 | 0,019659732 | 1,706422408 | 0,098996209 | WT -MAGE-D1 KO |
| **PTPRA** | P18433 | Receptor-type tyrosine-protein phosphatase alpha | -1,04810893 | 0,028520846 | 1,544837596 | 0,120262655 | WT -MAGE-D1 KO |
| **CALU** | O43852 | Calumenin | -1,037027338 | 0,031921714 | 1,495913797 | 0,12603979 | WT -MAGE-D1 KO |
| **PCDH7** | O60245 | Protocadherin-7 | -1,034419674 | 0,019807305 | 1,703174619 | 0,09918973 | WT -MAGE-D1 KO |
| **SLC4A2** | P04920 | Anion exchange protein 2 | -1,032754923 | 0,00871039 | 2,059962414 | 0,064143899 | WT -MAGE-D1 KO |
| **NCOA2** | Q15596 | Nuclear receptor coactivator 2 | -1,031529136 | 0,029319858 | 1,532838139 | 0,121123423 | WT -MAGE-D1 KO |
| **DCD** | P81605 | Dermcidin | -1,030463678 | 0,0244239 | 1,612184984 | 0,110561148 | WT -MAGE-D1 KO |
| **ATP2B4** | P23634 | Plasma membrane calcium-transporting ATPase 4 | -1,024224958 | 0,008413822 | 2,075006665 | 0,062919794 | WT -MAGE-D1 KO |
| **ADO** | Q96SZ5 | 2-aminoethanethiol dioxygenase | -1,018138591 | 0,042778223 | 1,368777257 | 0,147204721 | WT -MAGE-D1 KO |
| **CNPY3** | Q9BT09 | Protein canopy homolog 3 | -1,018121577 | 0,061840286 | 1,208728512 | 0,181670165 | WT -MAGE-D1 KO |
| **CD63** | P08962 | CD63 antigen | -1,016664642 | 0,019963961 | 1,699753296 | 0,09932534 | WT -MAGE-D1 KO |
| **PALS2** | Q9NZW5 | Protein PALS2 | -1,008248534 | 0,033687088 | 1,472536532 | 0,129320664 | WT -MAGE-D1 KO |
| **MRPS22** | P82650 | 28S ribosomal protein S22, mitochondrial | -1,007830313 | 0,025529361 | 1,592960059 | 0,113174297 | WT -MAGE-D1 KO |
| **DDR2** | Q16832 | Discoidin domain-containing receptor 2 | -1,003134414 | 0,008053091 | 2,094037419 | 0,061043946 | WT -MAGE-D1 KO |
| **FLG2** | Q5D862 | Filaggrin-2 | -1,001044954 | 0,028625056 | 1,543253648 | 0,120262655 | WT -MAGE-D1 KO |
| **DGLUCY** | Q7Z3D6 | D-glutamate cyclase, mitochondrial | -1,000982512 | 0,006856283 | 2,163911281 | 0,056648053 | WT -MAGE-D1 KO |

**Supplementary table 5:** Proteomics with reported hits and candidates of cell surface protein enrichment from differentiated SH-SY5Y WT versus MAGE-D1 KO cells. Related to Supplementary Figure 2G.

| Gene | Protein ID | Protein Description | log2(FC) | -log10(pvalue) | Comparison |
|---|---|---|---|---|---|
| **HIST1H2BP;HIST** | [illegible] | Histone H2B type 1-P;Histone H2B type 1-K;H | 2,06 | 2,06 | Cocaine - Saline |
| **HNRNPA2B1** | O88569 | Heterogeneous nuclear ribonucleoproteins A2 | 2,37 | 1,81 | Cocaine - Saline |
| **Neurabin-1** | H3BJD6 | | 2,39 | 1,72 | Cocaine - Saline |
| **HIST1H4A** | P62806 | Histone H4 | 2,45 | 1,99 | Cocaine - Saline |
| **Neurabin-2** | Q6R891 | Neurabin-2 | 2,52 | 1,73 | Cocaine - Saline |
| **IGHM** | P01872 | Ig mu chain C region | 2,58 | 1,62 | Cocaine - Saline |
| **IGHG3** | A0A075B5P5;A0A1Y7VJN6;P03987 | Ig gamma-3 chain C region | 3,14 | 3,88 | Cocaine - Saline |
| **HIST2H2AC;HIST** | [illegible] | Histone H2A type 2-C;Histone H2A type 2-A;H | 3,15 | 2,66 | Cocaine - Saline |
| **MYO5A** | Q99104;E9Q1F5;P21271 | Unconventional myosin-Va | 3,22 | 1,97 | Cocaine - Saline |
| **YWHAH** | P68510 | 14-3-3 protein eta | 3,48 | 5,44 | Cocaine - Saline |
| **DBN1** | Q9QXS6 | Drebrin | 3,52 | 1,78 | Cocaine - Saline |
| **CORO2B** | Q8BH44 | Coronin-2B | 3,80 | 1,78 | Cocaine - Saline |
| **IGHV6-6;IGHV6-7** | [illegible] | Ig heavy chain V-III region ABE-47N;Ig heavy | 4,52 | 2,58 | Cocaine - Saline |
| **RPS3** | P62908 | 40S ribosomal protein S3 | 0,62 | 1,26 | Cocaine - Saline |
| **RPL6** | P47911 | 60S ribosomal protein L6 | 0,66 | 1,22 | Cocaine - Saline |
| **PSD-95** | Q62108 | Disks large homolog 4 | 0,71 | 1,18 | Cocaine - Saline |
| **RPL18** | P35980 | 60S ribosomal protein L18 | 0,73 | 1,62 | Cocaine - Saline |
| **SNAP25** | P60879 | Synaptosomal-associated protein 25 | 0,73 | 1,29 | Cocaine - Saline |
| **RPS4X;GM15013** | P62702;V9GWY0 | 40S ribosomal protein S4, X isoform;40S ribos | 0,81 | 2,84 | Cocaine - Saline |
| **RPL26** | P61255 | 60S ribosomal protein L26 | 0,81 | 1,48 | Cocaine - Saline |
| **NT5DC3** | Q3UHB1;A0A1W2P799 | 5-nucleotidase domain-containing protein 3 | 0,84 | 1,06 | Cocaine - Saline |
| **TCP1** | P11983 | T-complex protein 1 subunit alpha | 0,85 | 1,05 | Cocaine - Saline |
| **DOCK7** | Q8R1A4;Q8VDR9 | Dedicator of cytokinesis protein 7 | 0,86 | 1,28 | Cocaine - Saline |
| **RPL3** | P27659 | 60S ribosomal protein L3 | 0,86 | 2,47 | Cocaine - Saline |
| **RPS6** | P62754 | 40S ribosomal protein S6 | 0,88 | 2,81 | Cocaine - Saline |
| **LUC7L** | Q9CYI4 | Putative RNA-binding protein Luc7-like 1 | 0,96 | 1,30 | Cocaine - Saline |
| **RPS8** | P62242 | 40S ribosomal protein S8 | 0,96 | 3,34 | Cocaine - Saline |
| **MAP6** | Q7TSJ2 | Microtubule-associated protein 6 | 1,02 | 4,96 | Cocaine - Saline |
| **SNAP47** | Q8R570 | Synaptosomal-associated protein 47 | 1,05 | 1,59 | Cocaine - Saline |
| **TRY10** | Q792Z1 | | 1,06 | 1,35 | Cocaine - Saline |
| **RPS20** | P60867 | 40S ribosomal protein S20 | 1,20 | 1,87 | Cocaine - Saline |
| **DMXL2** | Q8BPN8 | DmX-like protein 2 | 1,21 | 2,34 | Cocaine - Saline |
| **MPRIP** | P97434 | Myosin phosphatase Rho-interacting protein | 1,24 | 1,30 | Cocaine - Saline |
| **HSPA5** | P20029 | 78 kDa glucose-regulated protein | 1,27 | 1,15 | Cocaine - Saline |
| **RPL27** | P61358 | 60S ribosomal protein L27 | 1,33 | 3,98 | Cocaine - Saline |
| **ACTN4** | P57780;Q9JI91;O88990 | Alpha-actinin-4 | 1,48 | 1,06 | Cocaine - Saline |
| **BASP1** | Q91XV3 | Brain acid soluble protein 1 | 1,50 | 1,19 | Cocaine - Saline |
| **ATP2A2** | O55143;Q8R429;Q64518 | Sarcoplasmic/endoplasmic reticulum calcium | 1,50 | 2,29 | Cocaine - Saline |
| **ARF1;ARF3;ARF2** | P84078;P61205;Q8BSL7;P84084 | ADP-ribosylation factor 1;ADP-ribosylation fac | 1,51 | 1,35 | Cocaine - Saline |
| **RPL38** | Q9JJI8 | 60S ribosomal protein L38 | 1,55 | 1,56 | Cocaine - Saline |
| **PC** | Q05920 | Pyruvate carboxylase, mitochondrial | 1,55 | 1,42 | Cocaine - Saline |
| **STIP1** | Q60864 | Stress-induced-phosphoprotein 1 | 1,55 | 2,72 | Cocaine - Saline |
| **MAPT** | P10637 | Microtubule-associated protein tau | 1,57 | 1,30 | Cocaine - Saline |
| **G3BP2** | P97379 | Ras GTPase-activating protein-binding protei | 1,62 | 1,93 | Cocaine - Saline |
| **EWSR1** | Q61545 | RNA-binding protein EWS | 1,64 | 1,17 | Cocaine - Saline |
| **PGK1** | P09411;P09041 | Phosphoglycerate kinase 1 | 1,67 | 1,18 | Cocaine - Saline |
| **ACTB;ACTG1** | P60710;P63260;V9GXQ2 | Actin, cytoplasmic 1;Actin, cytoplasmic 1, N-te | 1,67 | 1,06 | Cocaine - Saline |
| **EPB41L3** | Q9WV92 | Band 4.1-like protein 3;Band 4.1-like protein 3 | 1,68 | 1,47 | Cocaine - Saline |
| **CKMT1** | P30275 | Creatine kinase U-type, mitochondrial | 1,76 | 1,86 | Cocaine - Saline |
| **RPL35A** | O55142 | 60S ribosomal protein L35a | 1,80 | 1,74 | Cocaine - Saline |
| **ARPC1A** | Q9R0Q6 | Actin-related protein 2/3 complex subunit 1A | 1,85 | 1,99 | Cocaine - Saline |
| **FXR1;FMR1** | Q61584;P35922 | Fragile X mental retardation syndrome-related | 1,91 | 1,65 | Cocaine - Saline |
| **SYN2** | Q64332 | Synapsin-2 | 1,94 | 1,12 | Cocaine - Saline |
| **CDC42BPB** | Q7TT50;Q3UU96;Q80UW5 | Serine/threonine-protein kinase MRCK beta | 1,95 | 2,02 | Cocaine - Saline |
| **ACTR3** | Q99JY9 | Actin-related protein 3 | 1,96 | 1,13 | Cocaine - Saline |
| **MAATS1** | Q8BRC6 | Protein MAATS1 | -9,08 | 2,24 | Cocaine - Saline |
| **PLP1** | P60202 | Myelin proteolipid protein | -4,54 | 2,88 | Cocaine - Saline |
| **ALDOA;ALDOAR** | P05064;A6ZI46;A6ZI47 | Fructose-bisphosphate aldolase A;Fructose-b | -4,16 | 2,48 | Cocaine - Saline |
| **SLC1A2** | P43006 | Excitatory amino acid transporter 2 | -3,36 | 2,41 | Cocaine - Saline |
| **SLC25A5** | P51881 | ADP/ATP translocase 2;ADP/ATP translocase | -3,34 | 1,65 | Cocaine - Saline |
| **FAM171A2** | A2A699 | Protein FAM171A2 | -3,17 | 2,88 | Cocaine - Saline |
| **JUP** | Q02257 | Junction plakoglobin | -3,05 | 2,29 | Cocaine - Saline |
| **C4B** | P01029 | Complement C4-B;Complement C4 beta chai | -2,83 | 1,85 | Cocaine - Saline |
| **AP3D1** | O54774 | AP-3 complex subunit delta-1 | -2,69 | 2,65 | Cocaine - Saline |
| **CMAS** | Q99KK2 | N-acylneuraminate cytidylyltransferase | -2,55 | 1,84 | Cocaine - Saline |
| **CTNNA2** | Q61301;P26231 | Catenin alpha-2 | -1,97 | 1,36 | Cocaine - Saline |
| **GLUD1** | P26443 | Glutamate dehydrogenase 1, mitochondrial | -1,95 | 2,18 | Cocaine - Saline |
| **SLC25A11** | Q9CR62 | Mitochondrial 2-oxoglutarate/malate carrier pr | -1,93 | 1,69 | Cocaine - Saline |
| **TUBB3** | [illegible] | Tubulin beta-3 chain | -1,92 | 1,38 | Cocaine - Saline |

| | | | | | |
|---|---|---|---|---|---|
| **MYL1;MYL3** | P05977;P09542 | Myosin light chain 1/3, skeletal muscle isoform | -1,89 | 1,25 | Cocaine - Saline |
| **SLC25A12** | Q8BH59 | Calcium-binding mitochondrial carrier protein | -1,83 | 1,09 | Cocaine - Saline |
| **HBA** | P01942 | Hemoglobin subunit alpha | -1,81 | 1,08 | Cocaine - Saline |
| **TPD52L2** | Q9CYZ2 | Tumor protein D54 | -1,80 | 1,18 | Cocaine - Saline |
| **IMMT** | Q8CAQ8 | MICOS complex subunit Mic60 | -1,79 | 1,25 | Cocaine - Saline |
| **EIF2S3X;EIF2S3Y** | Q9Z0N1;Q9Z0N2 | Eukaryotic translation initiation factor 2 subuni | -1,73 | 1,48 | Cocaine - Saline |
| **CNP** | P16330 | 2,3-cyclic-nucleotide 3-phosphodiesterase | -1,69 | 3,31 | Cocaine - Saline |
| **ELAVL2** | Q60899;Q60900 | ELAV-like protein 2 | -1,68 | 1,58 | Cocaine - Saline |
| **TNR** | Q8BYI9 | Tenascin-R | -1,62 | 1,13 | Cocaine - Saline |
| **TPM3** | P21107 | Tropomyosin alpha-3 chain | -1,59 | 1,62 | Cocaine - Saline |
| **ATAD3** | Q925I1 | ATPase family AAA domain-containing protein | -1,51 | 1,19 | Cocaine - Saline |
| **RPL19** | P84099 | 60S ribosomal protein L19 | -1,49 | 1,29 | Cocaine - Saline |
| **GNB1;GNB2** | P62874;P62880;P29387;Q61011 | Guanine nucleotide-binding protein G(I)/G(S)/ | -1,45 | 1,26 | Cocaine - Saline |
| **UQCRC2** | Q9DB77 | Cytochrome b-c1 complex subunit 2, mitochor | -1,44 | 1,04 | Cocaine - Saline |
| **C1QB** | P14106 | Complement C1q subcomponent subunit B | -1,22 | 1,01 | Cocaine - Saline |
| **RPS25** | P62852 | 40S ribosomal protein S25 | -1,21 | 2,20 | Cocaine - Saline |
| **ATP5A1** | Q03265 | ATP synthase subunit alpha, mitochondrial | -1,21 | 1,24 | Cocaine - Saline |
| **ATP2B1** | G5E829 | Plasma membrane calcium-transporting ATPa | -1,18 | 1,66 | Cocaine - Saline |
| **TUBB4B** | P68372 | Tubulin beta-4B chain | -1,01 | 1,03 | Cocaine - Saline |
| **CAMK2A** | P11798 | Calcium/calmodulin-dependent protein kinase | -0,96 | 3,81 | Cocaine - Saline |
| **TUBA1B;TUBA1C** | P05213;P68373;Q9JJZ2;A0A2R8VHF3;Q3UX10 | Tubulin alpha-1B chain;Tubulin alpha-1C cha | -0,92 | 1,62 | Cocaine - Saline |
| **SLC25A22** | Q9D6M3;Q9DB41 | Mitochondrial glutamate carrier 1 | -0,66 | 1,21 | Cocaine - Saline |

**Supplementary table 6:** Proteomics with reported hits and candidates of endogenous MAGE-D1 immunoprecipitation from from thalamus of control mice treated with saline versus mice treated with cocaine. Related to Supplementary Figure 3A.

| Gene | Protein ID | Protein Description | log2(FC) | pvalue | -log10(pvalue) | fdr | Comparison |
|---|---|---|---|---|---|---|---|
| PJA2 | O43164\|O43164-2 | E3 ubiquitin-protein ligase Praja-2 OS=H | 3,346210134 | 1,86219E-09 | 8,729975914 | 1,10661E-06 | (GFP_MAGED1_HD - GFP)-(GFP_MAGED1_UN - GFP) |
| MAGE-D1_MHD | P1711_HD | NA | 4,691877033 | 2,41872E-08 | 7,616414837 | 3,63749E-06 | (GFP_MAGED1_HD - GFP)-(GFP_MAGED1_UN - GFP) |
| NDRG1 | Q92597 | Protein NDRG1 OS=Homo sapiens GN= | 2,536116803 | 2,86432E-08 | 7,542978587 | 3,78249E-06 | (GFP_MAGED1_HD - GFP)-(GFP_MAGED1_UN - GFP) |
| SMC5 | Q8IY18 | Structural maintenance of chromosomes | 2,129591419 | 3,9839E-08 | 7,399691939 | 4,31573E-06 | (GFP_MAGED1_HD - GFP)-(GFP_MAGED1_UN - GFP) |
| RPS27A | P62979 | Ubiquitin-40S ribosomal protein S27a O | 2,141997189 | 4,64786E-08 | 7,332747362 | 4,41918E-06 | (GFP_MAGED1_HD - GFP)-(GFP_MAGED1_UN - GFP) |
| NSMCE3 | Q96MG7 | Non-structural maintenance of chromoso | 2,864395662 | 1,15891E-07 | 6,935950383 | 8,34766E-06 | (GFP_MAGED1_HD - GFP)-(GFP_MAGED1_UN - GFP) |
| SMC6 | Q96SB8\|Q96SB8-2 | Structural maintenance of chromosomes | 1,805253119 | 1,86179E-07 | 6,730068711 | 1,10637E-05 | (GFP_MAGED1_HD - GFP)-(GFP_MAGED1_UN - GFP) |
| HUWE1 | Q7Z6Z7\|Q7Z6Z7-2\|Q7Z6Z7-3 | E3 ubiquitin-protein ligase HUWE1 OS= | 2,822973876 | 1,92593E-07 | 6,715360417 | 1,11657E-05 | (GFP_MAGED1_HD - GFP)-(GFP_MAGED1_UN - GFP) |
| AP1S2 | P56377 | AP-1 complex subunit sigma-2 OS=Hom | 2,405256953 | 2,93073E-07 | 6,533024766 | 1,45132E-05 | (GFP_MAGED1_HD - GFP)-(GFP_MAGED1_UN - GFP) |
| PJA1 | Q8NG27\|Q8NG27-3 | E3 ubiquitin-protein ligase Praja-1 OS=H | 2,748579737 | 3,6139E-07 | 6,4420242 | 1,60679E-05 | (GFP_MAGED1_HD - GFP)-(GFP_MAGED1_UN - GFP) |
| PRKDC | P78527 | DNA-dependent protein kinase catalytic | 1,788326807 | 3,77287E-07 | 6,423327994 | 1,60679E-05 | (GFP_MAGED1_HD - GFP)-(GFP_MAGED1_UN - GFP) |
| WRNIP1 | Q96S55\|Q96S55-2 | ATPase WRNIP1 OS=Homo sapiens GN | 2,123701996 | 4,06734E-07 | 6,390689065 | 1,61135E-05 | (GFP_MAGED1_HD - GFP)-(GFP_MAGED1_UN - GFP) |
| PNPLA6 | Q8IY17\|Q8IY17-2\|Q8IY17-3\|( | Neuropathy target esterase OS=Homo s | 1,434794721 | 1,20079E-06 | 5,92053404 | 3,65932E-05 | (GFP_MAGED1_HD - GFP)-(GFP_MAGED1_UN - GFP) |
| NSMCE4A | Q9NXX6 | Non-structural maintenance of chromoso | 2,162243953 | 1,5577E-06 | 5,807515169 | 4,40793E-05 | (GFP_MAGED1_HD - GFP)-(GFP_MAGED1_UN - GFP) |
| DNAJC3 | Q13217 | DnaJ homolog subfamily C member 3 O | 1,698248059 | 1,73459E-06 | 5,760802329 | 4,61599E-05 | (GFP_MAGED1_HD - GFP)-(GFP_MAGED1_UN - GFP) |
| PCK2 | Q16822 | Phosphoenolpyruvate carboxykinase [G | 1,880319574 | 1,74775E-06 | 5,757521381 | 4,61599E-05 | (GFP_MAGED1_HD - GFP)-(GFP_MAGED1_UN - GFP) |
| ALDH1B1 | A0A0U1RQK9\|P30837 | Aldehyde dehydrogenase X, mitochondr | 1,532965793 | 1,80321E-06 | 5,74395444 | 4,71013E-05 | (GFP_MAGED1_HD - GFP)-(GFP_MAGED1_UN - GFP) |
| DAAM1 | Q9Y4D1\|Q9Y4D1-2 | Disheveled-associated activator of morp | 1,335606761 | 1,86693E-06 | 5,72887282 | 4,82357E-05 | (GFP_MAGED1_HD - GFP)-(GFP_MAGED1_UN - GFP) |
| VWA8 | A3KMH1\|A3KMH1-3 | von Willebrand factor A domain-containi | 1,303833787 | 2,06436E-06 | 5,685213623 | 5,11145E-05 | (GFP_MAGED1_HD - GFP)-(GFP_MAGED1_UN - GFP) |
| TAX1BP1 | Q86VP1-2 | Isoform 2 of Tax1-binding protein 1 OS= | 1,605511129 | 2,27618E-06 | 5,642792521 | 5,57783E-05 | (GFP_MAGED1_HD - GFP)-(GFP_MAGED1_UN - GFP) |
| VCP | P55072 | Transitional endoplasmic reticulum ATP | 1,61908833 | 2,38493E-06 | 5,622524817 | 5,74357E-05 | (GFP_MAGED1_HD - GFP)-(GFP_MAGED1_UN - GFP) |
| EPS15 | B1AUU8\|P42566 | Epidermal growth factor receptor substra | 1,98808768 | 2,41631E-06 | 5,616847351 | 5,74357E-05 | (GFP_MAGED1_HD - GFP)-(GFP_MAGED1_UN - GFP) |
| AIFM1 | O95831\|O95831-3 | Apoptosis-inducing factor 1, mitochondri | 1,308790185 | 2,93491E-06 | 5,532405561 | 6,70796E-05 | (GFP_MAGED1_HD - GFP)-(GFP_MAGED1_UN - GFP) |
| LPCAT2 | Q7L5N7 | Lysophosphatidylcholine acyltransferase | 1,317764956 | 3,11636E-06 | 5,506352341 | 7,05485E-05 | (GFP_MAGED1_HD - GFP)-(GFP_MAGED1_UN - GFP) |
| ASPH | Q12797 | Aspartyl/asparaginyl beta-hydroxylase C | 1,328045871 | 3,77093E-06 | 5,423551563 | 8,29954E-05 | (GFP_MAGED1_HD - GFP)-(GFP_MAGED1_UN - GFP) |
| PEX14 | O75381\|O75381-2 | Peroxisomal membrane protein PEX14 ( | 1,4918062 | 4,18621E-06 | 5,378178754 | 9,12902E-05 | (GFP_MAGED1_HD - GFP)-(GFP_MAGED1_UN - GFP) |
| ARMC8 | Q8IUR7\|Q8IUR7-2\|Q8IUR7-6 | Armadillo repeat-containing protein 8 OS | 1,397087022 | 4,32499E-06 | 5,364014647 | 9,26172E-05 | (GFP_MAGED1_HD - GFP)-(GFP_MAGED1_UN - GFP) |
| PIK3R1 | P27986\|P27986-4 | Phosphatidylinositol 3-kinase regulatory | 1,216710623 | 5,79084E-06 | 5,2372583 | 0,000114707 | (GFP_MAGED1_HD - GFP)-(GFP_MAGED1_UN - GFP) |
| BAT3\|BAG6 | A0A024RCR6\|A0A0G2JK23\| | HLA-B associated transcript 3, isoform C | 1,281635857 | 5,94193E-06 | 5,226072355 | 0,000116727 | (GFP_MAGED1_HD - GFP)-(GFP_MAGED1_UN - GFP) |
| VPS4A | Q9UN37 | Vacuolar protein sorting-associated prot | 1,260285456 | 7,24414E-06 | 5,140013331 | 0,000136661 | (GFP_MAGED1_HD - GFP)-(GFP_MAGED1_UN - GFP) |
| ASNA1 | A0A087WXS7\|O43681 | ATPase ASNA1 OS=Homo sapiens GN= | 1,512031759 | 8,60585E-06 | 5,065206331 | 0,000156153 | (GFP_MAGED1_HD - GFP)-(GFP_MAGED1_UN - GFP) |
| FAM63A | Q8N5J2 | Protein FAM63A OS=Homo sapiens GN | 1,22687334 | 8,83453E-06 | 5,053816651 | 0,000157892 | (GFP_MAGED1_HD - GFP)-(GFP_MAGED1_UN - GFP) |
| DNAJB1 | P25685 | DnaJ homolog subfamily B member 1 O | 1,56123859 | 9,12453E-06 | 5,039789564 | 0,000158713 | (GFP_MAGED1_HD - GFP)-(GFP_MAGED1_UN - GFP) |
| MYO6 | A0A0A0MRM8\|Q9UM54-6 | Unconventional myosin-VI OS=Homo sa | 1,280554769 | 9,74289E-06 | 5,011312199 | 0,000161714 | (GFP_MAGED1_HD - GFP)-(GFP_MAGED1_UN - GFP) |
| ARMC6 | B4E1N1\|Q6NXE6\|Q6NXE6-2 | Armadillo repeat-containing protein 6 OS | 1,205428725 | 9,74727E-06 | 5,011117022 | 0,000161714 | (GFP_MAGED1_HD - GFP)-(GFP_MAGED1_UN - GFP) |
| MAGED4 | Q96JG8\|Q96JG8-2\|Q96JG8- | Melanoma-associated antigen D4 OS=H | 1,720885982 | 1,03687E-05 | 4,984274249 | 0,000167663 | (GFP_MAGED1_HD - GFP)-(GFP_MAGED1_UN - GFP) |
| FTSJ1 | Q9UET6\|Q9UET6-2 | Putative tRNA (cytidine(32)/guanosine(3 | 1,244484489 | 1,04933E-05 | 4,979085967 | 0,000168459 | (GFP_MAGED1_HD - GFP)-(GFP_MAGED1_UN - GFP) |
| AP1G2 | O75843 | AP-1 complex subunit gamma-like 2 OS | 2,491941155 | 1,12924E-05 | 4,947211993 | 0,000175449 | (GFP_MAGED1_HD - GFP)-(GFP_MAGED1_UN - GFP) |
| TUBB4A | P04350 | Tubulin beta-4A chain OS=Homo sapien | 1,075522173 | 1,12931E-05 | 4,947187324 | 0,000175449 | (GFP_MAGED1_HD - GFP)-(GFP_MAGED1_UN - GFP) |
| AMOT | Q4VCS5 | Angiomotin OS=Homo sapiens GN=AMC | 1,29286009 | 1,17063E-05 | 4,931582064 | 0,000180687 | (GFP_MAGED1_HD - GFP)-(GFP_MAGED1_UN - GFP) |
| NPLOC4 | Q8TAT6\|Q8TAT6-2 | Nuclear protein localization protein 4 ho | 1,281057744 | 1,23628E-05 | 4,907883778 | 0,000188374 | (GFP_MAGED1_HD - GFP)-(GFP_MAGED1_UN - GFP) |
| ALCAM | Q13740-2 | Isoform 2 of CD166 antigen OS=Homo s | 1,458986547 | 1,31119E-05 | 4,882333379 | 0,00019726 | (GFP_MAGED1_HD - GFP)-(GFP_MAGED1_UN - GFP) |
| TRUB2 | O95900\|O95900-2 | Probable tRNA pseudouridine synthase | 1,880561866 | 1,35219E-05 | 4,868961635 | 0,000200885 | (GFP_MAGED1_HD - GFP)-(GFP_MAGED1_UN - GFP) |
| IRS4 | O14654 | Insulin receptor substrate 4 OS=Homo s | 1,300825455 | 1,70345E-05 | 4,768670605 | 0,0002454 | (GFP_MAGED1_HD - GFP)-(GFP_MAGED1_UN - GFP) |
| HSPD1 | P10809 | 60 kDa heat shock protein, mitochondria | 1,033785616 | 1,78242E-05 | 4,748989248 | 0,000253702 | (GFP_MAGED1_HD - GFP)-(GFP_MAGED1_UN - GFP) |
| RIF1 | Q5UIP0\|Q5UIP0-2 | Telomere-associated protein RIF1 OS=H | 1,005521059 | 1,97961E-05 | 4,703420136 | 0,000276796 | (GFP_MAGED1_HD - GFP)-(GFP_MAGED1_UN - GFP) |
| TUBB | P07437\|Q5JP53 | Tubulin beta chain OS=Homo sapiens G | 1,04559695 | 2,08721E-05 | 4,680434712 | 0,000290134 | (GFP_MAGED1_HD - GFP)-(GFP_MAGED1_UN - GFP) |
| AP1AR | Q63HQ0 | AP-1 complex-associated regulatory pro | 1,27416648 | 2,12608E-05 | 4,672420483 | 0,000292121 | (GFP_MAGED1_HD - GFP)-(GFP_MAGED1_UN - GFP) |
| TUFM | P49411 | Elongation factor Tu, mitochondrial OS= | 1,164792545 | 2,35789E-05 | 4,627475925 | 0,000311373 | (GFP_MAGED1_HD - GFP)-(GFP_MAGED1_UN - GFP) |
| UBL4A | P11441 | Ubiquitin-like protein 4A OS=Homo sapi | 1,295271292 | 2,42043E-05 | 4,616108044 | 0,000317865 | (GFP_MAGED1_HD - GFP)-(GFP_MAGED1_UN - GFP) |
| CLPB | Q9H078\|Q9H078-2 | Caseinolytic peptidase B protein homolo | 1,031931541 | 2,93051E-05 | 4,533056962 | 0,000375374 | (GFP_MAGED1_HD - GFP)-(GFP_MAGED1_UN - GFP) |
| CAND2 | O75155 | Cullin-associated NEDD8-dissociated p | 1,012781834 | 2,97757E-05 | 4,526137796 | 0,000376473 | (GFP_MAGED1_HD - GFP)-(GFP_MAGED1_UN - GFP) |
| STUB1 | Q9UNE7 | E3 ubiquitin-protein ligase CHIP OS=Ho | 1,125981733 | 3,36043E-05 | 4,473604971 | 0,00041985 | (GFP_MAGED1_HD - GFP)-(GFP_MAGED1_UN - GFP) |
| GTF2B | Q00403 | Transcription initiation factor IIB OS=Hor | 1,002657968 | 3,38051E-05 | 4,471018027 | 0,00041985 | (GFP_MAGED1_HD - GFP)-(GFP_MAGED1_UN - GFP) |
| VARS2 | A0A0A0MTG1\|A0A0G2JIA5\|/ | Valine--tRNA ligase, mitochondrial OS=I | 1,111906985 | 3,65781E-05 | 4,43677882 | 0,000443603 | (GFP_MAGED1_HD - GFP)-(GFP_MAGED1_UN - GFP) |
| KCT2 | Q8NC54 | Keratinocyte-associated transmembrane | 1,084609368 | 3,71352E-05 | 4,430213744 | 0,000448073 | (GFP_MAGED1_HD - GFP)-(GFP_MAGED1_UN - GFP) |
| ARFGEF1 | E5RIF2\|Q9Y6D6 | Brefeldin A-inhibited guanine nucleotide | 1,031073026 | 3,88796E-05 | 4,410278486 | 0,000464406 | (GFP_MAGED1_HD - GFP)-(GFP_MAGED1_UN - GFP) |
| H0YC42 | H0YC42 | Uncharacterized protein OS=Homo sapi | 1,053078018 | 4,02581E-05 | 4,395147211 | 0,000476087 | (GFP_MAGED1_HD - GFP)-(GFP_MAGED1_UN - GFP) |
| SYMPK | Q92797 | Symplekin OS=Homo sapiens GN=SYM | 1,045941177 | 4,25409E-05 | 4,371193281 | 0,000498127 | (GFP_MAGED1_HD - GFP)-(GFP_MAGED1_UN - GFP) |
| DNM2 | P50570\|P50570-2\|P50570-3 | Dynamin-2 OS=Homo sapiens GN=DNM | 1,048948065 | 4,62501E-05 | 4,334886856 | 0,000531095 | (GFP_MAGED1_HD - GFP)-(GFP_MAGED1_UN - GFP) |
| MICU2 | A0A0A0MTD5\|Q8IYU8 | Calcium uptake protein 2, mitochondrial | 1,065292931 | 5,19735E-05 | 4,284218328 | 0,000577294 | (GFP_MAGED1_HD - GFP)-(GFP_MAGED1_UN - GFP) |
| FANCI | F8W7R3\|Q9NVI1\|Q9NVI1-1\|( | Fanconi anemia group I protein OS=Hor | 1,062985753 | 5,44958E-05 | 4,263636783 | 0,000591491 | (GFP_MAGED1_HD - GFP)-(GFP_MAGED1_UN - GFP) |
| AP1G1 | O43747\|O43747-2 | AP-1 complex subunit gamma-1 OS=Ho | 1,263378109 | 6,30339E-05 | 4,200425569 | 0,000651442 | (GFP_MAGED1_HD - GFP)-(GFP_MAGED1_UN - GFP) |
| SPG7 | Q9UQ90 | Paraplegin OS=Homo sapiens GN=SPG | 1,161143619 | 6,68858E-05 | 4,174666156 | 0,000685291 | (GFP_MAGED1_HD - GFP)-(GFP_MAGED1_UN - GFP) |
| PIK3R2 | O00459 | Phosphatidylinositol 3-kinase regulatory | 1,065315772 | 7,90532E-05 | 4,102080312 | 0,000786232 | (GFP_MAGED1_HD - GFP)-(GFP_MAGED1_UN - GFP) |
| AHCTF1 | Q8WYP5\|Q8WYP5-2\|Q8WYF | Protein ELYS OS=Homo sapiens GN=AI | 1,157066897 | 8,79623E-05 | 4,055703484 | 0,000849945 | (GFP_MAGED1_HD - GFP)-(GFP_MAGED1_UN - GFP) |
| AFG3L2 | Q9Y4W6 | AFG3-like protein 2 OS=Homo sapiens ( | 1,010147511 | 9,36503E-05 | 4,028490942 | 0,00088336 | (GFP_MAGED1_HD - GFP)-(GFP_MAGED1_UN - GFP) |
| UBR5 | E7EMW7\|O95071\|O95071-2 | E3 ubiquitin-protein ligase UBR5 OS=H | 1,327239085 | 9,79061E-05 | 4,009190327 | 0,000909073 | (GFP_MAGED1_HD - GFP)-(GFP_MAGED1_UN - GFP) |
| SGPL1 | O95470 | Sphingosine-1-phosphate lyase 1 OS=H | 1,009874663 | 9,95117E-05 | 4,00212568 | 0,000920387 | (GFP_MAGED1_HD - GFP)-(GFP_MAGED1_UN - GFP) |
| TAP1 | A0A140T9T7\|Q03518 | Antigen peptide transporter 1 OS=Homo | 1,161134647 | 0,000103452 | 3,985262217 | 0,000942164 | (GFP_MAGED1_HD - GFP)-(GFP_MAGED1_UN - GFP) |
| NOP58 | Q9Y2X3 | Nucleolar protein 58 OS=Homo sapiens | 1,048333902 | 0,000106674 | 3,971940494 | 0,000964124 | (GFP_MAGED1_HD - GFP)-(GFP_MAGED1_UN - GFP) |
| SLC1A5 | Q15758 | Neutral amino acid transporter B(0) OS= | 1,161478399 | 0,000118668 | 3,925666327 | 0,001052514 | (GFP_MAGED1_HD - GFP)-(GFP_MAGED1_UN - GFP) |
| NSMCE2 | E5RFJ1\|Q96MF7 | E3 SUMO-protein ligase NSE2 OS=Hom | 1,493127522 | 0,000119662 | 3,922044625 | 0,001057383 | (GFP_MAGED1_HD - GFP)-(GFP_MAGED1_UN - GFP) |
| NSMCE1 | Q8WV22 | Non-structural maintenance of chromoso | 1,254977269 | 0,000165301 | 3,781723779 | 0,001338188 | (GFP_MAGED1_HD - GFP)-(GFP_MAGED1_UN - GFP) |
| PGRMC1 | O00264 | Membrane-associated progesterone rec | 1,035418038 | 0,000197412 | 3,704627215 | 0,001513702 | (GFP_MAGED1_HD - GFP)-(GFP_MAGED1_UN - GFP) |
| CNNM3 | Q8NE01 | Metal transporter CNNM3 OS=Homo sap | 1,188465718 | 0,000214109 | 3,669365612 | 0,001615671 | (GFP_MAGED1_HD - GFP)-(GFP_MAGED1_UN - GFP) |
| POMGNT1 | Q8WZA1\|Q8WZA1-2 | Protein O-linked-mannose beta-1,2-N-ac | 1,11456625 | 0,000217124 | 3,663292036 | 0,001628088 | (GFP_MAGED1_HD - GFP)-(GFP_MAGED1_UN - GFP) |
| CECR5 | Q9BXW7\|Q9BXW7-2 | Cat eye syndrome critical region protein | 1,171291392 | 0,000222219 | 3,653218958 | 0,001661052 | (GFP_MAGED1_HD - GFP)-(GFP_MAGED1_UN - GFP) |
| C1orf112 | Q9NSG2\|Q9NSG2-2\|Q9NSG | Uncharacterized protein C1orf112 OS=H | 1,611133564 | 0,000261843 | 3,581959542 | 0,001874699 | (GFP_MAGED1_HD - GFP)-(GFP_MAGED1_UN - GFP) |
| CCDC134 | Q9H6E4 | Coiled-coil domain-containing protein 13 | 1,072654163 | 0,000291461 | 3,535419301 | 0,002043667 | (GFP_MAGED1_HD - GFP)-(GFP_MAGED1_UN - GFP) |
| UFD1L | Q92890\|Q92890-1 | Ubiquitin fusion degradation protein 1 h | 1,553723087 | 0,000339838 | 3,468728009 | 0,00230141 | (GFP_MAGED1_HD - GFP)-(GFP_MAGED1_UN - GFP) |
| TMEM106B | Q9NUM4 | Transmembrane protein 106B OS=Hom | 1,052444311 | 0,00036907 | 3,432891731 | 0,002436885 | (GFP_MAGED1_HD - GFP)-(GFP_MAGED1_UN - GFP) |
| ATP6V1H | G3V126\|Q9UI12 | ATPase, H+ transporting, lysosomal 50/5 | 1,010458647 | 0,000498118 | 3,302668028 | 0,003044675 | (GFP_MAGED1_HD - GFP)-(GFP_MAGED1_UN - GFP) |
| TNPO1 | Q92973\|Q92973-2\|Q92973-3 | Transportin-1 OS=Homo sapiens GN=TN | 1,151442633 | 0,00051176 | 3,290933842 | 0,003056415 | (GFP_MAGED1_HD - GFP)-(GFP_MAGED1_UN - GFP) |
| EPS15L1 | M0R165\|Q9UBC2\|Q9UBC2-2 | Epidermal growth factor receptor substra | 1,146378259 | 0,000716418 | 3,144833538 | 0,003997478 | (GFP_MAGED1_HD - GFP)-(GFP_MAGED1_UN - GFP) |
| SLC1A4 | P43007 | Neutral amino acid transporter A OS=Ho | 1,034207546 | 0,000957983 | 3,018642301 | 0,004886534 | (GFP_MAGED1_HD - GFP)-(GFP_MAGED1_UN - GFP) |
| MTDH | E5RJU9\|Q86UE4 | Protein LYRIC OS=Homo sapiens GN=M | 1,024114205 | 0,001685842 | 2,773183129 | 0,007450432 | (GFP_MAGED1_HD - GFP)-(GFP_MAGED1_UN - GFP) |
| GALNT1 | Q10472 | Polypeptide N-acetylgalactosaminyltran | 1,101784192 | 0,001884397 | 2,724827533 | 0,008089969 | (GFP_MAGED1_HD - GFP)-(GFP_MAGED1_UN - GFP) |
| LRPPRC | P42704 | Leucine-rich PPR motif-containing prote | 1,010432255 | 0,002170465 | 2,663447204 | 0,008925944 | (GFP_MAGED1_HD - GFP)-(GFP_MAGED1_UN - GFP) |
| SARS2 | M0QWZ7\|Q9NP81 | Serine--tRNA ligase, mitochondrial OS= | 1,214143741 | 0,005237402 | 2,280884075 | 0,01726672 | (GFP_MAGED1_HD - GFP)-(GFP_MAGED1_UN - GFP) |
| PRDX1 | A0A0A0MSI0\|Q06830 | Peroxiredoxin-1 (Fragment) OS=Homo s | 1,074916 | 0,05781078 | 1,237991169 | 0,109845104 | (GFP_MAGED1_HD - GFP)-(GFP_MAGED1_UN - GFP) |
| TCAF1 | Q9Y4C2\|Q9Y4C2-2 | TRPM8 channel-associated factor 1 OS= | -4,508986416 | 7,99201E-11 | 10,09734377 | 1,8997E-07 | (GFP_MAGED1_HD - GFP)-(GFP_MAGED1_UN - GFP) |
| USP7 | Q93009 | Ubiquitin carboxyl-terminal hydrolase 7 ( | -3,886323133 | 3,08324E-10 | 9,51099335 | 3,66442E-07 | (GFP_MAGED1_HD - GFP)-(GFP_MAGED1_UN - GFP) |
| KLHL12 | Q53G59\|Q53G59-2 | Kelch-like protein 12 OS=Homo sapiens | -4,300805804 | 9,79767E-10 | 9,008877318 | 7,76302E-07 | (GFP_MAGED1_HD - GFP)-(GFP_MAGED1_UN - GFP) |
| ACAD11 | Q709F0 | Acyl-CoA dehydrogenase family membe | -3,496746936 | 3,01882E-09 | 8,520163092 | 1,26326E-06 | (GFP_MAGED1_HD - GFP)-(GFP_MAGED1_UN - GFP) |
| ACTL8 | Q9H568 | Actin-like protein 8 OS=Homo sapiens G | -3,5624947 | 3,1887E-09 | 8,496386493 | 1,26326E-06 | (GFP_MAGED1_HD - GFP)-(GFP_MAGED1_UN - GFP) |
| MAGE-D1_UNS | P1711_UNS | NA | -5,141487974 | 6,61981E-09 | 8,179154266 | 2,02145E-06 | (GFP_MAGED1_HD - GFP)-(GFP_MAGED1_UN - GFP) |
| MYO5C | Q9NQX4 | Unconventional myosin-Vc OS=Homo sa | -2,950207465 | 6,80335E-09 | 8,167277039 | 2,02145E-06 | (GFP_MAGED1_HD - GFP)-(GFP_MAGED1_UN - GFP) |
| KPNA6 | O60684 | Importin subunit alpha-7 OS=Homo sapi | -2,896911557 | 1,12524E-08 | 7,948753486 | 2,83237E-06 | (GFP_MAGED1_HD - GFP)-(GFP_MAGED1_UN - GFP) |
| PEG10 | A0A087WX23 | Retrotransposon-derived protein PEG10 | -3,17627087 | 1,30103E-08 | 7,885713509 | 2,83237E-06 | (GFP_MAGED1_HD - GFP)-(GFP_MAGED1_UN - GFP) |
| MIB2 | F2Z2L2\|Q96AX9-5 | E3 ubiquitin-protein ligase MIB2 OS=Ho | -2,967349218 | 1,31073E-08 | 7,882486358 | 2,83237E-06 | (GFP_MAGED1_HD - GFP)-(GFP_MAGED1_UN - GFP) |
| DNAJC30 | Q96LL9 | DnaJ homolog subfamily C member 30 ( | -2,668588718 | 1,67345E-08 | 7,776387955 | 3,25105E-06 | (GFP_MAGED1_HD - GFP)-(GFP_MAGED1_UN - GFP) |
| HELZ | J3QS41\|P42694 | Probable helicase with zinc finger doma | -2,442833366 | 1,77802E-08 | 7,750062463 | 3,25105E-06 | (GFP_MAGED1_HD - GFP)-(GFP_MAGED1_UN - GFP) |
| MIB1 | Q86YT6 | E3 ubiquitin-protein ligase MIB1 OS=Ho | -2,96235298 | 2,43868E-08 | 7,612845428 | 3,63749E-06 | (GFP_MAGED1_HD - GFP)-(GFP_MAGED1_UN - GFP) |
| IQGAP3 | Q86VI3 | Ras GTPase-activating-like protein IQGA | -2,78599242 | 2,59716E-08 | 7,58550063 | 3,63749E-06 | (GFP_MAGED1_HD - GFP)-(GFP_MAGED1_UN - GFP) |
| ASB7 | Q9H672 | Ankyrin repeat and SOCS box protein 7 | -3,782536092 | 2,60148E-08 | 7,584779014 | 3,63749E-06 | (GFP_MAGED1_HD - GFP)-(GFP_MAGED1_UN - GFP) |
| KPNA1 | P52294 | Importin subunit alpha-5 OS=Homo sapi | -2,453660941 | 3,33943E-08 | 7,476327012 | 3,9741E-06 | (GFP_MAGED1_HD - GFP)-(GFP_MAGED1_UN - GFP) |
| SLC25A12 | O75746 | Calcium-binding mitochondrial carrier pr | -2,621942127 | 3,34379E-08 | 7,475760486 | 3,9741E-06 | (GFP_MAGED1_HD - GFP)-(GFP_MAGED1_UN - GFP) |
| USP9X | Q93008\|Q93008-1 | Probable ubiquitin carboxyl-terminal hyd | -2,423823934 | 3,99437E-08 | 7,39855181 | 4,31573E-06 | (GFP_MAGED1_HD - GFP)-(GFP_MAGED1_UN - GFP) |
| TRIM33 | E7EN20\|Q9UPN9\|Q9UPN9-2 | E3 ubiquitin-protein ligase TRIM33 OS= | -2,7213474 | 4,24693E-08 | 7,371925188 | 4,38911E-06 | (GFP_MAGED1_HD - GFP)-(GFP_MAGED1_UN - GFP) |
| [illegible] | [illegible] | [illegible] | [illegible] | [illegible] | [illegible] | [illegible] | [illegible] |

<table>
<tr><td>MYO5B</td><td>Q9ULV0</td><td>Unconventional myosin-Vb OS=Homo s</td><td>-2,290669981</td><td>4,47364E-08</td><td>7,349339252</td><td>4,41918E-06</td><td>(GFP_MAGED1_HD - GFP)-(GFP_MAGED1_UN - GFP)</td></tr>
<tr><td>MYO1B</td><td>C9K0I9|E9PDF6|O43795|O4</td><td>Unconventional myosin-Ib (Fragment) O</td><td>-2,097722887</td><td>4,89725E-08</td><td>7,310048145</td><td>4,45835E-06</td><td>(GFP_MAGED1_HD - GFP)-(GFP_MAGED1_UN - GFP)</td></tr>
<tr><td>MYL6</td><td>B7Z6Z4|F8W1R7|G3V1V0|G</td><td>Myosin light polypeptide 6 OS=Homo sa</td><td>-3,896787152</td><td>5,06418E-08</td><td>7,295490858</td><td>4,45835E-06</td><td>(GFP_MAGED1_HD - GFP)-(GFP_MAGED1_UN - GFP)</td></tr>
<tr><td>TRAF2</td><td>Q12933|Q12933-2|Q12933-3</td><td>TNF receptor-associated factor 2 OS=Ho</td><td>-3,434920031</td><td>5,43984E-08</td><td>7,264414098</td><td>4,61803E-06</td><td>(GFP_MAGED1_HD - GFP)-(GFP_MAGED1_UN - GFP)</td></tr>
<tr><td>BIRC2</td><td>E9PMH5|Q13490|Q13490-2</td><td>Baculoviral IAP repeat-containing protei</td><td>-3,009509962</td><td>7,83394E-08</td><td>7,106019673</td><td>6,42113E-06</td><td>(GFP_MAGED1_HD - GFP)-(GFP_MAGED1_UN - GFP)</td></tr>
<tr><td>KLHL21</td><td>Q9UJP4|Q9UJP4-2</td><td>Kelch-like protein 21 OS=Homo sapiens</td><td>-2,109002303</td><td>8,42902E-08</td><td>7,074222844</td><td>6,67859E-06</td><td>(GFP_MAGED1_HD - GFP)-(GFP_MAGED1_UN - GFP)</td></tr>
<tr><td>PKP2</td><td>Q99959|Q99959-2</td><td>Plakophilin-2 OS=Homo sapiens GN=PK</td><td>-2,037411799</td><td>9,99472E-08</td><td>7,000229573</td><td>7,66369E-06</td><td>(GFP_MAGED1_HD - GFP)-(GFP_MAGED1_UN - GFP)</td></tr>
<tr><td>DCAF5</td><td>Q96JK2|Q96JK2-3</td><td>DDB1- and CUL4-associated factor 5 OS</td><td>-2,456911066</td><td>1,05454E-07</td><td>6,976938632</td><td>7,83322E-06</td><td>(GFP_MAGED1_HD - GFP)-(GFP_MAGED1_UN - GFP)</td></tr>
<tr><td>SMAP2</td><td>A0A087WV97|Q8WU79|Q8W</td><td>Stromal membrane-associated protein 2</td><td>-3,594630776</td><td>1,20756E-07</td><td>6,91808968</td><td>8,4423E-06</td><td>(GFP_MAGED1_HD - GFP)-(GFP_MAGED1_UN - GFP)</td></tr>
<tr><td>KLHL7</td><td>Q8IXQ5|Q8IXQ5-2|Q8IXQ5-5</td><td>Kelch-like protein 7 OS=Homo sapiens (</td><td>-2,502664559</td><td>1,27136E-07</td><td>6,895729927</td><td>8,63438E-06</td><td>(GFP_MAGED1_HD - GFP)-(GFP_MAGED1_UN - GFP)</td></tr>
<tr><td>DDX1</td><td>Q92499</td><td>ATP-dependent RNA helicase DDX1 OS</td><td>-1,905781087</td><td>1,45005E-07</td><td>6,838615862</td><td>9,57438E-06</td><td>(GFP_MAGED1_HD - GFP)-(GFP_MAGED1_UN - GFP)</td></tr>
<tr><td>HIPK3</td><td>Q9H422|Q9H422-2</td><td>Homeodomain-interacting protein kinase</td><td>-2,141386067</td><td>1,65129E-07</td><td>6,782176785</td><td>1,06084E-05</td><td>(GFP_MAGED1_HD - GFP)-(GFP_MAGED1_UN - GFP)</td></tr>
<tr><td>PUM1</td><td>H0YEH2|Q14671</td><td>Pumilio homolog 1 (Fragment) OS=Hom</td><td>-1,820366928</td><td>1,79363E-07</td><td>6,746267757</td><td>1,09954E-05</td><td>(GFP_MAGED1_HD - GFP)-(GFP_MAGED1_UN - GFP)</td></tr>
<tr><td>TRIM27</td><td>P14373|P14373-2</td><td>Zinc finger protein RFP OS=Homo sapie</td><td>-2,639531282</td><td>1,80404E-07</td><td>6,743754706</td><td>1,09954E-05</td><td>(GFP_MAGED1_HD - GFP)-(GFP_MAGED1_UN - GFP)</td></tr>
<tr><td>MED14</td><td>O60244</td><td>Mediator of RNA polymerase II transcript</td><td>-1,778071679</td><td>2,0057E-07</td><td>6,697733254</td><td>1,13513E-05</td><td>(GFP_MAGED1_HD - GFP)-(GFP_MAGED1_UN - GFP)</td></tr>
<tr><td>SLC25A13</td><td>Q9UJS0|Q9UJS0-2</td><td>Calcium-binding mitochondrial carrier pr</td><td>-1,850285812</td><td>2,60172E-07</td><td>6,584738688</td><td>1,43077E-05</td><td>(GFP_MAGED1_HD - GFP)-(GFP_MAGED1_UN - GFP)</td></tr>
<tr><td>TNIP2</td><td>Q8NFZ5</td><td>TNFAIP3-interacting protein 2 OS=Homo</td><td>-1,876481592</td><td>2,66801E-07</td><td>6,573812399</td><td>1,43077E-05</td><td>(GFP_MAGED1_HD - GFP)-(GFP_MAGED1_UN - GFP)</td></tr>
<tr><td>PUM2</td><td>Q8TB72-3|Q8TB72-4</td><td>Isoform 3 of Pumilio homolog 2 OS=Hom</td><td>-1,95453981</td><td>2,73767E-07</td><td>6,562619316</td><td>1,43077E-05</td><td>(GFP_MAGED1_HD - GFP)-(GFP_MAGED1_UN - GFP)</td></tr>
<tr><td>FAR1</td><td>Q8WVX9</td><td>Fatty acyl-CoA reductase 1 OS=Homo sa</td><td>-1,827900795</td><td>2,78924E-07</td><td>6,554513668</td><td>1,43077E-05</td><td>(GFP_MAGED1_HD - GFP)-(GFP_MAGED1_UN - GFP)</td></tr>
<tr><td>DCAF7</td><td>P61962</td><td>DDB1- and CUL4-associated factor 7 OS</td><td>-2,213531665</td><td>2,82903E-07</td><td>6,548362573</td><td>1,43077E-05</td><td>(GFP_MAGED1_HD - GFP)-(GFP_MAGED1_UN - GFP)</td></tr>
<tr><td>BTBD1</td><td>Q9H0C5|Q9H0C5-2</td><td>BTB/POZ domain-containing protein 1 O</td><td>-1,892678719</td><td>3,13809E-07</td><td>6,503334497</td><td>1,52229E-05</td><td>(GFP_MAGED1_HD - GFP)-(GFP_MAGED1_UN - GFP)</td></tr>
<tr><td>NYNRIN</td><td>Q9P2P1</td><td>Protein NYNRIN OS=Homo sapiens GN</td><td>-1,905888699</td><td>3,39166E-07</td><td>6,469587531</td><td>1,60679E-05</td><td>(GFP_MAGED1_HD - GFP)-(GFP_MAGED1_UN - GFP)</td></tr>
<tr><td>LRPAP1</td><td>P30533</td><td>Alpha-2-macroglobulin receptor-associa</td><td>-1,938583425</td><td>3,55748E-07</td><td>6,448857519</td><td>1,60679E-05</td><td>(GFP_MAGED1_HD - GFP)-(GFP_MAGED1_UN - GFP)</td></tr>
<tr><td>SON</td><td>P18583|P18583-10|P18583-</td><td>Protein SON OS=Homo sapiens GN=SO</td><td>-1,764239524</td><td>3,59389E-07</td><td>6,444435042</td><td>1,60679E-05</td><td>(GFP_MAGED1_HD - GFP)-(GFP_MAGED1_UN - GFP)</td></tr>
<tr><td>CRBN</td><td>J3QT87|Q96SW2|Q96SW2-2</td><td>Protein cereblon (Fragment) OS=Homo</td><td>-2,156725798</td><td>3,73005E-07</td><td>6,428285033</td><td>1,60679E-05</td><td>(GFP_MAGED1_HD - GFP)-(GFP_MAGED1_UN - GFP)</td></tr>
<tr><td>LRP4</td><td>O75096</td><td>Low-density lipoprotein receptor-related</td><td>-2,713523938</td><td>3,82627E-07</td><td>6,417224367</td><td>1,60679E-05</td><td>(GFP_MAGED1_HD - GFP)-(GFP_MAGED1_UN - GFP)</td></tr>
<tr><td>ANKRD40</td><td>Q6AI12</td><td>Ankyrin repeat domain-containing protei</td><td>-1,851280634</td><td>3,85304E-07</td><td>6,414196004</td><td>1,60679E-05</td><td>(GFP_MAGED1_HD - GFP)-(GFP_MAGED1_UN - GFP)</td></tr>
<tr><td>FHL2</td><td>A0A0A0MSG2|J3KNW4|Q14</td><td>Four and a half LIM domains protein 2 O</td><td>-1,932782097</td><td>3,93574E-07</td><td>6,404973995</td><td>1,61135E-05</td><td>(GFP_MAGED1_HD - GFP)-(GFP_MAGED1_UN - GFP)</td></tr>
<tr><td>MYO5A</td><td>A0A087WY00|F8W6H6|Q9Y</td><td>Unconventional myosin-Va OS=Homo s</td><td>-1,649967251</td><td>4,023E-07</td><td>6,395450209</td><td>1,61135E-05</td><td>(GFP_MAGED1_HD - GFP)-(GFP_MAGED1_UN - GFP)</td></tr>
<tr><td>CUL3</td><td>Q13618</td><td>Cullin-3 OS=Homo sapiens GN=CUL3 F</td><td>-1,559052083</td><td>4,51612E-07</td><td>6,345234963</td><td>1,75249E-05</td><td>(GFP_MAGED1_HD - GFP)-(GFP_MAGED1_UN - GFP)</td></tr>
<tr><td>HNRNPUL1</td><td>A0A0A0MRA5|B7Z4B8|Q9BU</td><td>Heterogeneous nuclear ribonucleoprote</td><td>-1,857014423</td><td>4,57108E-07</td><td>6,339981628</td><td>1,75249E-05</td><td>(GFP_MAGED1_HD - GFP)-(GFP_MAGED1_UN - GFP)</td></tr>
<tr><td>SPOP</td><td>O43791</td><td>Speckle-type POZ protein OS=Homo sap</td><td>-2,601542041</td><td>4,74849E-07</td><td>6,323444643</td><td>1,79161E-05</td><td>(GFP_MAGED1_HD - GFP)-(GFP_MAGED1_UN - GFP)</td></tr>
<tr><td>MAPK1</td><td>P28482</td><td>Mitogen-activated protein kinase 1 OS=I</td><td>-1,817160703</td><td>5,07589E-07</td><td>6,294487668</td><td>1,88522E-05</td><td>(GFP_MAGED1_HD - GFP)-(GFP_MAGED1_UN - GFP)</td></tr>
<tr><td>C14orf166</td><td>Q9Y224</td><td>UPF0568 protein C14orf166 OS=Homo</td><td>-1,725533987</td><td>5,95994E-07</td><td>6,224757795</td><td>2,17951E-05</td><td>(GFP_MAGED1_HD - GFP)-(GFP_MAGED1_UN - GFP)</td></tr>
<tr><td>SEC16A</td><td>J3KNL6|O15027|O15027-5</td><td>Protein transport protein Sec16A OS=Ho</td><td>-1,76952013</td><td>6,44655E-07</td><td>6,19067263</td><td>2,32173E-05</td><td>(GFP_MAGED1_HD - GFP)-(GFP_MAGED1_UN - GFP)</td></tr>
<tr><td>RTCB</td><td>Q9Y3I0</td><td>tRNA-splicing ligase RtcB homolog OS=</td><td>-1,628526225</td><td>6,98372E-07</td><td>6,155913305</td><td>2,46111E-05</td><td>(GFP_MAGED1_HD - GFP)-(GFP_MAGED1_UN - GFP)</td></tr>
<tr><td>MYO9A</td><td>B2RTY4|B2RTY4-4|H3BRD5</td><td>Unconventional myosin-IXa OS=Homo s</td><td>-1,85108965</td><td>7,04062E-07</td><td>6,152389398</td><td>2,46111E-05</td><td>(GFP_MAGED1_HD - GFP)-(GFP_MAGED1_UN - GFP)</td></tr>
<tr><td>MYO1C</td><td>F5H6E2|O00159|O00159-2|(</td><td>Unconventional myosin-Ic OS=Homo sa</td><td>-1,484489574</td><td>7,28533E-07</td><td>6,137550877</td><td>2,50974E-05</td><td>(GFP_MAGED1_HD - GFP)-(GFP_MAGED1_UN - GFP)</td></tr>
<tr><td>POLR3C</td><td>Q9BUI4</td><td>DNA-directed RNA polymerase III subun</td><td>-1,672238104</td><td>7,488E-07</td><td>6,12563445</td><td>2,54271E-05</td><td>(GFP_MAGED1_HD - GFP)-(GFP_MAGED1_UN - GFP)</td></tr>
<tr><td>DNAJC11</td><td>Q9NVH1|Q9NVH1-3</td><td>DnaJ homolog subfamily C member 11 (</td><td>-1,506791709</td><td>8,44353E-07</td><td>6,073476194</td><td>2,8268E-05</td><td>(GFP_MAGED1_HD - GFP)-(GFP_MAGED1_UN - GFP)</td></tr>
<tr><td>TRIM24</td><td>A0A087X1V3|O15164|O1516</td><td>Transcription intermediary factor 1-alpha</td><td>-2,87420384</td><td>8,7955E-07</td><td>6,055739274</td><td>2,90374E-05</td><td>(GFP_MAGED1_HD - GFP)-(GFP_MAGED1_UN - GFP)</td></tr>
<tr><td>KANK2</td><td>Q63ZY3|Q63ZY3-2|Q63ZY3-</td><td>KN motif and ankyrin repeat domain-con</td><td>-1,833628571</td><td>8,96095E-07</td><td>6,047645736</td><td>2,91783E-05</td><td>(GFP_MAGED1_HD - GFP)-(GFP_MAGED1_UN - GFP)</td></tr>
<tr><td>TRIM11</td><td>Q96F44|Q96F44-3|R4GMV1</td><td>E3 ubiquitin-protein ligase TRIM11 OS=</td><td>-1,730420417</td><td>9,32332E-07</td><td>6,030429221</td><td>2,97134E-05</td><td>(GFP_MAGED1_HD - GFP)-(GFP_MAGED1_UN - GFP)</td></tr>
<tr><td>STK4</td><td>Q13043</td><td>Serine/threonine-protein kinase 4 OS=H</td><td>-1,68241388</td><td>9,3753E-07</td><td>6,028014843</td><td>2,97134E-05</td><td>(GFP_MAGED1_HD - GFP)-(GFP_MAGED1_UN - GFP)</td></tr>
<tr><td>SSBP1</td><td>A0A0G2JLD8|E7EUY5|Q048</td><td>Single-stranded DNA-binding protein, m</td><td>-1,867220493</td><td>9,86987E-07</td><td>6,005688387</td><td>3,08693E-05</td><td>(GFP_MAGED1_HD - GFP)-(GFP_MAGED1_UN - GFP)</td></tr>
<tr><td>KLHL15</td><td>Q96M94</td><td>Kelch-like protein 15 OS=Homo sapiens</td><td>-1,60293816</td><td>1,08295E-06</td><td>5,965391989</td><td>3,34308E-05</td><td>(GFP_MAGED1_HD - GFP)-(GFP_MAGED1_UN - GFP)</td></tr>
<tr><td>CNOT1</td><td>A5YKK6</td><td>CCR4-NOT transcription complex subun</td><td>-1,404621595</td><td>1,23919E-06</td><td>5,906861102</td><td>3,72856E-05</td><td>(GFP_MAGED1_HD - GFP)-(GFP_MAGED1_UN - GFP)</td></tr>
<tr><td>IPP</td><td>Q9Y573|Q9Y573-2</td><td>Actin-binding protein IPP OS=Homo sap</td><td>-2,774121927</td><td>1,33856E-06</td><td>5,873362923</td><td>3,97719E-05</td><td>(GFP_MAGED1_HD - GFP)-(GFP_MAGED1_UN - GFP)</td></tr>
<tr><td>LRP1</td><td>Q07954</td><td>Prolow-density lipoprotein receptor-relat</td><td>-1,430379077</td><td>1,42878E-06</td><td>5,845034258</td><td>4,19286E-05</td><td>(GFP_MAGED1_HD - GFP)-(GFP_MAGED1_UN - GFP)</td></tr>
<tr><td>LETMD1</td><td>F8VP71|F8VVQ3|F8VZA2|Q6</td><td>LETM1 domain-containing protein 1 OS=</td><td>-1,869669865</td><td>1,44729E-06</td><td>5,839443269</td><td>4,19539E-05</td><td>(GFP_MAGED1_HD - GFP)-(GFP_MAGED1_UN - GFP)</td></tr>
<tr><td>TNRC6B</td><td>Q9UPQ9|Q9UPQ9-1</td><td>Trinucleotide repeat-containing gene 6B</td><td>-2,261872469</td><td>1,48928E-06</td><td>5,827023444</td><td>4,26508E-05</td><td>(GFP_MAGED1_HD - GFP)-(GFP_MAGED1_UN - GFP)</td></tr>
<tr><td>MBNL3</td><td>B1AKI6|Q9NUK0|Q9NUK0-2</td><td>Muscleblind-like protein 3 OS=Homo sa</td><td>-1,540362908</td><td>1,61352E-06</td><td>5,792226336</td><td>4,51215E-05</td><td>(GFP_MAGED1_HD - GFP)-(GFP_MAGED1_UN - GFP)</td></tr>
<tr><td>RBM14</td><td>Q96PK6</td><td>RNA-binding protein 14 OS=Homo sapie</td><td>-1,756880727</td><td>1,6395E-06</td><td>5,785288024</td><td>4,53151E-05</td><td>(GFP_MAGED1_HD - GFP)-(GFP_MAGED1_UN - GFP)</td></tr>
<tr><td>SLC25A16</td><td>P16260</td><td>Graves disease carrier protein OS=Hom</td><td>-1,950382536</td><td>1,66447E-06</td><td>5,778723365</td><td>4,53934E-05</td><td>(GFP_MAGED1_HD - GFP)-(GFP_MAGED1_UN - GFP)</td></tr>
<tr><td>CYP2J2</td><td>P51589</td><td>Cytochrome P450 2J2 OS=Homo sapien</td><td>-2,84804528</td><td>1,68053E-06</td><td>5,774553628</td><td>4,53934E-05</td><td>(GFP_MAGED1_HD - GFP)-(GFP_MAGED1_UN - GFP)</td></tr>
<tr><td>CPVL</td><td>Q9H3G5</td><td>Probable serine carboxypeptidase CPVI</td><td>-1,478383068</td><td>1,92315E-06</td><td>5,715987111</td><td>4,9154E-05</td><td>(GFP_MAGED1_HD - GFP)-(GFP_MAGED1_UN - GFP)</td></tr>
<tr><td>AGPS</td><td>O00116</td><td>Alkyldihydroxyacetonephosphate syntha</td><td>-1,581085344</td><td>1,95669E-06</td><td>5,708477734</td><td>4,94793E-05</td><td>(GFP_MAGED1_HD - GFP)-(GFP_MAGED1_UN - GFP)</td></tr>
<tr><td>KPNA3</td><td>O00505</td><td>Importin subunit alpha-4 OS=Homo sapi</td><td>-1,804403758</td><td>2,01487E-06</td><td>5,695752722</td><td>5,04142E-05</td><td>(GFP_MAGED1_HD - GFP)-(GFP_MAGED1_UN - GFP)</td></tr>
<tr><td>SORT1</td><td>Q99523|Q99523-2</td><td>Sortilin OS=Homo sapiens GN=SORT1 I</td><td>-1,383408818</td><td>2,39795E-06</td><td>5,620160591</td><td>5,74357E-05</td><td>(GFP_MAGED1_HD - GFP)-(GFP_MAGED1_UN - GFP)</td></tr>
<tr><td>DHCR24|Nbla03</td><td>Q15392|Q15392-2|Q3LIE7</td><td>Delta(24)-sterol reductase OS=Homo sa</td><td>-1,725729192</td><td>2,50819E-06</td><td>5,600639038</td><td>5,88907E-05</td><td>(GFP_MAGED1_HD - GFP)-(GFP_MAGED1_UN - GFP)</td></tr>
<tr><td>LRP6</td><td>F5H7J9|O75581</td><td>Low-density lipoprotein receptor-related</td><td>-1,740013465</td><td>2,52707E-06</td><td>5,597382542</td><td>5,88907E-05</td><td>(GFP_MAGED1_HD - GFP)-(GFP_MAGED1_UN - GFP)</td></tr>
<tr><td>ITCH</td><td>Q96J02|Q96J02-2|Q96J02-3</td><td>E3 ubiquitin-protein ligase Itchy homolog</td><td>-2,279280329</td><td>2,67132E-06</td><td>5,573273374</td><td>6,16479E-05</td><td>(GFP_MAGED1_HD - GFP)-(GFP_MAGED1_UN - GFP)</td></tr>
<tr><td>NDUFA12</td><td>F8VRD8|H0YID5|Q9UI09|Q9</td><td>NADH dehydrogenase [ubiquinone] 1 al</td><td>-1,278689831</td><td>3,19173E-06</td><td>5,495974429</td><td>7,15729E-05</td><td>(GFP_MAGED1_HD - GFP)-(GFP_MAGED1_UN - GFP)</td></tr>
<tr><td>LMNB1</td><td>P20700</td><td>Lamin-B1 OS=Homo sapiens GN=LMNE</td><td>-1,729155939</td><td>3,50354E-06</td><td>5,455492943</td><td>7,7831E-05</td><td>(GFP_MAGED1_HD - GFP)-(GFP_MAGED1_UN - GFP)</td></tr>
<tr><td>AP2A1</td><td>O95782|O95782-2</td><td>AP-2 complex subunit alpha-1 OS=Hom</td><td>-1,365127532</td><td>4,29994E-06</td><td>5,366537283</td><td>9,26172E-05</td><td>(GFP_MAGED1_HD - GFP)-(GFP_MAGED1_UN - GFP)</td></tr>
<tr><td>POMGNT2</td><td>Q8NAT1</td><td>Protein O-linked-mannose beta-1,4-N-ac</td><td>-1,754118245</td><td>4,47528E-06</td><td>5,349180019</td><td>9,39231E-05</td><td>(GFP_MAGED1_HD - GFP)-(GFP_MAGED1_UN - GFP)</td></tr>
<tr><td>RNMT</td><td>O43148|O43148-2</td><td>mRNA cap guanine-N7 methyltransferas</td><td>-1,594465903</td><td>4,49606E-06</td><td>5,347167495</td><td>9,39231E-05</td><td>(GFP_MAGED1_HD - GFP)-(GFP_MAGED1_UN - GFP)</td></tr>
<tr><td>FAM120A</td><td>Q9NZB2|Q9NZB2-4|Q9NZB2</td><td>Constitutive coactivator of PPAR-gamma</td><td>-1,308869153</td><td>4,53973E-06</td><td>5,342970219</td><td>9,39231E-05</td><td>(GFP_MAGED1_HD - GFP)-(GFP_MAGED1_UN - GFP)</td></tr>
<tr><td>CETN3</td><td>E5RJF8|E5RK82|O15182</td><td>Centrin-3 OS=Homo sapiens GN=CETN</td><td>-1,195002282</td><td>4,54403E-06</td><td>5,342559118</td><td>9,39231E-05</td><td>(GFP_MAGED1_HD - GFP)-(GFP_MAGED1_UN - GFP)</td></tr>
<tr><td>CDC23</td><td>Q9UJX2</td><td>Cell division cycle protein 23 homolog C</td><td>-1,216308696</td><td>4,76339E-06</td><td>5,322083442</td><td>9,76085E-05</td><td>(GFP_MAGED1_HD - GFP)-(GFP_MAGED1_UN - GFP)</td></tr>
<tr><td>NEFL</td><td>P07196</td><td>Neurofilament light polypeptide OS=Hor</td><td>-1,177826491</td><td>5,02587E-06</td><td>5,298789096</td><td>0,000102107</td><td>(GFP_MAGED1_HD - GFP)-(GFP_MAGED1_UN - GFP)</td></tr>
<tr><td>EMD</td><td>P50402</td><td>Emerin OS=Homo sapiens GN=EMD PE</td><td>-1,236146368</td><td>5,49602E-06</td><td>5,259951738</td><td>0,000110712</td><td>(GFP_MAGED1_HD - GFP)-(GFP_MAGED1_UN - GFP)</td></tr>
<tr><td>MAP1B</td><td>P46821</td><td>Microtubule-associated protein 1B OS=I</td><td>-1,446715563</td><td>5,74502E-06</td><td>5,240708479</td><td>0,000114707</td><td>(GFP_MAGED1_HD - GFP)-(GFP_MAGED1_UN - GFP)</td></tr>
<tr><td>CNOT2</td><td>F8VV52|Q9NZN8</td><td>CCR4-NOT transcription complex subun</td><td>-1,51193301</td><td>6,31555E-06</td><td>5,199588851</td><td>0,00012305</td><td>(GFP_MAGED1_HD - GFP)-(GFP_MAGED1_UN - GFP)</td></tr>
<tr><td>TRNAU1AP</td><td>Q9NX07|Q9NX07-2</td><td>tRNA selenocysteine 1-associated prote</td><td>-1,83573335</td><td>6,46138E-06</td><td>5,189674509</td><td>0,000123911</td><td>(GFP_MAGED1_HD - GFP)-(GFP_MAGED1_UN - GFP)</td></tr>
<tr><td>HNRNPUL2</td><td>Q1KMD3</td><td>Heterogeneous nuclear ribonucleoprote</td><td>-1,452293298</td><td>6,46401E-06</td><td>5,189497979</td><td>0,000123911</td><td>(GFP_MAGED1_HD - GFP)-(GFP_MAGED1_UN - GFP)</td></tr>
<tr><td>MCMBP</td><td>Q9BTE3|Q9BTE3-2</td><td>Mini-chromosome maintenance complex</td><td>-1,307541014</td><td>7,20642E-06</td><td>5,142280536</td><td>0,000136661</td><td>(GFP_MAGED1_HD - GFP)-(GFP_MAGED1_UN - GFP)</td></tr>
<tr><td>FBRSL1</td><td>Q9HCM7</td><td>Fibrosin-1-like protein OS=Homo sapien</td><td>-2,74705261</td><td>7,4426E-06</td><td>5,128275591</td><td>0,0001393</td><td>(GFP_MAGED1_HD - GFP)-(GFP_MAGED1_UN - GFP)</td></tr>
<tr><td>ATXN2L</td><td>Q8WWM7|Q8WWM7-2|Q8W</td><td>Ataxin-2-like protein OS=Homo sapiens</td><td>-1,239875888</td><td>7,69072E-06</td><td>5,114032957</td><td>0,000142819</td><td>(GFP_MAGED1_HD - GFP)-(GFP_MAGED1_UN - GFP)</td></tr>
<tr><td>IFFO1</td><td>A0A087WZ16|Q0D2I5|Q0D2</td><td>Intermediate filament family orphan 1 OS</td><td>-1,782833885</td><td>7,91509E-06</td><td>5,101543988</td><td>0,000145846</td><td>(GFP_MAGED1_HD - GFP)-(GFP_MAGED1_UN - GFP)</td></tr>
<tr><td>STAT3</td><td>P40763|P40763-2|P40763-3</td><td>Signal transducer and activator of transc</td><td>-1,191168177</td><td>8,5142E-06</td><td>5,069856374</td><td>0,000155679</td><td>(GFP_MAGED1_HD - GFP)-(GFP_MAGED1_UN - GFP)</td></tr>
<tr><td>MYO1D</td><td>O94832</td><td>Unconventional myosin-Id OS=Homo sa</td><td>-1,259550174</td><td>8,8063E-06</td><td>5,055206534</td><td>0,000157892</td><td>(GFP_MAGED1_HD - GFP)-(GFP_MAGED1_UN - GFP)</td></tr>
<tr><td>FAM83H</td><td>J3KPS2|Q6ZRV2</td><td>Protein FAM83H (Fragment) OS=Homo</td><td>-1,235921297</td><td>8,93996E-06</td><td>5,048664279</td><td>0,000158584</td><td>(GFP_MAGED1_HD - GFP)-(GFP_MAGED1_UN - GFP)</td></tr>
<tr><td>GNS</td><td>F5H4C6|F6S8M0|H7C3P4|P</td><td>N-acetylglucosamine-6-sulfatase (Fragm</td><td>-1,116738005</td><td>9,05208E-06</td><td>5,043251514</td><td>0,000158713</td><td>(GFP_MAGED1_HD - GFP)-(GFP_MAGED1_UN - GFP)</td></tr>
<tr><td>PIGA</td><td>P37287|P37287-3</td><td>Phosphatidylinositol N-acetylglucosamir</td><td>-1,398797999</td><td>9,14754E-06</td><td>5,038695811</td><td>0,000158713</td><td>(GFP_MAGED1_HD - GFP)-(GFP_MAGED1_UN - GFP)</td></tr>
<tr><td>SLC25A11</td><td>I3L1P8|Q02978</td><td>Mitochondrial 2-oxoglutarate/malate car</td><td>-1,40063787</td><td>9,21567E-06</td><td>5,03547304</td><td>0,000158737</td><td>(GFP_MAGED1_HD - GFP)-(GFP_MAGED1_UN - GFP)</td></tr>
<tr><td>POM121C</td><td>A0A075B7F8|A8CG34</td><td>Nuclear envelope pore membrane prote</td><td>-1,477747211</td><td>9,32725E-06</td><td>5,030246217</td><td>0,000159503</td><td>(GFP_MAGED1_HD - GFP)-(GFP_MAGED1_UN - GFP)</td></tr>
<tr><td>SORL1</td><td>Q92673</td><td>Sortilin-related receptor OS=Homo sapie</td><td>-1,188205444</td><td>9,45999E-06</td><td>5,024109358</td><td>0,000160613</td><td>(GFP_MAGED1_HD - GFP)-(GFP_MAGED1_UN - GFP)</td></tr>
<tr><td>XRN2</td><td>Q9H0D6</td><td>5'-3' exoribonuclease 2 OS=Homo sapie</td><td>-1,163966455</td><td>9,52731E-06</td><td>5,021029559</td><td>0,000160613</td><td>(GFP_MAGED1_HD - GFP)-(GFP_MAGED1_UN - GFP)</td></tr>
<tr><td>IFT172</td><td>Q9UG01</td><td>Intraflagellar transport protein 172 homo</td><td>-1,171532231</td><td>9,79674E-06</td><td>5,00891822</td><td>0,000161714</td><td>(GFP_MAGED1_HD - GFP)-(GFP_MAGED1_UN - GFP)</td></tr>
<tr><td>AGO1</td><td>Q5TA58|Q9UL18</td><td>Protein argonaute-1 OS=Homo sapiens</td><td>-1,474591208</td><td>1,00529E-05</td><td>4,997708184</td><td>0,000164798</td><td>(GFP_MAGED1_HD - GFP)-(GFP_MAGED1_UN - GFP)</td></tr>
<tr><td>OFD1</td><td>O75665</td><td>Oral-facial-digital syndrome 1 protein OS</td><td>-1,243993006</td><td>1,02828E-05</td><td>4,987890581</td><td>0,000167412</td><td>(GFP_MAGED1_HD - GFP)-(GFP_MAGED1_UN - GFP)</td></tr>
<tr><td>ERC1</td><td>G8JLD3|Q8IUD2|Q8IUD2-2|(</td><td>ELKS/Rab6-interacting/CAST family mer</td><td>-1,457751381</td><td>1,05597E-05</td><td>4,976349509</td><td>0,000168459</td><td>(GFP_MAGED1_HD - GFP)-(GFP_MAGED1_UN - GFP)</td></tr>
<tr><td>CSNK2A1</td><td>E7EU96|P68400|Q5U5J2</td><td>Casein kinase II subunit alpha OS=Hom</td><td>-1,051921011</td><td>1,11628E-05</td><td>4,952225692</td><td>0,000175449</td><td>(GFP_MAGED1_HD - GFP)-(GFP_MAGED1_UN - GFP)</td></tr>
<tr><td>CCDC85C</td><td>A6NKD9</td><td>Coiled-coil domain-containing protein 8</td><td>-2,237066093</td><td>1,12894E-05</td><td>4,947328394</td><td>0,000175449</td><td>(GFP_MAGED1_HD - GFP)-(GFP_MAGED1_UN - GFP)</td></tr>
<tr><td>LMNB2</td><td>J9JID7|Q03252</td><td>Lamin B2, isoform CRA_a OS=Homo sa</td><td>-1,227244563</td><td>1,20453E-05</td><td>4,919181857</td><td>0,000184721</td><td>(GFP_MAGED1_HD - GFP)-(GFP_MAGED1_UN - GFP)</td></tr>
<tr><td>AHCY</td><td>P23526</td><td>Adenosylhomocysteinase OS=Homo sa</td><td>-1,043557936</td><td>1,28736E-05</td><td>4,890299878</td><td>0,000194908</td><td>(GFP_MAGED1_HD - GFP)-(GFP_MAGED1_UN - GFP)</td></tr>
<tr><td>GLUL</td><td>P15104</td><td>Glutamine synthetase OS=Homo sapien</td><td>-1,499267788</td><td>1,33407E-05</td><td>4,874822546</td><td>0,000199439</td><td>(GFP_MAGED1_HD - GFP)-(GFP_MAGED1_UN - GFP)</td></tr>
<tr><td>PABPC4</td><td>Q13310-2|Q13310-3</td><td>Isoform 2 of Polyadenylate-binding prote</td><td>-1,128987508</td><td>1,52085E-05</td><td>4,81791371</td><td>0,000224538</td><td>(GFP_MAGED1_HD - GFP)-(GFP_MAGED1_UN - GFP)</td></tr>
<tr><td>ANP32A</td><td>H0YN26|P39687</td><td>Acidic leucine-rich nuclear phosphoprot</td><td>-2,327720336</td><td>1,55738E-05</td><td>4,807604219</td><td>0,000228512</td><td>(GFP_MAGED1_HD - GFP)-(GFP_MAGED1_UN - GFP)</td></tr>
<tr><td>PDCD6</td><td>O75340|O75340-2</td><td>Programmed cell death protein 6 OS=Ho</td><td>-2,131908124</td><td>1,66637E-05</td><td>4,778229693</td><td>0,000243003</td><td>(GFP_MAGED1_HD - GFP)-(GFP_MAGED1_UN - GFP)</td></tr>
<tr><td>RGPD5</td><td>Q99666</td><td>RANBP2-like and GRIP domain-contain</td><td>-1,22498604</td><td>1,68568E-05</td><td>4,773225309</td><td>0,000244321</td><td>(GFP_MAGED1_HD - GFP)-(GFP_MAGED1_UN - GFP)</td></tr>
<tr><td>KPNA2</td><td>P52292</td><td>Importin subunit alpha-1 OS=Homo sapi</td><td>-1,054877848</td><td>1,85983E-05</td><td>4,730526378</td><td>0,000263144</td><td>(GFP_MAGED1_HD - GFP)-(GFP_MAGED1_UN - GFP)</td></tr>
<tr><td>ANP32B</td><td>Q92688|Q92688-2</td><td>Acidic leucine-rich nuclear phosphoprot</td><td>-3,093888329</td><td>1,97505E-05</td><td>4,704421596</td><td>0,000276796</td><td>(GFP_MAGED1_HD - GFP)-(GFP_MAGED1_UN - GFP)</td></tr>
<tr><td>DDA1</td><td>Q9BW61</td><td>DET1- and DDB1-associated protein 1 C</td><td>-1,959437246</td><td>2,10507E-05</td><td>4,676733801</td><td>0,000290916</td><td>(GFP_MAGED1_HD - GFP)-(GFP_MAGED1_UN - GFP)</td></tr>
<tr><td>AP1B1</td><td>Q10567|Q10567-2|Q10567-3</td><td>AP-1 complex subunit beta-1 OS=Homo</td><td>-1,715659364</td><td>2,16299E-05</td><td>4,664946389</td><td>0,000295484</td><td>(GFP_MAGED1_HD - GFP)-(GFP_MAGED1_UN - GFP)</td></tr>
<tr><td>WWP1</td><td>Q9H0M0|Q9H0M0-3</td><td>NEDD4-like E3 ubiquitin-protein ligase V</td><td>-1,125053618</td><td>2,20579E-05</td><td>4,65643612</td><td>0,000299609</td><td>(GFP_MAGED1_HD - GFP)-(GFP_MAGED1_UN - GFP)</td></tr>
<tr><td>ASPM</td><td>Q8IZT6</td><td>Abnormal spindle-like microcephaly-ass</td><td>-1,329289851</td><td>2,26946E-05</td><td>4,644077183</td><td>0,000305969</td><td>(GFP_MAGED1_HD - GFP)-(GFP_MAGED1_UN - GFP)</td></tr>
<tr><td>RNGTT</td><td>O60942|O60942-4</td><td>mRNA-capping enzyme OS=Homo sapi</td><td>-1,143564407</td><td>2,27836E-05</td><td>4,64237801</td><td>0,000305969</td><td>(GFP_MAGED1_HD - GFP)-(GFP_MAGED1_UN - GFP)</td></tr>
<tr><td>VPRBP</td><td>Q9Y4B6|Q9Y4B6-2</td><td>Protein VPRBP OS=Homo sapiens GN=</td><td>-1,076284854</td><td>2,2955E-05</td><td>4,639122315</td><td>0,00030654</td><td>(GFP_MAGED1_HD - GFP)-(GFP_MAGED1_UN - GFP)</td></tr>
<tr><td>ENPP4</td><td>Q9Y6X5</td><td>Bis(5'-adenosyl)-triphosphatase ENPP4</td><td>-1,034183116</td><td>2,32479E-05</td><td>4,633615496</td><td>0,000308717</td><td>(GFP_MAGED1_HD - GFP)-(GFP_MAGED1_UN - GFP)</td></tr>
</table>

<table>
<tr><td>ANKHD1</td><td>Q8IWZ3|Q8IWZ3-4|Q8IWZ3-6</td><td>Ankyrin repeat and KH domain-containir</td><td>-1,617348887</td><td>2,50787E-05</td><td>4,600694802</td><td>0,000325749</td><td>(GFP_MAGED1_HD - GFP)-(GFP_MAGED1_UN - GFP)</td></tr>
<tr><td>SLC25A33</td><td>Q9BSK2</td><td>Solute carrier family 25 member 33 OS=</td><td>-1,051055569</td><td>2,53224E-05</td><td>4,596494529</td><td>0,000327127</td><td>(GFP_MAGED1_HD - GFP)-(GFP_MAGED1_UN - GFP)</td></tr>
<tr><td>CSNK1A1</td><td>P48729|P48729-2</td><td>Casein kinase I isoform alpha OS=Homo</td><td>-1,148747068</td><td>2,9373E-05</td><td>4,532051781</td><td>0,000375374</td><td>(GFP_MAGED1_HD - GFP)-(GFP_MAGED1_UN - GFP)</td></tr>
<tr><td>CETN2</td><td>P41208</td><td>Centrin-2 OS=Homo sapiens GN=CETN</td><td>-1,343966116</td><td>2,96011E-05</td><td>4,528692274</td><td>0,000376266</td><td>(GFP_MAGED1_HD - GFP)-(GFP_MAGED1_UN - GFP)</td></tr>
<tr><td>TOMM20</td><td>Q15388</td><td>Mitochondrial import receptor subunit TO</td><td>-1,171654002</td><td>3,10843E-05</td><td>4,507458984</td><td>0,000390938</td><td>(GFP_MAGED1_HD - GFP)-(GFP_MAGED1_UN - GFP)</td></tr>
<tr><td>LMNA</td><td>P02545</td><td>Prelamin-A/C OS=Homo sapiens GN=LM</td><td>-1,226247706</td><td>3,3913E-05</td><td>4,46963348</td><td>0,00041985</td><td>(GFP_MAGED1_HD - GFP)-(GFP_MAGED1_UN - GFP)</td></tr>
<tr><td>TMEM33</td><td>D6RAA6|P57088</td><td>Transmembrane protein 33 (Fragment) (</td><td>-1,269093389</td><td>3,58637E-05</td><td>4,445344355</td><td>0,000440664</td><td>(GFP_MAGED1_HD - GFP)-(GFP_MAGED1_UN - GFP)</td></tr>
<tr><td>B4DLN1</td><td>B4DLN1</td><td>Uncharacterized protein OS=Homo sapi</td><td>-1,039259018</td><td>3,5965E-05</td><td>4,444119732</td><td>0,000440664</td><td>(GFP_MAGED1_HD - GFP)-(GFP_MAGED1_UN - GFP)</td></tr>
<tr><td>ALMS1</td><td>A0A087WTU9|A0A087WZY3</td><td>Alstrom syndrome protein 1 OS=Homo s</td><td>-1,433248775</td><td>3,62544E-05</td><td>4,440639395</td><td>0,000441932</td><td>(GFP_MAGED1_HD - GFP)-(GFP_MAGED1_UN - GFP)</td></tr>
<tr><td>HNRNPC</td><td>B2R5W2|B4DY08|G3V2Q1|G</td><td>Heterogeneous nuclear ribonucleoprote</td><td>-1,016229044</td><td>4,09979E-05</td><td>4,387238343</td><td>0,000482436</td><td>(GFP_MAGED1_HD - GFP)-(GFP_MAGED1_UN - GFP)</td></tr>
<tr><td>ANP32E</td><td>E9PPH5|Q9BTT0</td><td>Acidic leucine-rich nuclear phosphoprot</td><td>-2,644473173</td><td>4,40022E-05</td><td>4,356525972</td><td>0,000512711</td><td>(GFP_MAGED1_HD - GFP)-(GFP_MAGED1_UN - GFP)</td></tr>
<tr><td>TIMM23|TIMM23</td><td>O14925|Q5SRD1</td><td>Mitochondrial import inner membrane tra</td><td>-1,465401165</td><td>4,65861E-05</td><td>4,331743476</td><td>0,000532381</td><td>(GFP_MAGED1_HD - GFP)-(GFP_MAGED1_UN - GFP)</td></tr>
<tr><td>UBAP2L</td><td>F8W726|Q14157|Q14157-1|Q</td><td>Ubiquitin-associated protein 2-like OS=H</td><td>-1,784048372</td><td>4,76722E-05</td><td>4,321734876</td><td>0,000542186</td><td>(GFP_MAGED1_HD - GFP)-(GFP_MAGED1_UN - GFP)</td></tr>
<tr><td>EIF4G1</td><td>E7EUU4|Q04637|Q04637-3|</td><td>Eukaryotic translation initiation factor 4 g</td><td>-1,214609355</td><td>5,16278E-05</td><td>4,287116412</td><td>0,000576147</td><td>(GFP_MAGED1_HD - GFP)-(GFP_MAGED1_UN - GFP)</td></tr>
<tr><td>FLII</td><td>Q13045|Q13045-2|Q13045-3</td><td>Protein flightless-1 homolog OS=Homo s</td><td>-1,463082029</td><td>5,59313E-05</td><td>4,252345187</td><td>0,000598818</td><td>(GFP_MAGED1_HD - GFP)-(GFP_MAGED1_UN - GFP)</td></tr>
<tr><td>SAP18</td><td>O00422|X6RAL5</td><td>Histone deacetylase complex subunit SA</td><td>-1,013435038</td><td>5,63907E-05</td><td>4,248792639</td><td>0,000598818</td><td>(GFP_MAGED1_HD - GFP)-(GFP_MAGED1_UN - GFP)</td></tr>
<tr><td>MTA3</td><td>E7EQY4|E7EV10|Q9BTC8|Q</td><td>Metastasis-associated protein MTA3 OS</td><td>-1,129032703</td><td>5,64304E-05</td><td>4,248486523</td><td>0,000598818</td><td>(GFP_MAGED1_HD - GFP)-(GFP_MAGED1_UN - GFP)</td></tr>
<tr><td>GSTM3</td><td>A0A0A0MTN3|P21266</td><td>Glutathione S-transferase Mu 3 OS=Hon</td><td>-1,168932366</td><td>6,21624E-05</td><td>4,206472393</td><td>0,00064524</td><td>(GFP_MAGED1_HD - GFP)-(GFP_MAGED1_UN - GFP)</td></tr>
<tr><td>PDCD6IP</td><td>Q8WUM4</td><td>Programmed cell death 6-interacting pro</td><td>-1,162325505</td><td>6,5794E-05</td><td>4,181813815</td><td>0,000677023</td><td>(GFP_MAGED1_HD - GFP)-(GFP_MAGED1_UN - GFP)</td></tr>
<tr><td>DDB1</td><td>Q16531</td><td>DNA damage-binding protein 1 OS=Hor</td><td>-1,06904757</td><td>7,36981E-05</td><td>4,132543785</td><td>0,00074229</td><td>(GFP_MAGED1_HD - GFP)-(GFP_MAGED1_UN - GFP)</td></tr>
<tr><td>MED12</td><td>Q93074|Q93074-2|Q93074-3</td><td>Mediator of RNA polymerase II transcrip</td><td>-1,178917951</td><td>7,50009E-05</td><td>4,124933549</td><td>0,000749064</td><td>(GFP_MAGED1_HD - GFP)-(GFP_MAGED1_UN - GFP)</td></tr>
<tr><td>MSI2</td><td>B4DHE8|J3KTC1|Q96DH6|Q</td><td>RNA-binding protein Musashi homolog 2</td><td>-1,079998222</td><td>8,19335E-05</td><td>4,086538319</td><td>0,000808116</td><td>(GFP_MAGED1_HD - GFP)-(GFP_MAGED1_UN - GFP)</td></tr>
<tr><td>CNOT8</td><td>E5RIL2|Q9UFF9|Q9UFF9-3</td><td>CCR4-NOT transcription complex subun</td><td>-1,337389037</td><td>8,41904E-05</td><td>4,074737344</td><td>0,000826454</td><td>(GFP_MAGED1_HD - GFP)-(GFP_MAGED1_UN - GFP)</td></tr>
<tr><td>LAMB1</td><td>G3XAI2|P07942</td><td>Laminin subunit beta-1 OS=Homo sapie</td><td>-1,465245882</td><td>8,73995E-05</td><td>4,058491234</td><td>0,000847953</td><td>(GFP_MAGED1_HD - GFP)-(GFP_MAGED1_UN - GFP)</td></tr>
<tr><td>SLC25A14</td><td>F6SL11|O95258|O95258-2</td><td>Brain mitochondrial carrier protein 1 OS=</td><td>-1,008451166</td><td>9,24331E-05</td><td>4,034172669</td><td>0,000880135</td><td>(GFP_MAGED1_HD - GFP)-(GFP_MAGED1_UN - GFP)</td></tr>
<tr><td>SMG7</td><td>E9PD50|Q92540|Q92540-2|Q</td><td>Protein SMG7 OS=Homo sapiens GN=S</td><td>-1,380649943</td><td>9,42051E-05</td><td>4,025925394</td><td>0,000885082</td><td>(GFP_MAGED1_HD - GFP)-(GFP_MAGED1_UN - GFP)</td></tr>
<tr><td>CNOT7</td><td>Q9UIV1</td><td>CCR4-NOT transcription complex subun</td><td>-1,040563099</td><td>9,99268E-05</td><td>4,000318183</td><td>0,000920643</td><td>(GFP_MAGED1_HD - GFP)-(GFP_MAGED1_UN - GFP)</td></tr>
<tr><td>LDOC1</td><td>O95751</td><td>Protein LDOC1 OS=Homo sapiens GN=</td><td>-1,400874495</td><td>0,000107783</td><td>3,967448665</td><td>0,000970458</td><td>(GFP_MAGED1_HD - GFP)-(GFP_MAGED1_UN - GFP)</td></tr>
<tr><td>AP2B1</td><td>A0A087X253|P63010|P6301</td><td>AP-2 complex subunit beta OS=Homo sa</td><td>-1,017365919</td><td>0,000126124</td><td>3,899200756</td><td>0,001094153</td><td>(GFP_MAGED1_HD - GFP)-(GFP_MAGED1_UN - GFP)</td></tr>
<tr><td>NDUFA9</td><td>Q16795</td><td>NADH dehydrogenase [ubiquinone] 1 al</td><td>-1,387983512</td><td>0,000127325</td><td>3,895086875</td><td>0,00110055</td><td>(GFP_MAGED1_HD - GFP)-(GFP_MAGED1_UN - GFP)</td></tr>
<tr><td>HGS</td><td>O14964</td><td>Hepatocyte growth factor-regulated tyros</td><td>-1,089435875</td><td>0,00013123</td><td>3,881967567</td><td>0,001126113</td><td>(GFP_MAGED1_HD - GFP)-(GFP_MAGED1_UN - GFP)</td></tr>
<tr><td>ZDHHC17</td><td>Q8IUH5</td><td>Palmitoyltransferase ZDHHC17 OS=Hor</td><td>-1,229472028</td><td>0,000136911</td><td>3,863560702</td><td>0,001166445</td><td>(GFP_MAGED1_HD - GFP)-(GFP_MAGED1_UN - GFP)</td></tr>
<tr><td>PABPC1</td><td>P11940</td><td>Polyadenylate-binding protein 1 OS=Ho</td><td>-1,077365524</td><td>0,000147744</td><td>3,830489051</td><td>0,001230866</td><td>(GFP_MAGED1_HD - GFP)-(GFP_MAGED1_UN - GFP)</td></tr>
<tr><td>ATXN2</td><td>F8VQP2|F8WB06|H0YH87|Q</td><td>Ataxin-2 OS=Homo sapiens GN=ATXN2</td><td>-1,160377741</td><td>0,000153852</td><td>3,812895968</td><td>0,001265422</td><td>(GFP_MAGED1_HD - GFP)-(GFP_MAGED1_UN - GFP)</td></tr>
<tr><td>RALY</td><td>Q5QPL9|Q5QPM0|Q9UKM9|</td><td>RNA-binding protein Raly (Fragment) OS</td><td>-1,113337365</td><td>0,000160249</td><td>3,795203421</td><td>0,001308979</td><td>(GFP_MAGED1_HD - GFP)-(GFP_MAGED1_UN - GFP)</td></tr>
<tr><td>GK|GK3P</td><td>P32189|P32189-1|P32189-2</td><td>Glycerol kinase OS=Homo sapiens GN=</td><td>-1,009120874</td><td>0,000163375</td><td>3,786814406</td><td>0,00132994</td><td>(GFP_MAGED1_HD - GFP)-(GFP_MAGED1_UN - GFP)</td></tr>
<tr><td>EXPH5</td><td>Q8NEV8|Q8NEV8-2</td><td>Exophilin-5 OS=Homo sapiens GN=EXF</td><td>-1,55134363</td><td>0,000176129</td><td>3,754167951</td><td>0,001409629</td><td>(GFP_MAGED1_HD - GFP)-(GFP_MAGED1_UN - GFP)</td></tr>
<tr><td>MLF2</td><td>Q15773</td><td>Myeloid leukemia factor 2 OS=Homo sap</td><td>-1,163305079</td><td>0,000186514</td><td>3,729289639</td><td>0,001468022</td><td>(GFP_MAGED1_HD - GFP)-(GFP_MAGED1_UN - GFP)</td></tr>
<tr><td>ATAD1</td><td>Q8NBU5</td><td>ATPase family AAA domain-containing p</td><td>-1,028842557</td><td>0,00021186</td><td>3,673951265</td><td>0,001608917</td><td>(GFP_MAGED1_HD - GFP)-(GFP_MAGED1_UN - GFP)</td></tr>
<tr><td>SAV1</td><td>H0YJH0|Q9H4B6</td><td>Protein salvador homolog 1 (Fragment) (</td><td>-1,472282087</td><td>0,00022539</td><td>3,647066085</td><td>0,001673216</td><td>(GFP_MAGED1_HD - GFP)-(GFP_MAGED1_UN - GFP)</td></tr>
<tr><td>FAM57A</td><td>Q8TBR7|Q8TBR7-1</td><td>Protein FAM57A OS=Homo sapiens GN</td><td>-1,176202268</td><td>0,000347995</td><td>3,458426474</td><td>0,002336681</td><td>(GFP_MAGED1_HD - GFP)-(GFP_MAGED1_UN - GFP)</td></tr>
<tr><td>TOMM40</td><td>O96008|O96008-2</td><td>Mitochondrial import receptor subunit TO</td><td>-1,400627252</td><td>0,00035595</td><td>3,448611139</td><td>0,002376665</td><td>(GFP_MAGED1_HD - GFP)-(GFP_MAGED1_UN - GFP)</td></tr>
<tr><td>HNRNPA1</td><td>F8W6I7|P09651|P09651-2|P</td><td>Heterogeneous nuclear ribonucleoprote</td><td>-1,013402869</td><td>0,000411276</td><td>3,385867014</td><td>0,002656528</td><td>(GFP_MAGED1_HD - GFP)-(GFP_MAGED1_UN - GFP)</td></tr>
<tr><td>FAM83D</td><td>A0A087WXK8|Q9H4H8|Q9H</td><td>Chromosome 20 open reading frame 12</td><td>-1,377434886</td><td>0,000433688</td><td>3,362822426</td><td>0,002771174</td><td>(GFP_MAGED1_HD - GFP)-(GFP_MAGED1_UN - GFP)</td></tr>
<tr><td>SLC35A4</td><td>L0R6Q1</td><td>Alternative protein SLC35A4 OS=Homo</td><td>-1,255007073</td><td>0,000481034</td><td>3,317824504</td><td>0,002985423</td><td>(GFP_MAGED1_HD - GFP)-(GFP_MAGED1_UN - GFP)</td></tr>
<tr><td>MLF1</td><td>C9J3M3|C9K0D3|P58340|P5</td><td>Myeloid leukemia factor 1 (Fragment) OS</td><td>-1,20240436</td><td>0,000671532</td><td>3,172933101</td><td>0,003800553</td><td>(GFP_MAGED1_HD - GFP)-(GFP_MAGED1_UN - GFP)</td></tr>
<tr><td>XRN1</td><td>Q8IZH2|Q8IZH2-2</td><td>5'-3' exoribonuclease 1 OS=Homo sapie</td><td>-1,076952434</td><td>0,000725122</td><td>3,139589014</td><td>0,004027137</td><td>(GFP_MAGED1_HD - GFP)-(GFP_MAGED1_UN - GFP)</td></tr>
<tr><td>BTBD2</td><td>Q9BX70</td><td>BTB/POZ domain-containing protein 2 O</td><td>-1,643710063</td><td>0,00100759</td><td>2,996716166</td><td>0,005102899</td><td>(GFP_MAGED1_HD - GFP)-(GFP_MAGED1_UN - GFP)</td></tr>
<tr><td>PEF1</td><td>Q9UBV8</td><td>Peflin OS=Homo sapiens GN=PEF1 PE=</td><td>-1,548064023</td><td>0,001008987</td><td>2,996114393</td><td>0,005102899</td><td>(GFP_MAGED1_HD - GFP)-(GFP_MAGED1_UN - GFP)</td></tr>
<tr><td>ATAD3B</td><td>Q5T9A4</td><td>ATPase family AAA domain-containing p</td><td>-1,176029039</td><td>0,001085775</td><td>2,964260003</td><td>0,005399348</td><td>(GFP_MAGED1_HD - GFP)-(GFP_MAGED1_UN - GFP)</td></tr>
<tr><td>PRRC2A</td><td>P48634|P48634-2|P48634-3</td><td>Protein PRRC2A OS=Homo sapiens GN</td><td>-1,07200831</td><td>0,00109869</td><td>2,959124952</td><td>0,005440803</td><td>(GFP_MAGED1_HD - GFP)-(GFP_MAGED1_UN - GFP)</td></tr>
<tr><td>SLC25A19</td><td>Q9HC21|Q9HC21-2</td><td>Mitochondrial thiamine pyrophosphate c</td><td>-1,196704806</td><td>0,001179526</td><td>2,928292365</td><td>0,005710253</td><td>(GFP_MAGED1_HD - GFP)-(GFP_MAGED1_UN - GFP)</td></tr>
<tr><td>GRB2</td><td>P62993</td><td>Growth factor receptor-bound protein 2 (</td><td>-1,353776378</td><td>0,001200203</td><td>2,920745328</td><td>0,005798541</td><td>(GFP_MAGED1_HD - GFP)-(GFP_MAGED1_UN - GFP)</td></tr>
<tr><td>LARP4B</td><td>Q92615</td><td>La-related protein 4B OS=Homo sapiens</td><td>-1,202706207</td><td>0,00138474</td><td>2,858631919</td><td>0,006568025</td><td>(GFP_MAGED1_HD - GFP)-(GFP_MAGED1_UN - GFP)</td></tr>
<tr><td>BAG4</td><td>O95429|O95429-2</td><td>BAG family molecular chaperone regula</td><td>-1,236977864</td><td>0,001387105</td><td>2,857890638</td><td>0,006568025</td><td>(GFP_MAGED1_HD - GFP)-(GFP_MAGED1_UN - GFP)</td></tr>
<tr><td>RBFOX2</td><td>O43251-6|O43251-8</td><td>Isoform 6 of RNA binding protein fox-1 h</td><td>-1,441615693</td><td>0,001454798</td><td>2,837197199</td><td>0,006793823</td><td>(GFP_MAGED1_HD - GFP)-(GFP_MAGED1_UN - GFP)</td></tr>
<tr><td>RWDD2B</td><td>P57060</td><td>RWD domain-containing protein 2B OS=</td><td>-1,063449012</td><td>0,001634868</td><td>2,786517436</td><td>0,007346087</td><td>(GFP_MAGED1_HD - GFP)-(GFP_MAGED1_UN - GFP)</td></tr>
<tr><td>PRDX3</td><td>P30048|P30048-2</td><td>Thioredoxin-dependent peroxide reduct</td><td>-1,00647912</td><td>0,002039404</td><td>2,690496754</td><td>0,00854967</td><td>(GFP_MAGED1_HD - GFP)-(GFP_MAGED1_UN - GFP)</td></tr>
<tr><td>NDUFB6</td><td>A0A087WZX2|O95139</td><td>NADH dehydrogenase [ubiquinone] 1 be</td><td>-1,00426314</td><td>0,002905359</td><td>2,536800221</td><td>0,011174819</td><td>(GFP_MAGED1_HD - GFP)-(GFP_MAGED1_UN - GFP)</td></tr>
<tr><td>CSRP2</td><td>F8VW96|Q16527</td><td>Cysteine and glycine-rich protein 2 OS=</td><td>-1,440089506</td><td>0,003798349</td><td>2,420405153</td><td>0,013558612</td><td>(GFP_MAGED1_HD - GFP)-(GFP_MAGED1_UN - GFP)</td></tr>
<tr><td>PFN2</td><td>C9J2N0|C9J712|P35080</td><td>Profilin OS=Homo sapiens GN=PFN2 PE</td><td>-1,38488829</td><td>0,004676542</td><td>2,330075198</td><td>0,015767574</td><td>(GFP_MAGED1_HD - GFP)-(GFP_MAGED1_UN - GFP)</td></tr>
<tr><td>FN3KRP</td><td>Q9HA64</td><td>Ketosamine-3-kinase OS=Homo sapiens</td><td>-1,048963625</td><td>0,006600639</td><td>2,180414022</td><td>0,020617239</td><td>(GFP_MAGED1_HD - GFP)-(GFP_MAGED1_UN - GFP)</td></tr>
<tr><td>PPIA</td><td>P62937</td><td>Peptidyl-prolyl cis-trans isomerase A OS</td><td>-1,206949517</td><td>0,009171757</td><td>2,037547463</td><td>0,026522222</td><td>(GFP_MAGED1_HD - GFP)-(GFP_MAGED1_UN - GFP)</td></tr>
<tr><td>CKB</td><td>P12277</td><td>Creatine kinase B-type OS=Homo sapie</td><td>-1,061356515</td><td>0,013101734</td><td>1,882671212</td><td>0,034757614</td><td>(GFP_MAGED1_HD - GFP)-(GFP_MAGED1_UN - GFP)</td></tr>
<tr><td>EIF2S1</td><td>P05198</td><td>Eukaryotic translation initiation factor 2 s</td><td>-1,029900287</td><td>0,023406977</td><td>1,630654663</td><td>0,054334361</td><td>(GFP_MAGED1_HD - GFP)-(GFP_MAGED1_UN - GFP)</td></tr>
<tr><td>ARG1</td><td>P05089|P05089-2</td><td>Arginase-1 OS=Homo sapiens GN=ARG</td><td>-1,134714432</td><td>0,039697801</td><td>1,401233551</td><td>0,082773397</td><td>(GFP_MAGED1_HD - GFP)-(GFP_MAGED1_UN - GFP)</td></tr>
<tr><td>BASP1</td><td>P80723</td><td>Brain acid soluble protein 1 OS=Homo s</td><td>0,8255626</td><td>0,0028439</td><td>2,5460835</td><td>0,0111085</td><td>(GFP_MAGED1_HD - GFP)-(GFP_MAGED1_UN - GFP)</td></tr>
</table>

**Supplementary table 7:** Proteomics with reported hits and candidates of GFP immunoprecipitation of GFP_MAGED1_MHD versus GFP_MAGE-D1_UNS. over-expressed in HEK cells. Related to Figure 4A.

| Gene | Protein ID | Protein Description | log2(FC) | pvalue | -log10(pvalue) | fdr | Comparison |
|---|---|---|---|---|---|---|---|
| PRAJA-2 | O43164 | E3 ubiquitin-protein ligase Praja-2 | 1,850306794 | 6,89E-09 | 8,16184986 | 1,77E-05 | (MAGED1_WT - GFP) - (MAGED1_V569D-GFP) |
| PRAJA-1 | Q8NG27 | E3 ubiquitin-protein ligase Praja-1 | 1,768903447 | 5,98E-08 | 7,223069977 | 7,68E-05 | (MAGED1_WT - GFP) - (MAGED1_V569D-GFP) |
| PEBP1 | P30086 | Phosphatidylethanolamine-binding protein 1 | 1,225387416 | 9,85E-05 | 4,006750902 | 0,04212102 | (MAGED1_WT - GFP) - (MAGED1_V569D-GFP) |
| DNAJA3 | Q96EY1 | DnaJ homolog subfamily A member 3, mitochondrial | 1,080754326 | 4,04E-05 | 4,393126198 | 0,020763704 | (MAGED1_WT - GFP) - (MAGED1_V569D-GFP) |
| CFL1 | P23528 | Cofilin-1 | 1,048419598 | 2,13E-05 | 4,672191731 | 0,013650524 | (MAGED1_WT - GFP) - (MAGED1_V569D-GFP) |
| TBCA | O75347 | Tubulin-specific chaperone A | 1,359497773 | 0,001123563 | 2,949402669 | 0,15998044 | (MAGED1_WT - GFP) - (MAGED1_V569D-GFP) |
| CEP44 | Q9C0F1 | Centrosomal protein of 44 kDa | 1,331039574 | 0,000578266 | 3,237872492 | 0,123693618 | (MAGED1_WT - GFP) - (MAGED1_V569D-GFP) |
| ARL2BP | Q9Y2Y0 | ADP-ribosylation factor-like protein 2-binding protein | 1,159935476 | 0,000481419 | 3,317477204 | 0,114119368 | (MAGED1_WT - GFP) - (MAGED1_V569D-GFP) |
| CALML5 | Q9NZT1 | Calmodulin-like protein 5 | -1,461500695 | 2,07E-05 | 4,684852528 | 0,013452117 | (MAGED1_WT - GFP) - (MAGED1_V569D-GFP) |
| JAK1 | P23458 | Tyrosine-protein kinase JAK1 | -1,982406592 | 0,000781377 | 3,107139555 | 0,140109267 | (MAGED1_WT - GFP) - (MAGED1_V569D-GFP) |
| TUFT1 | Q9NNX1 | Tuftelin | -1,431803371 | 0,000966244 | 3,014913076 | 0,151027028 | (MAGED1_WT - GFP) - (MAGED1_V569D-GFP) |
| NUP35 | Q8NFH5 | Nucleoporin NUP35 | -1,191092076 | 0,001767697 | 2,752592114 | 0,196751325 | (MAGED1_WT - GFP) - (MAGED1_V569D-GFP) |

**Supplementary table 8:** Proteomics with reported hits and candidates of GFP immunoprecipitation of GFP_MAGED1_WT versus GFP_MAGE-D1_V569D mutant over-expressed in HEK cells. Related to Figure 4F

| Gene | Protein ID | Protein Description | log2(FC) | pvalue | -log10(pvalue) | fdr | Comparison |
|---|---|---|---|---|---|---|---|
| **MAGE-D4** | Q96JG8 | Melanoma-associated antigen D4 | 1,700055153 | 2,22E-16 | 1,57E+01 | 1,48E-13 | (PRAJA1_HA_WT - HA) - (PJA1_HA_F514D_F517D-HA) |
| **MAGE-D1** | Q9Y5V3 | Melanoma-associated antigen D1 | 2,602865375 | 2,22E-16 | 1,57E+01 | 1,48E-13 | (PRAJA1_HA_WT - HA) - (PJA1_HA_F514D_F517D-HA) |
| **USP7** | Q93009 | Ubiquitin carboxyl-terminal hydrolase 7 | 0,968722201 | 2,22E-16 | 15,654 | 1,48E-13 | (PRAJA1_HA_WT - HA) - (PJA1_HA_F514D_F517D-HA) |

**Supplementary table 9:** Proteomics with reported hits and candidates of HA immunoprecipitation of Praja-1_HA_WT versus Praja-1_F514D_F517D_HA over-expressed in HEK cells. Related to Supplementary Figure 4F.

| Gene | Protein.ID | Protein.Description | -log2(FC) | pvalue | -log10(pvalue) | fdr | Comparison |
|---|---|---|---|---|---|---|---|
| **MAGE-D1** | Q9Y5V3 | Melanoma-associated antigen D1 | 1,523348879 | 2,22E-16 | 15,6536 | 3,14E-13 | (PRAJA2_HA_WT - HA)-(PJA2_HA_F553D_F556D-HA) |

**Supplementary table 10:** Proteomics with reported hits and candidates of HA immunoprecipitation of Praja-2_HA_WT versus Praja-2_F553D_F556D_HA over-expressed in HEK cells. Related to Supplementary Figure 4G.

| Peptide Sequence | Modified Sequence | Prev AA | Next AA | Start | End | Peptide Length | Charges | Assigned Modifications | Protein | Protein ID | B3 Spectral Count | B6 Spectral Count | B3 Intensity | B6 Intensity | B3 Match Type | B6 Match Type | B3 Localization | B6 Localization |
|---|---|---|---|---|---|---|---|---|---|---|---|---|---|---|---|---|---|---|
| AAEAAAAPAES | AAEAAAAPAES | K | E | 122 | 143 | 22 | 2,3 | | sp\|P4452_BA | P4452_BASP | 3 | 1 | 3,02E+07 | 3525165,8 | MS/MS | MS/MS | | |
| AAEAAAAPAES | AAEAAAAPAES | K | K | 122 | 149 | 28 | 2,3,4 | | sp\|P4452_BA | P4452_BASP | 21 | 47 | 1,02E+10 | 2,21E+09 | MS/MS | MS/MS | | |
| AAEAAAAPAES | AAEAAAAPAES | K | T | 122 | 150 | 29 | 2,3,4 | | sp\|P4452_BA | P4452_BASP | 8 | 8 | 3,68E+08 | 1,05E+09 | MS/MS | MS/MS | | |
| AAEAAAAPAES | AAEAAAAPAES | K | S | 122 | 163 | 42 | 3,4,5 | | sp\|P4452_BA | P4452_BASP | 16 | 31 | 6,78E+09 | 1,88E+10 | MS/MS | MS/MS | | |
| AAEAAAAPAES | AAEAAAAPAES | K | S | 122 | 163 | 42 | 5 | 29K(114.0429) | sp\|P4452_BA | P4452_BASP | 0 | 1 | 0 | 3557944 | unmatched | MS/MS | | K:114.0429@AAEAAAAPAESAAPAAGEEPSK(0.0123)EEGEPK(0.0375)K(0.9429)TEAPAAPAAQETK(0.0073); |
| AAEAAAAPAES | AAEAAAAPAES | K | S | 122 | 163 | 42 | 5 | 28K(114.0429) | sp\|P4452_BA | P4452_BASP | 0 | 1 | 0 | 3557944 | unmatched | MS/MS | | K:114.0429@AAEAAAAPAESAAPAAGEEPSK(0.0067)EEGEPK(0.4943)K(0.4943)TEAPAAPAAQETK(0.0046); |
| AAEAAAAPAES | AAEAAAAPAES | K | K | 122 | 149 | 28 | 3 | 22K(114.0429) | sp\|P4452_BA | P4452_BASP | 0 | 4 | 0 | 1039640,25 | unmatched | MS/MS | | K:114.0429@AAEAAAAPAESAAPAAGEEPSK(0.9701)EEGEPK(0.0299); |
| AAEAAAAPAES | AAEAAAAPAES | K | S | 122 | 163 | 42 | 5 | 22K(114.0429) | sp\|P4452_BA | P4452_BASP | 0 | 1 | 0 | 3557944 | unmatched | MS/MS | | K:114.0429@AAEAAAAPAESAAPAAGEEPSK(0.3319)EEGEPK(0.3319)K(0.3319)TEAPAAPAAQETK(0.0042); |
| AAEAAAAPAES | AAEAAAAPAES | K | K | 122 | 149 | 28 | 4 | 22K(114.0429),28K | sp\|P4452_BA | P4452_BASP | 0 | 1 | 0 | 0 | unmatched | MS/MS | | K:114.0429@AAEAAAAPAESAAPAAGEEPSK(1.0000)EEGEPK(1.0000); |
| AAEAAAAPAES | AAEAAAAPAES | K | T | 122 | 150 | 29 | 4 | 22K(114.0429),28K | sp\|P4452_BA | P4452_BASP | 0 | 1 | 0 | 0 | unmatched | MS/MS | | K:114.0429@AAEAAAAPAESAAPAAGEEPSK(0.6667)EEGEPK(0.6667)K(0.6667); |
| AEEKEGEKDA | AEEKEGEKDA | K | A | 66 | 84 | 19 | 2,3,4,5 | | sp\|P4452_BA | P4452_BASP | 5 | 5 | 6,85E+07 | 3,40E+08 | MS/MS | MS/MS | | |
| AEGAATEEEG | AEGAATEEEG | K | E | 26 | 38 | 13 | 2 | | sp\|P4452_BA | P4452_BASP | 4 | 5 | 1,50E+09 | 5,63E+09 | MS/MS | MS/MS | | |
| AEGAATEEEG | AEGAATEEEG | K | E | 26 | 52 | 27 | 2,3 | | sp\|P4452_BA | P4452_BASP | 10 | 13 | 1,65E+09 | 9,20E+07 | MS/MS | MS/MS | | |
| AEGAATEEEG | AEGAATEEEG | K | E | 26 | 55 | 30 | 2,3,4 | | sp\|P4452_BA | P4452_BASP | 13 | 13 | 1,88E+09 | 6,61E+09 | MS/MS | MS/MS | | |
| AEGAATEEEG | AEGAATEEEG | K | A | 26 | 65 | 40 | 4,5,6 | | sp\|P4452_BA | P4452_BASP | 5 | 5 | 4,79E+07 | 7,06E+07 | MS/MS | MS/MS | | |
| AEGAATEEEG | AEGAATEEEG | K | E | 26 | 52 | 27 | 2,3 | 13K(114.0429) | sp\|P4452_BA | P4452_BASP | 2 | 2 | 0 | 0 | MS/MS | MS/MS | K:114.0429@ | K:114.0429@AEGAATEEEGTPK(0.9791)ESEPQAAAEPAEAK(0.0209); |
| AEPEKTEGAA | AEPEKTEGAA | K | A | 85 | 97 | 13 | 2,3 | | sp\|P4452_BA | P4452_BASP | 4 | 5 | 4,01E+08 | 3,79E+09 | MS/MS | MS/MS | | |
| AEPEKTEGAA | AEPEKTEGAA | K | A | 85 | 121 | 37 | 3,4,5 | | sp\|P4452_BA | P4452_BASP | 19 | 21 | 2,61E+09 | 9,01E+09 | MS/MS | MS/MS | | |
| AEPPKAPEQE | AEPPKAPEQE | K | A | 98 | 121 | 24 | 2,3,4 | | sp\|P4452_BA | P4452_BASP | 22 | 47 | 6,63E+09 | 1,93E+10 | MS/MS | MS/MS | | |
| AEPPKAPEQE | AEPPKAPEQE | K | E | 98 | 143 | 46 | 4 | | sp\|P4452_BA | P4452_BASP | 1 | 0 | 2895243,8 | 0 | MS/MS | unmatched | | |
| AEPPKAPEQE | AEPPK[114.0 | K | A | 98 | 121 | 24 | 3 | 5K(114.0429) | sp\|P4452_BA | P4452_BASP | 1 | 1 | 0 | 3473214,5 | MS/MS | MS/MS | K:114.0429@ | K:114.0429@AEPPK(0.9715)APEQEQAAPGPAAGGEAPK(0.0285); |
| APEQEQAAP( | APEQEQAAP( | K | A | 103 | 121 | 19 | 2,3 | | sp\|P4452_BA | P4452_BASP | 4 | 5 | 1,10E+08 | 3,44E+08 | MS/MS | MS/MS | | |
| APEQEQAAP( | APEQEQAAP( | K | K | 103 | 149 | 47 | 4 | | sp\|P4452_BA | P4452_BASP | 2 | 0 | 4244403 | 0 | MS/MS | unmatched | | |
| AQGPAASAEE | AQGPAASAEE | K | E | 199 | 226 | 28 | 2,3 | | sp\|P4452_BA | P4452_BASP | 5 | 8 | 1,47E+08 | 9,35E+08 | MS/MS | MS/MS | | |
| AQGPAASAEE | AQGPAASAEE | K | - | 199 | 227 | 29 | 2,3,4 | | sp\|P4452_BA | P4452_BASP | 68 | 95 | 1,75E+10 | 1,10E+10 | MS/MS | MS/MS | | |
| AQGPAASAEE | AQGPAASAEE | K | E | 199 | 226 | 28 | 4 | 28K(114.0429) | sp\|P4452_BA | P4452_BASP | 1 | 0 | 4158789 | 0 | MS/MS | unmatched | K:114.0429@AQGPAASAEEPK(0.0294)PVEAPAANSDQTVTVK(0.9706); | |
| AQGPAASAEE | AQGPAASAEE | K | E | 199 | 226 | 28 | 3 | 12K(114.0429) | sp\|P4452_BA | P4452_BASP | 0 | 1 | 0 | 0 | unmatched | MS/MS | | K:114.0429@AQGPAASAEEPK(0.9780)PVEAPAANSDQTVTVK(0.0220); |
| AQGPAASAEE | AQGPAASAEE | K | - | 199 | 227 | 29 | 3 | 12K(114.0429) | sp\|P4452_BA | P4452_BASP | 3 | 1 | 2103061,8 | 0 | MS/MS | MS/MS | K:114.0429@ | K:114.0429@AQGPAASAEEPK(0.8973)PVEAPAANSDQTVTVK(0.1027)E; |
| AQGPAASAEE | AQGPAASAEE | K | E | 199 | 226 | 28 | 2 | 12K(114.0429),28K | sp\|P4452_BA | P4452_BASP | 0 | 1 | 0 | 0 | unmatched | MS/MS | | K:114.0429@AQGPAASAEEPK(1.0000)PVEAPAANSDQTVTVK(1.0000); |
| AQGPAASAEE | AQGPAASAEE | K | - | 199 | 227 | 29 | 3 | 12K(114.0429),28K | sp\|P4452_BA | P4452_BASP | 0 | 1 | 0 | 1933836,1 | unmatched | MS/MS | | K:114.0429@AQGPAASAEEPK(1.0000)PVEAPAANSDQTVTVK(1.0000)E; |
| DAAAAKEEAP | DAAAAKEEAP | K | A | 74 | 97 | 24 | 2,3,4 | | sp\|P4452_BA | P4452_BASP | 2 | 4 | 4,23E+07 | 3,90E+08 | MS/MS | MS/MS | | |
| DKKAEGAATE | DKKAEGAATE | K | E | 23 | 38 | 16 | 2,3 | | sp\|P4452_BA | P4452_BASP | 4 | 5 | 3,91E+08 | 2,59E+09 | MS/MS | MS/MS | | |
| DKKAEGAATE | DKKAEGAATE | K | E | 23 | 52 | 30 | 2,3,4,5 | | sp\|P4452_BA | P4452_BASP | 17 | 25 | 2,11E+09 | 1,43E+10 | MS/MS | MS/MS | | |
| DKKAEGAATE | DKK[114.042 | K | E | 23 | 52 | 30 | 4 | 3K(114.0429) | sp\|P4452_BA | P4452_BASP | 0 | 1 | 0 | 3899981,5 | unmatched | MS/MS | | K:114.0429@DK(0.0982)K(0.8893)AEGAATEEEGTPK(0.0057)ESEPQAAAEPAEAK(0.0068); |
| DKKAEGAATE | DK[114.0429 | K | E | 23 | 38 | 16 | 3 | 2K(114.0429) | sp\|P4452_BA | P4452_BASP | 0 | 2 | 0 | 1,16E+07 | unmatched | MS/MS | | K:114.0429@DK(0.9785)K(0.0106)AEGAATEEEGTPK(0.0109); |
| EEAPKAEPEK | EEAPKAEPEK | K | A | 80 | 97 | 18 | 2,3,4 | | sp\|P4452_BA | P4452_BASP | 5 | 6 | 3,03E+08 | 3,78E+09 | MS/MS | MS/MS | | |
| EEGEPKKTEA | EEGEPKKTEA | K | S | 144 | 163 | 20 | 3 | | sp\|P4452_BA | P4452_BASP | 1 | 1 | 667007 | 1274676,6 | MS/MS | MS/MS | | |
| EGEKDAAAAK | EGEKDAAAAK | K | A | 70 | 84 | 15 | 2,3,4 | | sp\|P4452_BA | P4452_BASP | 5 | 7 | 4,75E+07 | 6,87E+08 | MS/MS | MS/MS | | |
| EGEKDAAAAK | EGEKDAAAAK | K | T | 70 | 89 | 20 | 3,4 | | sp\|P4452_BA | P4452_BASP | 0 | 2 | 0 | 2,44E+07 | unmatched | MS/MS | | |
| EGKEKPDQD | EGKEKPDQD | K | E | 53 | 69 | 17 | 2,3 | | sp\|P4452_BA | P4452_BASP | 3 | 3 | 1,03E+08 | 2,05E+09 | MS/MS | MS/MS | | |
| EGKEKPDQD | EGKEKPDQD | K | D | 53 | 73 | 21 | 3,5 | | sp\|P4452_BA | P4452_BASP | 1 | 3 | 1,03E+07 | 9,46E+07 | MS/MS | MS/MS | | |
| EGKEKPDQD | EGKEKPDQD | K | D | 53 | 73 | 21 | 3 | 21K(114.0429) | sp\|P4452_BA | P4452_BASP | 0 | 1 | 0 | 0 | unmatched | MS/MS | | K:114.0429@EGK(0.0298)EK(0.0522)PDQDAEGK(0.0088)AEEK(0.0065)EGEK(0.9027); |
| EKDKKAEGAA | EKDKKAEGAA | K | E | 21 | 38 | 18 | 2,3,4 | | sp\|P4452_BA | P4452_BASP | 10 | 8 | 1,12E+09 | 5,36E+09 | MS/MS | MS/MS | | |
| EKDKKAEGAA | EKDK[114.04 | K | E | 21 | 38 | 18 | 3 | 4K(114.0429) | sp\|P4452_BA | P4452_BASP | 0 | 1 | 0 | 2,91E+07 | unmatched | MS/MS | | K:114.0429@EK(0.0698)DK(0.4621)K(0.4621)AEGAATEEEGTPK(0.0060); |
| EKPDQDAEG | EKPDQDAEG | K | E | 56 | 69 | 14 | 2,3 | | sp\|P4452_BA | P4452_BASP | 4 | 4 | 3,10E+08 | 4,50E+09 | MS/MS | MS/MS | | |
| EKPDQDAEG | EKPDQDAEG | K | D | 56 | 73 | 18 | 4 | | sp\|P4452_BA | P4452_BASP | 1 | 3 | 1,55E+07 | 1,36E+08 | MS/MS | MS/MS | | |
| EKPDQDAEG | EKPDQDAEG | K | E | 56 | 79 | 24 | 2,3,4,5 | | sp\|P4452_BA | P4452_BASP | 8 | 14 | 3,62E+08 | 5,94E+09 | MS/MS | MS/MS | | |
| EKPDQDAEG | EK[114.0429] | K | E | 56 | 79 | 24 | 3,5 | 10K(114.0429),14K | sp\|P4452_BA | P4452_BASP | 0 | 2 | 0 | 1,27E+07 | unmatched | MS/MS | | K:114.0429@EK(0.9628)PDQDAEGK(0.9631)AEEK(0.9631)EGEK(0.0755)DAAAAK(0.0356); |
| ESEPQAAAEP | ESEPQAAAEP | K | E | 39 | 52 | 14 | 2 | | sp\|P4452_BA | P4452_BASP | 8 | 10 | 7,35E+08 | 4,36E+09 | MS/MS | MS/MS | | |
| ESEPQAAAEP | ESEPQAAAEP | K | E | 39 | 55 | 17 | 2,3 | | sp\|P4452_BA | P4452_BASP | 8 | 12 | 2,63E+08 | 3,09E+07 | MS/MS | MS/MS | | |
| ESEPQAAAEP | ESEPQAAAEP | K | A | 39 | 65 | 27 | 3,4,5 | | sp\|P4452_BA | P4452_BASP | 5 | 6 | 3,25E+07 | 1,07E+08 | MS/MS | MS/MS | | |
| ESEPQAAAEP | ESEPQAAAEP | K | E | 39 | 69 | 31 | 3,4,5,6 | | sp\|P4452_BA | P4452_BASP | 6 | 6 | 4,96E+07 | 2,16E+08 | MS/MS | MS/MS | | |
| ESEPQAAAEP | ESEPQAAAEP | K | E | 39 | 69 | 31 | 6 | 14K(114.0429),17K | sp\|P4452_BA | P4452_BASP | 0 | 1 | 0 | 0 | unmatched | MS/MS | | K:114.0429@ESEPQAAAEPAEAK(0.9124)EGK(0.9124)EK(0.9126)PDQDAEGK(0.1286)AEEK(0.1340); |
| ETPAATEAPSS | ETPAATEAPSS | K | A | 185 | 198 | 14 | 2,3 | | sp\|P4452_BA | P4452_BASP | 12 | 26 | 1,35E+10 | 7,08E+09 | MS/MS | MS/MS | | |
| ETPAATEAPSS | ETPAATEAPSS | K | E | 185 | 226 | 42 | 4 | | sp\|P4452_BA | P4452_BASP | 3 | 3 | 7781301 | 0 | MS/MS | MS/MS | | |
| ETPAATEAPSS | ETPAATEAPSS | K | - | 185 | 227 | 43 | 3,4,5 | | sp\|P4452_BA | P4452_BASP | 12 | 3 | 2,21E+09 | 0 | MS/MS | MS/MS | | |
| ETPAATEAPSS | ETPAATEAPSS | K | - | 185 | 227 | 43 | 4 | 26K(114.0429) | sp\|P4452_BA | P4452_BASP | 0 | 1 | 0 | 9164933 | unmatched | MS/MS | | K:114.0429@ETPAATEAPSSTPK(0.4586)AQGPAASAEEPK(0.5342)PVEAPAANSDQTVTVK(0.0072)E; |
| ETPAATEAPSS | ETPAATEAPSS | K | - | 185 | 227 | 43 | 4 | 14K(114.0429) | sp\|P4452_BA | P4452_BASP | 0 | 2 | 0 | 9164933 | unmatched | MS/MS | | K:114.0429@ETPAATEAPSSTPK(0.9579)AQGPAASAEEPK(0.0287)PVEAPAANSDQTVTVK(0.0134)E; |
| ETPAATEAPSS | ETPAATEAPSS | K | E | 185 | 226 | 42 | 3 | 14K(114.0429),26K | sp\|P4452_BA | P4452_BASP | 0 | 2 | 0 | 0 | unmatched | MS/MS | | K:114.0429@ETPAATEAPSSTPK(0.9453)AQGPAASAEEPK(0.9453)PVEAPAANSDQTVTVK(0.1093); |
| GYNVNDEK | GYNVNDEK | K | A | 11 | 18 | 8 | 2 | | sp\|P4452_BA | P4452_BASP | 1 | 2 | 1,58E+07 | 3,03E+07 | MS/MS | MS/MS | | |
| GYNVNDEKA | GYNVNDEKA | K | E | 11 | 20 | 10 | 2,3 | | sp\|P4452_BA | P4452_BASP | 8 | 8 | 2,47E+09 | 7,09E+09 | MS/MS | MS/MS | | |
| GYNVNDEKA | GYNVNDEKA | K | D | 11 | 22 | 12 | 2,3 | | sp\|P4452_BA | P4452_BASP | 7 | 8 | 2,06E+09 | 7,26E+09 | MS/MS | MS/MS | | |
| GYNVNDEKA | GYNVNDEKA | K | K | 11 | 24 | 14 | 2,3,4 | | sp\|P4452_BA | P4452_BASP | 17 | 13 | 6,50E+08 | 1,20E+09 | MS/MS | MS/MS | | |
| GYNVNDEKA | GYNVNDEKA | K | D | 11 | 22 | 12 | 3 | 10K(114.0429) | sp\|P4452_BA | P4452_BASP | 0 | 3 | 0 | 1,81E+07 | unmatched | MS/MS | | K:114.0429@GYNVNDEK(0.0125)AK(0.9769)EK(0.0105); |
| GYNVNDEKA | GYNVNDEKA | K | K | 11 | 24 | 14 | 3 | 10K(114.0429) | sp\|P4452_BA | P4452_BASP | 0 | 1 | 0 | 480821,12 | unmatched | MS/MS | | K:114.0429@GYNVNDEK(0.3265)AK(0.6534)EK(0.0086)DK(0.0116); |
| GYNVNDEKA | GYNVNDEK[1 | K | E | 11 | 20 | 10 | 2,3 | 8K(114.0429) | sp\|P4452_BA | P4452_BASP | 0 | 3 | 0 | 8,93E+07 | unmatched | MS/MS | | K:114.0429@GYNVNDEK(0.9820)AK(0.0180); |
| GYNVNDEKA | GYNVNDEK[1 | K | D | 11 | 22 | 12 | 2 | 8K(114.0429) | sp\|P4452_BA | P4452_BASP | 0 | 3 | 0 | 2,12E+07 | unmatched | MS/MS | | K:114.0429@GYNVNDEK(0.9831)AK(0.0086)EK(0.0083); |
| GYNVNDEKA | GYNVNDEK[1 | K | K | 11 | 24 | 14 | 3 | 8K(114.0429) | sp\|P4452_BA | P4452_BASP | 0 | 1 | 0 | 2,39E+07 | unmatched | MS/MS | | K:114.0429@GYNVNDEK(0.9334)AK(0.0404)EK(0.0161)DK(0.0101); |
| KAEGAATEEE | KAEGAATEEE | K | E | 25 | 38 | 14 | 2,3 | | sp\|P4452_BA | P4452_BASP | 4 | 7 | 0 | 2273951 | MS/MS | MS/MS | | |
| KAEGAATEEE | KAEGAATEEE | K | E | 25 | 52 | 28 | 2,3,4 | | sp\|P4452_BA | P4452_BASP | 15 | 13 | 1,49E+09 | 7,92E+09 | MS/MS | MS/MS | | |
| KAEGAATEEE | KAEGAATEEE | K | E | 25 | 55 | 31 | 2,3,4,5 | | sp\|P4452_BA | P4452_BASP | 12 | 17 | 5,71E+07 | 6,08E+09 | MS/MS | MS/MS | | |
| KAEGAATEEE | KAEGAATEEE | K | E | 25 | 52 | 28 | 4 | 14K(114.0429) | sp\|P4452_BA | P4452_BASP | 0 | 2 | 0 | 3793984 | unmatched | MS/MS | | K:114.0429@K(0.0098)AEGAATEEEGTPK(0.9818)ESEPQAAAEPAEAK(0.0084); |
| KGYNVNDEK | KGYNVNDEK | K | A | 10 | 18 | 9 | 2 | | sp\|P4452_BA | P4452_BASP | 2 | 2 | 7482745,5 | 1,76E+07 | MS/MS | MS/MS | | |
| KGYNVNDEK | KGYNVNDEK | K | E | 10 | 20 | 11 | 2,3 | | sp\|P4452_BA | P4452_BASP | 15 | 11 | 7,83E+08 | 2,19E+09 | MS/MS | MS/MS | | |
| KGYNVNDEK | KGYNVNDEK | K | D | 10 | 22 | 13 | 2,3,4 | | sp\|P4452_BA | P4452_BASP | 23 | 22 | 3,06E+08 | 8,83E+08 | MS/MS | MS/MS | | |
| KKGYNVNDE | KKGYNVNDE | K | A | 9 | 18 | 10 | 3 | | sp\|P4452_BA | P4452_BASP | 0 | 1 | 0 | 4877586,5 | unmatched | MS/MS | | |
| KKGYNVNDE | KKGYNVNDE | K | E | 9 | 20 | 12 | 2,3 | | sp\|P4452_BA | P4452_BASP | 14 | 17 | 8531730 | 1,92E+07 | MS/MS | MS/MS | | |
| KKKGYNVND | KKKGYNVND | K | A | 8 | 18 | 11 | 3 | | sp\|P4452_BA | P4452_BASP | 27 | 0 | 1,10E+07 | 0 | MS/MS | unmatched | | |
| KKKGYNVND | KKK[114.042 | K | A | 8 | 18 | 11 | 3 | 3K(114.0429) | sp\|P4452_BA | P4452_BASP | 0 | 2 | 0 | 6750669,5 | unmatched | MS/MS | | K:114.0429@K(0.0165)K(0.0399)K(0.9380)GYNVNDEK(0.0055); |
| KKGYNVNDE | KK[114.0429] | K | A | 9 | 18 | 10 | 2 | 2K(114.0429) | sp\|P4452_BA | P4452_BASP | 0 | 1 | 0 | 1,99E+07 | unmatched | MS/MS | | K:114.0429@K(0.1553)K(0.8276)GYNVNDEK(0.0171); |
| KKKGYNVND | KK[114.0429] | K | A | 8 | 18 | 11 | 3 | 2K(114.0429) | sp\|P4452_BA | P4452_BASP | 0 | 2 | 0 | 8919806 | unmatched | MS/MS | | K:114.0429@K(0.0252)K(0.4852)K(0.4852)GYNVNDEK(0.0044); |
| KTEAPAAPAA( | KTEAPAAPAA( | K | S | 150 | 163 | 14 | 2,3 | | sp\|P4452_BA | P4452_BASP | 10 | 17 | 2,11E+09 | 2,00E+10 | MS/MS | MS/MS | | |
| KTEAPAAPAA( | KTEAPAAPAA( | K | E | 150 | 184 | 35 | 3,4,5 | | sp\|P4452_BA | P4452_BASP | 11 | 47 | 3,60E+09 | 2,57E+10 | MS/MS | MS/MS | | |
| KTEAPAAPAA( | KTEAPAAPAA( | K | A | 150 | 198 | 49 | 3,4,5,6 | | sp\|P4452_BA | P4452_BASP | 22 | 15 | 7,75E+09 | 7,02E+09 | MS/MS | MS/MS | | |
| KTEAPAAPAA( | KTEAPAAPAA( | K | E | 150 | 184 | 35 | 5 | 24K(114.0429),35K | sp\|P4452_BA | P4452_BASP | 1 | 0 | 3654815 | 0 | MS/MS | unmatched | K:114.0429@K(0.0158)TEAPAAPAAQETK(0.0173)SDGAPASDSK(0.9834)PGSSEAAPSSK(0.9834); | |
| KAEGAATEEE | K[114.0429]A | K | E | 25 | 38 | 14 | 2 | 1K(114.0429) | sp\|P4452_BA | P4452_BASP | 0 | 1 | 0 | 0 | unmatched | MS/MS | | K:114.0429@K(0.9813)AEGAATEEEGTPK(0.0187); |
| KAEGAATEEE | K[114.0429]A | K | E | 25 | 52 | 28 | 4 | 1K(114.0429) | sp\|P4452_BA | P4452_BASP | 0 | 1 | 0 | 8743760 | unmatched | MS/MS | | K:114.0429@K(0.9514)AEGAATEEEGTPK(0.0329)ESEPQAAAEPAEAK(0.0158); |
| KAEGAATEEE | K[114.0429]A | K | E | 25 | 52 | 28 | 3,4,5 | 14K(114.0429),1K( | sp\|P4452_BA | P4452_BASP | 1 | 5 | 0 | 0 | MS/MS | MS/MS | K:114.0429@ | K:114.0429@K(0.9817)AEGAATEEEGTPK(0.9817)ESEPQAAAEPAEAK(0.0366); |
| KAEGAATEEE | K[114.0429]A | K | E | 25 | 55 | 31 | 5 | 14K(114.0429),1K( | sp\|P4452_BA | P4452_BASP | 2 | 0 | 0 | 0 | MS/MS | unmatched | K:114.0429@K(0.9784)AEGAATEEEGTPK(0.9783)ESEPQAAAEPAEAK(0.0197)EGK(0.0236); | |
| KGYNVNDEK | K[114.0429]G | K | A | 10 | 18 | 9 | 2,3 | 1K(114.0429) | sp\|P4452_BA | P4452_BASP | 0 | 6 | 0 | 8,85E+08 | unmatched | MS/MS | | K:114.0429@K(0.9701)GYNVNDEK(0.0299); |
| KGYNVNDEK | K[114.0429]G | K | E | 10 | 20 | 11 | 2,3 | 1K(114.0429) | sp\|P4452_BA | P4452_BASP | 0 | 5 | 0 | 4,21E+08 | unmatched | MS/MS | | K:114.0429@K(0.9703)GYNVNDEK(0.0167)AK(0.0130); |
| KKGYNVNDE | K[114.0429]K | K | A | 9 | 18 | 10 | 3 | 1K(114.0429) | sp\|P4452_BA | P4452_BASP | 0 | 1 | 0 | 4,58E+07 | unmatched | MS/MS | | K:114.0429@K(0.9665)K(0.0107)GYNVNDEK(0.0229); |
| KKGYNVNDE | K[114.0429]K | K | E | 9 | 20 | 12 | 3 | 1K(114.0429) | sp\|P4452_BA | P4452_BASP | 0 | 2 | 0 | 2638355,8 | unmatched | MS/MS | | K:114.0429@K(0.8554)K(0.0607)GYNVNDEK(0.0522)AK(0.0316); |
| KKKGYNVND | K[114.0429]K | K | A | 8 | 18 | 11 | 3 | 1K(114.0429) | sp\|P4452_BA | P4452_BASP | 0 | 3 | 0 | 2136502,5 | unmatched | MS/MS | | K:114.0429@K(0.4720)K(0.4720)K(0.0514)GYNVNDEK(0.0046); |
| SDGAPASDSI | SDGAPASDSI | K | E | 164 | 184 | 21 | 2,3 | | sp\|P4452_BA | P4452_BASP | 4 | 16 | 1,65E+09 | 1,68E+10 | MS/MS | MS/MS | | |
| SDGAPASDSI | SDGAPASDSI | K | A | 164 | 198 | 35 | 2,3,4 | | sp\|P4452_BA | P4452_BASP | 9 | 13 | 4,29E+09 | 7,02E+09 | MS/MS | MS/MS | | |
| SDGAPASDSI | SDGAPASDSI | K | A | 164 | 198 | 35 | 4 | 21K(114.0429) | sp\|P4452_BA | P4452_BASP | 0 | 1 | 0 | 4578969 | unmatched | MS/MS | | K:114.0429@SDGAPASDSK(0.3756)PGSSEAAPSSK(0.6182)ETPAATEAPSSTPK(0.0061); |
| SDGAPASDSI | SDGAPASDSI | K | E | 164 | 184 | 21 | 2,3 | 10K(114.0429) | sp\|P4452_BA | P4452_BASP | 0 | 2 | 0 | 5634780 | unmatched | MS/MS | | K:114.0429@SDGAPASDSK(0.9829)PGSSEAAPSSK(0.0171); |
| TEAPAAPAAQ | TEAPAAPAAQ | K | S | 151 | 163 | 13 | 2 | | sp\|P4452_BA | P4452_BASP | 3 | 3 | 7,70E+07 | 2,99E+08 | MS/MS | MS/MS | | |
| TEAPAAPAAQ | TEAPAAPAAQ | K | E | 151 | 184 | 34 | 3,4 | | sp\|P4452_BA | P4452_BASP | 5 | 6 | 4,59E+07 | 5,19E+08 | MS/MS | MS/MS | | |
| TEAPAAPAAQ | TEAPAAPAAQ | K | A | 151 | 198 | 48 | 4,5,6 | | sp\|P4452_BA | P4452_BASP | 5 | 5 | 1,57E+08 | 8,25E+07 | MS/MS | MS/MS | | |
| TEAPAAPAAQ | TEAPAAPAAQ | K | A | 151 | 198 | 48 | 5 | 23K(114.0429) | sp\|P4452_BA | P4452_BASP | 1 | 0 | 0 | 0 | MS/MS | unmatched | K:114.0429@TEAPAAPAAQETK(0.0080)SDGAPASDSK(0.9719)PGSSEAAPSSK(0.0140)ETPAATEAPSSTPK(0.0061); | |
| TEAPAAPAAQ | TEAPAAPAAQ | K | A | 151 | 198 | 48 | 4,5 | 23K(114.0429),34K | sp\|P4452_BA | P4452_BASP | 0 | 5 | 0 | 8973840 | unmatched | MS/MS | | K:114.0429@TEAPAAPAAQETK(0.0198)SDGAPASDSK(0.9773)PGSSEAAPSSK(0.9773)ETPAATEAPSSTPK(0.0256 |
| TEAPAAPAAQ | TEAPAAPAAQ | K | E | 151 | 184 | 34 | 5 | 13K(114.0429) | sp\|P4452_BA | P4452_BASP | 0 | 1 | 0 | 0 | unmatched | MS/MS | | K:114.0429@TEAPAAPAAQETK(0.9374)SDGAPASDSK(0.0431)PGSSEAAPSSK(0.0195); |
| TEGAAEAKAE | TEGAAEAKAE | K | A | 90 | 121 | 32 | 3,4 | | sp\|P4452_BA | P4452_BASP | 9 | 6 | 7,24E+07 | 3,48E+08 | MS/MS | MS/MS | | |

**Supplementary table 11:** Proteomics of ubiquitin-modified BASP1 peptides. BASP1 gel bands obtained following the *in vitro* ubiquitination assay were excised, subjected to tryptic digestion, and analyzed by mass spectrometry (MS). The table lists the identified ubiquitin-modified peptides and their assigned modification sites. Related to Supplementary Figure 4I.

| Praja1/MHD fusion protein | |
|---|---|
| **Data collection** | |
| Space group | P42212 |
| Cell dimensions | |
| a, b, c | 95.40, 95.40, 70.43 |
| a, b, g (°) | 90, 90, 90 |
| Resolution (Å) | 56.66 – 1.922 |
| $R_{merge}$ | 0.124 |
| $R_{meas}$ | 0.131 |
| $R_{pim}$ | 0.041 |
| $CC_{1/2}$ | 0.999 |
| I / sI | 11.7 |
| Completeness (%) | 99.9 |
| Redundancy | 9.9 |
| **Refinement** | |
| Resolution (Å) | 1.922 |
| No. reflections | 207119 |
| $R_{work}$ | 0.2537 |
| $R_{free}$ | 0.2866 |
| No. atoms | |
| Protein | 4345 |
| Water | 430 |
| Wilsons B-factor/Average B-factor | 28.61/40.64 |
| R.m.s. deviations | |
| Bond lengths (Å) | 0.0088 |
| Bond angles (°) | 2.0466 |
| Ramachandran | |
| favored (%) | 96.4 |
| allowed (%) | 3.6 |
| outliers (%) | 0.00 |

**Supplementary table 12:** X-ray data collection and refinement statistics for the Praja1/MHD fusion protein
